# Interactive effects of salinity, temperature and food web configuration on performance and harmfulness of the raphidophyte *Heterosigma akashiwo*

**DOI:** 10.1101/2023.06.23.546213

**Authors:** Jakob Karl Giesler, Daniel Alan Lemley, Janine Barbara Adams, Stefanie Devi Moorthi

## Abstract

The cosmopolitan raphidophyte *Heterosigma akashiwo* commonly forms harmful algal blooms (HABs) in diverse estuaries discharging into Algoa Bay, South Africa, potentially leading to hypoxia, fish kills and a decline in key primary consumers. Despite the high environmental variability in these estuaries, little is known about how abiotic factors such as temperature and salinity constrain bloom formation and harmfulness of *H. akashiwo*. The present study therefore investigates growth, competition, and grazing interactions of H. akashiwo in laboratory experiments in response to two naturally relevant levels of salinity (15, 30) and temperature (16, 22°C), respectively. Experiments were set up with the naturally co-occurring dinoflagellate competitor *Heterocapsa rotundata* and two estuarine microzooplankton consumers, i.e., nauplii of the copepod *Acartia tonsa* and the rotifer *Brachionus plicatilis*. In monoculture, *H. akashiwo* growth was promoted at high temperature – low salinity conditions, while *H. rotundata* thrived under low temperature – high salinity conditions. In polyculture, *H. akashiwo* dominated at high temperature irrespective of the salinity regime, while at low temperature, it only dominated at low salinity and was suppressed by *H. rotundata* at high salinity. Grazing assays revealed highly negative effects of *H. akashiwo* on copepod nauplii survival and growth as well as mucus-induced immobilization, especially at high temperatures in combination with low salinity, while the estuarine adapted rotifers showed highest mortalities at the higher salinity level. The presence of *H. rotundata* significantly alleviated the harmful effects of *H. akashiwo* on both grazers, and the selectively feeding copepod nauplii actively avoided *H. akashiwo* when non-harmful prey was present. Overall, this study demonstrates that population dynamics and harmful effects of *H. akashiwo* are interactively determined by both abiotic conditions and food web configuration, implying competitor and consumer specific tolerances to the abiotic environment and their susceptibility to the harmful alga *H. akashiwo*.

## 2 Introduction

*Heterosigma akashiwo* is a harmful raphidophyte, occurring in various coastal and estuarine waters around the world. Numerous bloom events have been reported for temperate regions including Canada (Taylor and Haigh, 1993), the United States (Kempton et al., 2007), Japan (Li and Smayda, 2000) and China (Wang et al., 2007), as well as for subtropical waters of Brazil (Branco et al., 2014) and multiple South African estuaries (Lemley et al., 2021, Lemley et al., 2018b). The cosmopolitan distribution of *H. akashiwo* reflects its broad ecological tolerance to wide ranges of temperature, salinity, and light intensity (Allaf and Trick, 2019, Strom et al., 2013, Martinez et al., 2010). As such, this species is well-adapted to the highly variable and potentially stressful temperature and salinity regimes of estuarine ecosystems, such as the Sundays and Hartenbos estuaries in South Africa. These microtidal estuarine systems are characterized by highly variable abiotic conditions (Lemley et al., 2018b), mouth conditions (Adams et al., 2020), and levels of anthropogenic pressures (e.g., flow modification, nutrient loading) (Lemley et al., 2021, Dalu et al., 2018). Both estuaries experience aggravated nutrient pollution, however, the primary source of anthropogenic nutrients varies between the Sundays (agricultural practices) and Hartenbos (treated wastewater discharges) estuaries. Yet, the augmented nature of freshwater inflow regimes and continuous supply of nutrient-rich baseflows make both systems particularly prone to harmful algae bloom (HAB) events by *H. akashiwo* (Lemley et al., 2018c). This species dominates the phytoplankton community in the Sundays Estuary mainly during spring/summer when elevated water temperatures and stratified mesohaline waters facilitate nearly monospecific bloom events (>550 µg Chl a L^−1^) (Lemley et al. 2018b). Such HAB events have been shown to severely impact the ecosystem by, for example, inducing bottom-water hypoxia (Lemley et al., 2018b), reduced abundance of key primary consumers (Smit et al., 2021) as well as deleterious effects on mugilid fish gills (Bornman et al., 2022) and embryonic fish cells (Sandoval-Sanhueza et al., 2022). While for most HAB-forming species the predominating mode of harmfulness is well understood, different harmful mechanisms have been proposed for *H. akashiwo*. They include the production of neurotoxins (Khan et al., 1997), cytotoxins (Sandoval-Sanhueza et al., 2022), allelochemicals (Yamasaki et al., 2009, Yamasaki et al., 2007, Qiu et al., 2012), reactive oxygen species (Yang et al., 1995, Twiner and Trick, 2000, Oda et al., 1997), and adhesive mucus, with the latter potentially leading to gill clogging of fish and blocking of the zooplankton filtration apparatus (Nakamura et al., 1998, Lemley et al., 2018b). Yet, there is no scientific consensus about which of these mechanisms is the predominant harmful mode in the raphidophyte.

Another bloom-forming species in the Sundays Estuary is the dinoflagellate *Heterocapsa rotundata*, which represents a dominant competitor of *H. akashiwo* (Adams et al., 2020). Although *H. rotundata* is well documented as red tide-forming species, it is not recognized as causing harmful effects (Iwataki, 2008). Field data indicate that the two species proliferate at contrary environmental conditions in terms of temperature and, locally, also in terms of salinity, with *H. akashiwo* generally preferring increased water temperatures and *H. rotundata* thriving at lower temperatures (Rothenberger et al., 2009, Lemley et al., 2018b). Therefore, the variability of these abiotic conditions can be considered an important driving force for the bloom dynamics of these two species, consequently also affecting herbivorous grazer populations and higher trophic levels (Burkholder et al., 2018). While Lemley et al. (2018a, 2020) demonstrated the in-situ limiting effect of *H. akashiwo* blooms on co-occurring plankton taxa in the Sundays Estuary, Qiu et al. (2012) investigated competitive interactions between *H. akashiwo* and other phytoplankton species in controlled laboratory experiments. The outcome revealed a strong inhibitory effect of a Japanese strain of *H. akashiwo* on the dinoflagellate *Akashiwo sanguinea* caused by allelochemical polysaccharide-protein complexes that even led to cell deformation in the competitor. Other studies focused on the harmful effects of *H. akashiwo* on primary consumers and provided evidence for adverse effects of this HAB species on in-situ larval fish and zooplankton densities (Smit et al., 2021), on survival, grazing, and recruitment of the copepod *Schmackeria inopinus* (Yu et al., 2010), as well as on population growth of the rotifer *Brachionus plicatilis* (Yan et al., 2009, Xie et al., 2007). However, neither the competition studies nor the grazing assays conducted so far took into account how competitive outcome and the coupled grazer proliferation could be altered by different abiotic environmental conditions. Yet, interactive effects of abiotic and biotic determinants are crucial to be considered in order to fully understand the mechanisms driving bloom dynamics of harmful species and to enhance our predictive understanding of such blooms (Purz et al., 2021).

The present study therefore investigates the bloom dynamics and associated harmfulness of a South African *H. akashiwo* strain in response to altered temperature and salinity regimes in monoculture, as well as in a food web context including a dinoflagellate competitor and two different herbivorous consumers. A full-factorial competition experiment was conducted at two naturally relevant levels of salinity and temperature, respectively, to identify growth preferences of *H. akashiwo* in monoculture and its competitive ability in polyculture under different environmental conditions. In subsequent grazing assays, two different consumers were added to the system (separately) under the same temperature and salinity regimes to test for consumer mortality, grazing rates, and consumer-specific sensitivity to the harmful algae. For all experiments, only species were used that are known to co-occur with *H. akashiwo* in brackish estuarine waters, i.e., the dinoflagellate *H. rotundata* as competitor, the non-selectively feeding rotifer *Brachionus plicatilis* and nauplii of the selectively feeding copepod *Acartia tonsa* as grazers.

Based on the patterns previously observed in the field and in small-scale laboratory experiments, temperature and salinity variations were expected to differentially select for the dominance of the two phytoplankton species in polyculture. In both monoculture and polyculture experimental setups, *H. akashiwo* was expected to be promoted at high temperature – low salinity conditions and *H. rotundata* at low temperature – high salinity conditions. Furthermore, herbivorous consumers were predicted to experience negative effects in terms of survival, grazing, and developmental stages in the presence of *H. akashiwo.* The strongest harmful effects were expected to be found under optimum growth conditions of the harmful alga (i.e., at high temperature – low salinity conditions). The presence of non-harmful prey was hypothesized to promote consumer growth in monoculture and alleviate the harmful effects of *H. akashiwo* in mixed prey cultures.

## 3 Material and Methods

### 3.1 Experimental Cultures and Culture Conditions

The raphidophyte *Heterosigma akashiwo* was isolated from a water sample taken from the Hartenbos Estuary along the warm-temperate south coast of South Africa in 2019, and the dinoflagellate *Heterocapsa rotundata* was obtained from the Roscoff Culture Collection, France (strain number RCC3042). Both algae were cultivated in f/2 medium (Guillard, 1975) prepared from 0.2 μm filtered and sterilized seawater with a salinity of 30, taken from the Jade Bay in Wilhelmshaven, Germany. By dilution with f/2 medium, the cultures were kept in exponential growth phase and were cultivated under controlled conditions in a climate chamber at 18°C, 70 μmol photons m^-2^ s^-1^ and a 12:12 h light:dark period. Eggs of the copepod *Acartia tonsa* were obtained from the AWI Biological Institute Helgoland (BAH), Germany, and stored at 6°C in total darkness until use. The rotifer *Brachionus plicatilis* was obtained from the aquatic retailer Interaquaristik, Germany, and was grown under the same culture conditions as described for the phytoplankton species. Every two weeks, the *Brachionus* culture was transferred into new medium and was fed with the green algae *Tetraselmis* sp. (also derived from the Roscoff Culture Collection, France).

### 3.2 Competition Experiment

A competition experiment was conducted to investigate the growth and competitive ability of *H. akashiwo* in the presence of the dinoflagellate *H. rotundata* under variable estuarine conditions by establishing two levels of temperature (16 and 22°C) in combination with two levels of salinity (15 and 30) in a full factorial design. Temperature and salinity levels were selected based on the findings of Lemley et al. (2018b) in the Sundays Estuary, South Africa, and thus represent typical ranges for the naturally occurring variability of temperature and salinity in South African estuaries. Both algae were inoculated at equal biovolume proportions to account for the size difference of the two species. For that purpose, the dimensions of 30 cells of each species were measured using a Zeiss Axiovert 10 inverted microscope and the individual cell biovolume of *H. akashiwo* and *H. rotundata* was calculated according to Hillebrand et al. (1999). Both species were inoculated at a total biovolume of 2.43 × 10^6^ μm^3^, corresponding to an initial cell density of 5 × 10^3^ cells ml^-1^ for *H. akashiwo* and 21 × 10^3^ cells ml^-1^ for *H. rotundata* (cell volume *H. akashiwo*: 487 μm^3^, cell volume *H. rotundata*: 114 μm^3^). Both species were set up in monoculture and polyculture, using an additive design, i.e., inoculating each species with the same biovolume in monoculture and polyculture, resulting in twice the total biovolume in the polycultures compared to monocultures. This additive design was chosen in consideration of the subsequent grazing assays, where the consumers were incubated with the same algal monocultures and polycultures. This assured equal amounts of the harmful algae (i.e., equal concentrations of potentially harmful substances) in all treatments. Each treatment was replicated by four, resulting in a total of 48 experimental units (3 species combinations x 2 salinity levels x 2 temperature levels x 4 replicates), containing 200 ml culture in a sterile 250 ml Erlenmeyer flask. To establish the lower salinity level of 15, f/2 medium was diluted with fully desalinated water enriched with f/2 nutrient concentrations and the salinity was checked with a Multi 3630 IDS SET KS2 sensor. The experimental units were incubated, using the empty containers of an indoor mesocosm facility (Planktotrons, (Gall et al., 2017) at their respective experimental temperature (16°C and 22°C) and a light intensity of 100 μmol photons m^-2^ s^-1^ with a light:dark period of 12:12 hours. Sampling was conducted every 48 hours and comprised subsamples of 500 μl from each experimental unit. Samples were fixed in a 48-well microplate (83.3923 Sarstedt AG & Co KG ©) with formalin at a final concentration of 2% for quantification conducted with a Leica DMIL inverted bright field microscope at a magnification of 100x. Every fourth day, an additional 10 ml of each experimental unit were sampled for subsequent dissolved macronutrient analysis. Furthermore, at the end of the experiment, 10 ml samples were taken from each experimental unit for particulate macronutrient analysis. The experiment was terminated after 28 days, when the last treatment had reached the stationary growth phase.

### 3.3 Copepod grazing assay

A grazing assay was conducted to evaluate whether and how the potential harmful effects of *H. akashiwo* on copepod nauplii are altered by different temperature and salinity regimes, as well as by the presence of a non-harmful competitor. Prior to the main experiment, a pilot study was performed in which the effect of a *H. akashiwo* culture (4 x 10^4^ cells ml^-1^) on *A. tonsa* nauplii (1 individual ml^-1^) survival was compared to a starvation control (sterile filtered seawater without algal food) in a five-day incubation. This was done to test whether the harmful effects of *H. akashiwo* on copepod nauplii exceed a pure starvation effect, i.e., whether the potential mucus production of the harmful algae mainly hinders filter feeding in the consumer, leading to starvation, or whether the harmful effects are stronger, indicating the production of allelochemicals or other harmful substances.

For the subsequent full factorial grazing assay, the same experimental salinity and temperature levels and the same algae species combinations were used as in the competition experiment *(Section 3.2)*, but with the addition of *A. tonsa* nauplii at an initial density of 1 individual ml^-1^. To provide sufficient prey for the nauplii, *H. akashiwo* was inoculated with an initial cell density of 1.5 × 10^4^ cells ml^-1^, corresponding to 6.4 × 10^4^ cells ml^-1^ *H. rotundata*, resulting in equal biovolume proportions of each phytoplankton species in monoculture and polyculture. Phytoplankton were inoculated again using the additive design, resulting in twice the total algal biovolume in polyculture to ensure that the grazers were exposed to the same biovolume of *H. akashiwo* and thus the same amount of potentially harmful substances in all treatments (see above). Due to space constraints, no additional non-grazer control treatments containing only the phytoplankton species were set up; instead, the competition experiment, which was conducted only a few weeks before, was used as non-grazer control.

After hatching, nauplii stock cultures were prepared according to Meunier et al. (2016). Experimental units were set up with growth medium and phytoplankton / consumer stock cultures in a total volume of 200 ml contained in 250 ml Erlenmeyer flasks. The preparation of the experimental units in terms of salinity was identical to the procedure described for the competition experiment. After incubation in the Planktotron containers, phytoplankton sampling was conducted every 24 hours with the identical procedure as described for the competition experiment. For evaluation of zooplankton mortality, additional 10 ml subsamples were taken daily from each experimental unit and transferred into 6-well microplates for quantification with an inverted light microscope. The mortality was then calculated by dividing the number of dead individuals in the live samples by the total number of nauplii counted in each sample after fixation with formalin at 2% final concentration. An individual was considered dead after no physical movement for at least 60 seconds was observed. The experiment was terminated after five days, when the first treatment showed a mortality of 100%. At the end of the experiment, the developmental stages of the nauplii in different treatments were evaluated by micrographs of three individuals within each replicate taken with a Jenoptik Gryphax Kapella camera connected to a Zeiss Axiovert 10 microscope. At the end of the experiment, additional 10 ml subsamples were taken from each experimental unit for dissolved and particulate macronutrient analysis, respectively. For the particulate macronutrient subsamples, grazers were removed by filtration through a 20 µm gaze.

### 3.4 Rotifer grazing assay

A second grazing assay was conducted with the rotifer *Brachionus plicatilis* to test for potential differences in grazer sensitivity to the harmful algae. Since the results of the prior competition experiment (*Section 3.2*) indicated that salinity effects depended on temperature, the rotifer bioassay was reduced to the two salinity levels (15 and 30) at a fixed experimental temperature of 16°C. This reduced experimental setup allowed for the addition of non-grazer control treatments containing only the phytoplankton species in monoculture and polyculture. All treatments were replicated three times.

Prior to the start of the experiment, the rotifers were starved for 24 hours by careful rinsing with sterile seawater on a 50 μm gaze mesh to remove any prey items, followed by resuspension in 500 ml beakers. After the starvation period, these cultures were used as stocks for the experiment. Next, the experimental units, with a total volume of 150 ml each, were inoculated with the rotifers at a density of 1 individual ml^-1^. The preparation of the experimental units in terms of salinity, as well as the initial cell densities of the two phytoplankton species, was identical to the copepod nauplii bioassay (*Section 3.3*). After incubation in the Planktotron containers under the same controlled conditions at 16°C, grazer and phytoplankton sampling for microscopy were conducted every third day (including day 0) with identical procedures as previously described in *Section 2.2* and *2.3*, respectively. The experiment was terminated after six days.

### 3.5 Analysis of particulate and dissolved macronutrients

Samples for dissolved inorganic nutrients (nitrogen, phosphorus, silicate) were prefiltered through 0.2 µm syringe filters (Sarstedt Filtropour S) and stored in PE bottles at -20°C until analysis using a SAN++ Continuous Flow Analyzer (CFA, Skalar Analytical B.V., Netherlands). For particulate intracellular carbon (C), nitrogen (N), and phosphorus (P) analyses, two 10 ml subsamples were filtered on pre-combusted and acid washed Whatman GF/F filters for CN and P analyses, respectively. The CN elemental composition was measured using a CHN analyzer (Thermo, Flash EA 1112). Particulate phosphate was measured as orthophosphate by molybdate reaction after sulfuric acid digestion according to Grasshoff et al. (2009).

### 3.6 Data Analyses

To test for optimum growth conditions of the two phytoplankton species, as well as for the impact of the competitor in the competition experiment (*Section 3.2*), treatment effects on phytoplankton biovolume were tested for each species separately, using linear mixed models (LMMs) with temperature, salinity, presence of the competitor, and time as fixed factors, as well as the index number of the experimental unit representing the random factor. The LMMs were conducted using the “lmer” function from the “lmerTest” package (Kuznetsova et al., 2017) that includes the Satterthwaite approximation for obtaining p-values. To meet the model assumptions of homoscedasticity, algal biovolumes were square root transformed prior to the analysis. For testing dominance shifts over time in the polycultures, log ratios of the two phytoplankton species biovolumes were calculated for each sampling point which were then analysed separately for each factorial treatment combination against time using Kruskal-Wallis tests. Subsequently, non-parametric pairwise Wilcoxon rank sum post hoc tests were used to compare the log ratios of every sampling day to the log ratio of day 0 at which both species were inoculated at equal biovolume proportions. Moreover, non-linear models were fitted using the “nls” base R function to obtain the maximum growth rates and carrying capacities of the two phytoplankton species in each treatment which were then tested separately for both species using two-way ANOVAs with salinity and temperature as independent variables. To test for differences in phytoplankton particulate molar C:N ratios at the end of the competition experiment and nauplii grazing assay, a three-way ANOVA was performed with species composition, temperature, and salinity as independent variables. Model assumptions of homogeneous variances and normally distributed residuals were tested with Fligner-Killeen tests and Shapiro-Wilk tests, respectively. Due to technical problems with the particulate phosphorous analysis, P-concentrations could not be considered here and no C:P or N:P ratios could be calculated.

In the nauplii and rotifer grazing assays, effects on mortality were analyzed using LMMs with temperature, salinity, prey composition, and time as fixed factors, as well as the index of the experimental unit as random factor for the nauplii grazing assay. The same factors, except for temperature (i.e., experiment was conducted at only one temperature level) were used for rotifers. To calculate grazing rates, phytoplankton growth rates in the absence of consumers (gross growth rate = μ) and with consumers (net growth rate = r) were calculated between day 0 and day 5 by subtracting ln-transformed starting biovolume from ln-transformed final biovolume and dividing this by the respective number of experimental days (five days for *A. tonsa* nauplii, six days for *B. plicatilis*). The net population grazing rate (g) was then calculated as μ − r. For the prey mixture treatments that contained both phytoplankton species, biovolumes were pooled prior to the grazing rate calculations. As previously described, there were no additional non-grazer controls for the copepod grazing assay so that the competition experiment was used as such. Consequently, the resulting grazing rates require cautious interpretation as the two experiments were set up with different initial biovolumes. However, no density-dependent effects on phytoplankton growth could be expected in any of the experiments during the first five days of the experiments (duration of the nauplii grazing assay). Moreover, the experiments were conducted in the same mesocosm containers with only a short time interval in between and represented identical conditions, allowing to compare phytoplankton growth rates in the different setups. To test for differences in relative population grazing rates of the nauplii in different temperature and salinity treatments, generalized least squares (GLS) models were conducted, to compensate for groups with non-homogeneous variances by using weights. Effects on rotifer relative population grazing rates were tested using three-way ANOVAs with temperature, salinity, and prey composition as independent variables. Moreover, to test for selective grazing effects in the mixed prey cultures, separate grazing rates were calculated for both phytoplankton species which were then tested in a three-way ANOVA with temperature, salinity, and phytoplankton species identity for the nauplii bioassay, and a two-way ANOVA with salinity and phytoplankton species identity for the rotifer bioassay. For comparing the sensitivity to harmful effects of *A. tonsa* and *B. plicatilis*, another LMM was performed on grazer mortality as dependent variable and salinity, prey composition, grazer identity, and time as independent variables. However, from the nauplii grazing assay, only data from the lower temperature treatments were considered here, since the rotifer grazing assay was conducted only at 16°C. Differences in population grazing rates between the two grazer species were tested using a three-way ANOVA with the same set of independent variables (except for time).

Post-hoc analyses were conducted with estimated marginal means pairwise comparisons using the “emmeans” function from the “emmeans” package (Lenth, 2021) for GLS models and Tukey post-hoc tests for LMMs and ANOVAs, respectively. All statistical analyses and graphs of this study were performed using the R statistical environment 4.0.2. (R Core Team, 2020).

## 4 Results

### 4.1 Competition Experiment

*H. akashiwo* and *H. rotundata* biovolumes were both significantly and interactively affected by salinity, temperature, time, and the presence of the competitor, except for *H. rotundata* biovolumes that did not show a significant effect by its competitor (Table 1).

**Table 1:** Results of linear mixed models testing the effects of salinity, temperature, competitor and day (fixed effects) on the square rooted biovolumes of *H. akashiwo* and *H. rotundata* (monocultures and polyculture treatments together) with beaker number (id) as random factor. Degrees of freedom (df), F and p-values are given for each effect. Values marked with an asterisk (*) indicate significant effects (p < 0.05).

| Fixed Effect | Df | *H. akashiwo*<br>sqrt(biovolume) |  | *H. rotundata*<br>sqrt(biovolume) |  |
| --- | --- | --- | --- | --- | --- |
|  |  | *F* | *p* | *F* | *p* |
| Salinity | 1 | 355.0 | <0.001 * | 5715.3 | <0.001 * |
| Temp. | 1 | 347.5 | <0.001 * | 1807.2 | <0.001 * |
| Competitor | 1 | 470.2 | <0.001 * | 3.3 | 0.08 |
| Day | 14 | 2961.8 | <0.001 * | 812.6 | <0.001 * |
| Salinity × Temp. | 1 | 21.5 | <0.001 * | 326.0 | <0.001 * |
| Salinity × Competitor | 1 | 171.0 | <0.001 * | 26.0 | <0.001 * |
| Temp. × Competitor | 1 | 226.5 | <0.001 * | 58.4 | <0.001 * |
| Salinity × Day | 14 | 49.7 | <0.001 * | 342.4 | <0.001 * |
| Temp. × Day | 14 | 34.0 | <0.001 * | 65.8 | <0.001 * |
| Competitor × Day | 14 | 61.1 | <0.001 * | 6.1 | <0.001 * |
| Salinity × Temp. × Competitor | 1 | 27.4 | <0.001 * | 16.3 | <0.001 * |
| Salinity × Temp. × Day | 14 | 31.9 | <0.001 * | 113.3 | <0.001 * |
| Salinity × Competitor × Day | 14 | 25.7 | <0.001 * | 18.4 | <0.001 * |
| Temp. × Competitor × Day | 14 | 64.5 | <0.001 * | 15.2 | <0.001 * |
| Salinity × Temp. × Competitor × Day | 14 | 20.0 | <0.001 * | 11.2 | <0.001 * |
| **Random Effect** |  |  |  |  |  |
| $\sigma^2$ | | 209087.9 | | 77523.5 | |
| $\tau_{00 \text{ id}}$ | | 58417.7 | | 5474.0 | |
| ICC |  | 0.22 |  | 0.07 |  |
| $N_{\text{id}}$ | | 32 | | 32 | |
| Observations |  | 480 |  | 480 |  |
| Marginal $R^2$ / Conditional $R^2$ | | 0.989 / 0.991 | | 0.986 / 0.987 | |

In monoculture, *H. akashiwo* biovolume was significantly higher at the higher experimental temperature and lower salinity level (Fig. 1a, 1d). Low salinity treatments resulted in higher carrying capacities for *H. akashiwo* (p < 0.001, Table 2). The positive temperature effect was reflected in higher maximum growth rates for *H. akashiwo* at 22°C compared to 16°C (Table S1), being highest (0.53 day^-1^) in the low salinity – high temperature treatment, differing from all other treatment combinations (p < 0.004). Depending on the respective treatment, *H. akashiwo* reached its maximum biovolume in monoculture between day 18 and 24, with the 16°C treatments generally achieving a time-delayed, but notably higher carrying capacity than at 22°C (Fig. 1a, 1d, Table S2).

**Figure 1:**
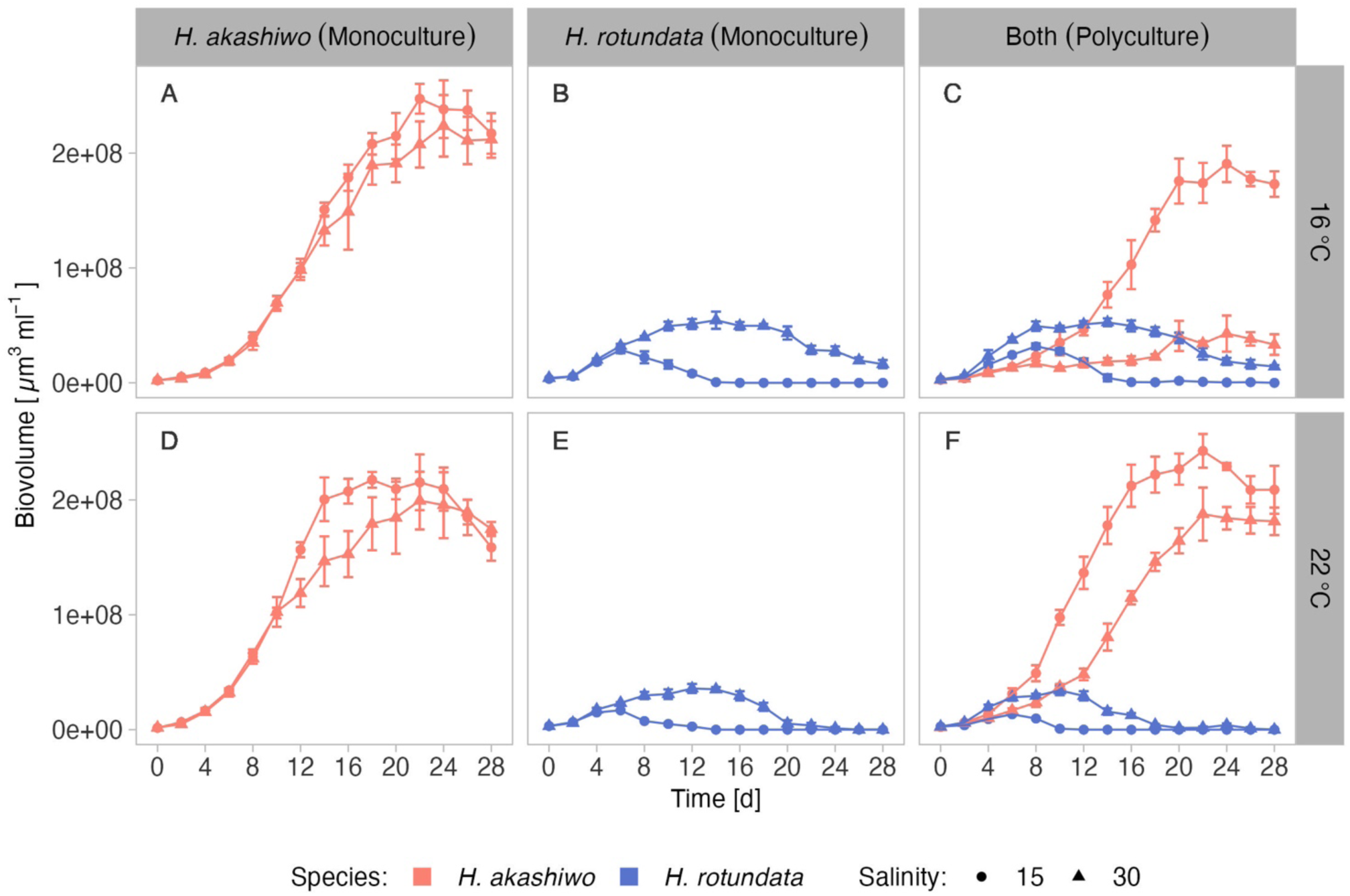
Mean biovolume ml ^-1^ of *H. akashiwo* and *H. rotundata* (indicated by color) over time ± SD incubated in monocultures and polycultures (facet columns) at two levels of salinity (15 and 30) represented by shape and two levels of temperature (16 °C and 22 °C) represented by facet rows.

**Table 2:**
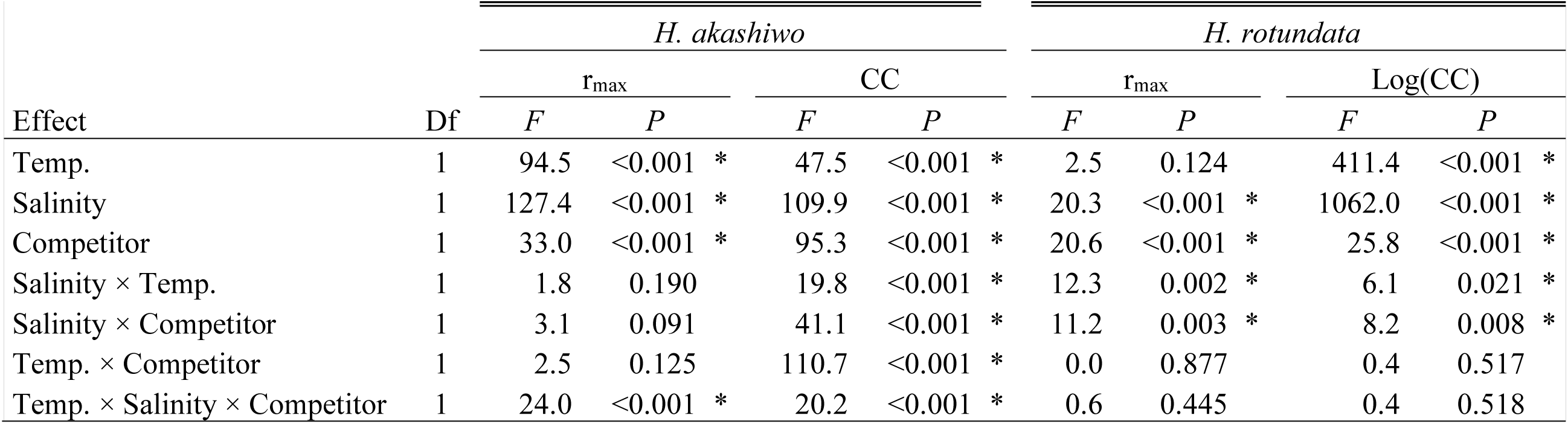
ANOVA results testing the effects of temperature, salinity and the presence of a competitor (Comp) on the maximum growth rate (r_max_) and carrying capacities (CC) of *H. akashiwo* and *H. rotundata* (monocultures and polyculture treatments together). Degrees of freedom (Df), F and p-values are given for each effect. Values marked with an asterisk (*) indicate significant effects (p < 0.05).

*H. rotundata* biovolume was promoted by the higher salinity level of 30 in combination with the lower experimental temperature of 16°C compared to the other treatment combinations (p < 0.001, Fig. 1b, 1e), which resulted in a maximum growth rate of 0.52 day^-1^ in this treatment and the highest carrying capacity (p < 0.001, Table S4). By contrast, under low salinity – high temperature conditions, *H. rotundata* biovolume production was strongly inhibited, reflected by a lower maximum growth rate compared to the other treatments (Table S3), and a maximum biovolume accounting for only 29% of the maximum biovolume achieved under the optimum high-salinity low-temperature conditions. Compared to *H. akashiwo,* the dinoflagellate generally exhibited lower total carrying capacities that were, however, achieved between day 6 and 14 depending on the treatment, and therefore several days earlier than for *H. akashiwo* in monoculture.

Temperature and salinity effects on *H. akashiwo* biovolume were generally stronger in polycultures compared to monocultures (Fig. 1c, 1f). Within the high temperature treatments, *H. akashiwo* showed slightly higher maximum growth rates at the lower salinity level and a suppressed carrying capacity at the higher salinity level accounting for only 78% of the carrying capacity achieved in the corresponding monoculture treatment. At higher temperature, *H. akashiwo* dominated over *H. rotundata* for most of the experimental duration (Fig. 1f, S1) indicated by significant differences of the log ratio from day 0 (Table S5). More precisely, within the high temperature treatments, *H. akashiwo* dominated over *H. rotundata* at the lower salinity level from day 2 onwards, whereas at the higher salinity level *H. rotundata* exhibited a temporary dominance from day 4 to 6, after which *H. akashiwo* dominated again from day 12 until the end of the experiment. Within the low temperature treatments, *H. akashiwo* dominated the low salinity treatment from day 12 onwards but was significantly inhibited by its competitor at high salinity until day 18, after which it dominated the community again. As such, competitive interactions strongly depended on salinity, especially within the low temperature treatments (Fig. 1c), which was also reflected by the significant interactive effects of temperature, salinity, and competitor presence for *H. akashiwo* biovolume, maximum growth rate, and carrying capacity (Table 1, Table 2). At low temperature – low salinity, *H. akashiwo* performed similar in the presence of *H. rotundata* as under the corresponding monoculture conditions. At the higher salinity level, *H. akashiwo* showed strongly suppressed growth rates and reached only 20% of the maximum biovolume of its corresponding monoculture.

Dissolved nutrient concentrations indicated phosphorous depletion in the *H. akashiwo* monocultures as well as in the polycultures starting from day 12 onwards (Fig. S2). Dissolved nitrogen was depleted several days later at day 20 (Fig. S3). However, C:N ratios across the different treatment combinations were relatively low and ranged from 5.1 to 9.1, indicating that algal cells were not N limited. C:N ratios were significantly affected by species composition, with *H. akashiwo* generally having higher C:N ratios than *H. rotundata* (p < 0.001), yet showing interactive effects with temperature and salinity (Fig. S4, Table S6).

### 4.2 Nauplii pilot study and grazing assay

In the preliminary pilot study, the mortality of the *A. tonsa* nauplii was higher when incubated with *H. akashiwo* compared to the filtered seawater control (p = 0.032). In the filtered seawater treatment, nauplii mortality showed a steady linear increase over time (Fig. S5), resulting in a mean mortality of 72% after five days of incubation. In contrast, mean nauplii mortality increased from 3 to 83% between day 1 and day 2 in the *H. akashiwo* treatment.

In the full factorial nauplii grazing assay conducted under the same temperature and salinity levels used in the competition experiment, nauplii mortality was interactively affected by salinity, temperature, prey composition, and time (Table 3, Fig. 2). Nauplii mortality at the end of the experiment was higher when incubated with *H. akashiwo* monocultures compared to *H. rotundata* monocultures or the polyculture treatments containing both phytoplankton species (p < 0.001, Table S7). In the *H. akashiwo* monocultures, nauplii mortality was higher at 22°C than at 16°C (p < 0.001), starting from day 2 onwards. Contrastingly, in the 16°C treatments, mortality was still 0% after two days of incubation and started to increase thereafter. Across both temperatures, nauplii mortality increased earlier at the lower salinity level compared to high salinity but converged in both salinities over the course of the experiment, which is reflected in the significant interaction of salinity and time (Table 3). The highest overall mortality was observed in *H. akashiwo* monocultures under low salinity – high temperature conditions, i.e., the optimal growth conditions of *H. akashiwo*, with the first experimental unit reaching a mortality of 100% on day 3. Treatments containing the non-harmful prey *H. rotundata,* either in monoculture or in polyculture with *H. akashiwo,* were similar to each other and showed low nauplii mortality over time with no notable differences among each other (Table S7). Regarding the developmental stages of the nauplii at the end of the experiment, nauplii exposed to *H. akashiwo* monocultures did not develop beyond the N1 state in any treatment combination of temperature and salinity, while nauplii in the *H. rotundata* monocultures and the polycultures developed to stages N4 and N3, respectively, at 16°C and even reached copepodite stages C2 – C4 in the 22°C treatments (Fig. S6).

**Table 3:** (a) Results of a linear mixed model (LMM) testing the effects of salinity, temperature, prey composition and day (fixed effects) on nauplii mortality with beaker number (id) as random factor. (b) Results of a generalized least squares model (GLS) testing the effects of salinity, temperature and prey composition on relative population grazing rate. Degrees of freedom (df), F and p-values are given for each effect. Values marked with an asterisk (*) indicate significant effects (p < 0.05).

| Fixed Effect | Mortality (LMM) |  |  | Grazing Rate (GLS) |  |  |
| --- | --- | --- | --- | --- | --- | --- |
|  | Df | F | p | Df | F | p |
| Salinity | 1 | 18.1 | <0.001 * | 1 | 3.0 | 0.092 |
| Temp. | 1 | 75.8 | <0.001 * | 1 | 104.2 | <0.001 * |
| Prey | 2 | 172.3 | <0.001 * | 2 | 31.1 | <0.001 * |
| Day | 5 | 57.0 | <0.001 * | - | - | - |
| Salinity × Temp. | 1 | 0.4 | 0.552 | 1 | 6.6 | 0.014 * |
| Salinity × Prey | 2 | 6.3 | 0.005 * | 2 | 5.3 | 0.009 * |
| Temp. × Prey Composition | 2 | 57.6 | <0.001 * | 2 | 21.5 | <0.001 * |
| Salinity × Day | 5 | 2.7 | 0.021 * | - | - | - |
| Temp. × Day | 5 | 11.6 | <0.001 * | - | - | - |
| Prey × Day | 10 | 32.9 | <0.001 * | - | - | - |
| Salinity × Temp. × Prey | 2 | 1.4 | 0.250 | 2 | 1.2 | 0.315 |
| Salinity × Temp. × Day | 5 | 1.7 | 0.147 | - | - | - |
| Salinity × Prey × Day | 10 | 1.3 | 0.213 | - | - | - |
| Temp. × Prey × Day | 10 | 8.0 | <0.001 * | - | - | - |
| Salinity × Temp. × Prey × Day | 10 | 1.2 | 0.301 | - | - | - |
| **Random Effect** |  |  |  |  |  |  |
| $\sigma^2$ | | 117.48 | | - | - | - |
| $\tau_{00 \text{ id}}$ | | 6.03 | | - | - | - |
| ICC |  | 0.05 |  | - | - | - |
| $N_{\text{id}}$ | | 48 | | - | - | - |
| Observations |  | 288 |  | - | - | - |
| Marginal $R^2$ / Conditional $R^2$ | | 0.836 / 0.844 | | - | - | - |

**Figure 2:**
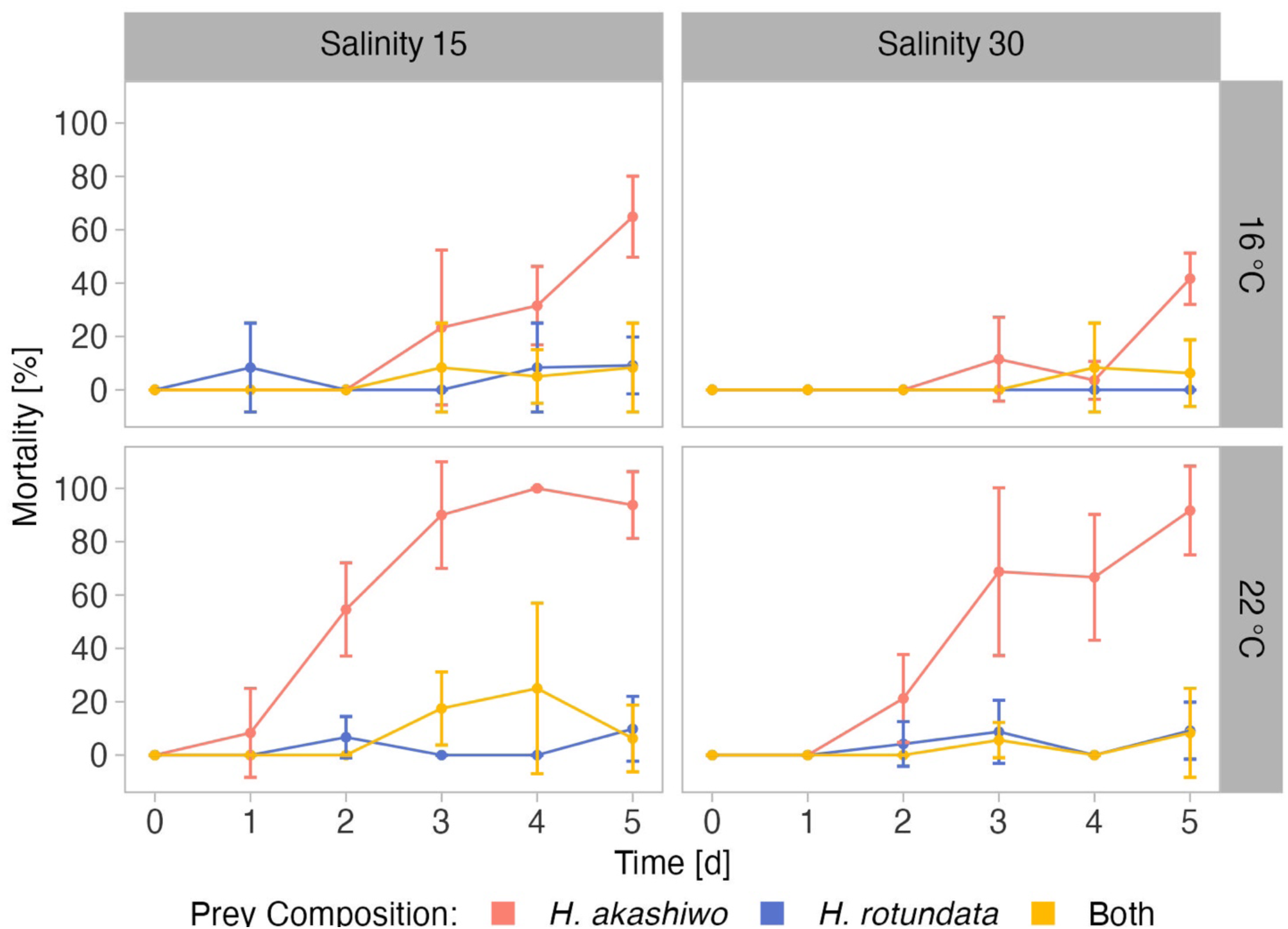
Mortality of *A. tonsa* nauplii over time when exposed to *H. akashiwo*, *H. rotundata* or both algae species in combination (indicated by color) across two levels of temperature (16 °C and 22 °C) and salinity (15 and 30) represented by facet grid rows and columns, respectively.

Population grazing rates of the *A. tonsa* nauplii were significantly and interactively affected by prey composition and temperature, while the salinity effect was not significant, but depended on prey composition and temperature (twofold interactions of salinity with both factors, Table 3). Average nauplii grazing rates were lowest in *H. akashiwo* monoculture treatments irrespective of temperature and salinity levels (Fig. 3). In the *H. rotundata* monoculture as well as in the polyculture treatments, however, nauplii grazing rates strongly differed with temperature and salinity. While grazing rates were generally very low at 16°C and did not exceed a mean of 0.1 day^-1^, grazing rates increased with temperature, especially at low salinity, resulting in significant differences of this treatment compared to most other treatment combinations (Table S8). *H. rotundata* monoculture treatments experienced the highest grazing impact (0.39 day^-1^). Considering separate grazing rates for both phytoplankton species in the polyculture treatments, grazing rates on *H. rotundata* were higher than on *H. akashiwo* in the higher experimental temperature (p < 0.001, Fig. 4, Table 4, S9), regardless of salinity.

**Figure 3:**
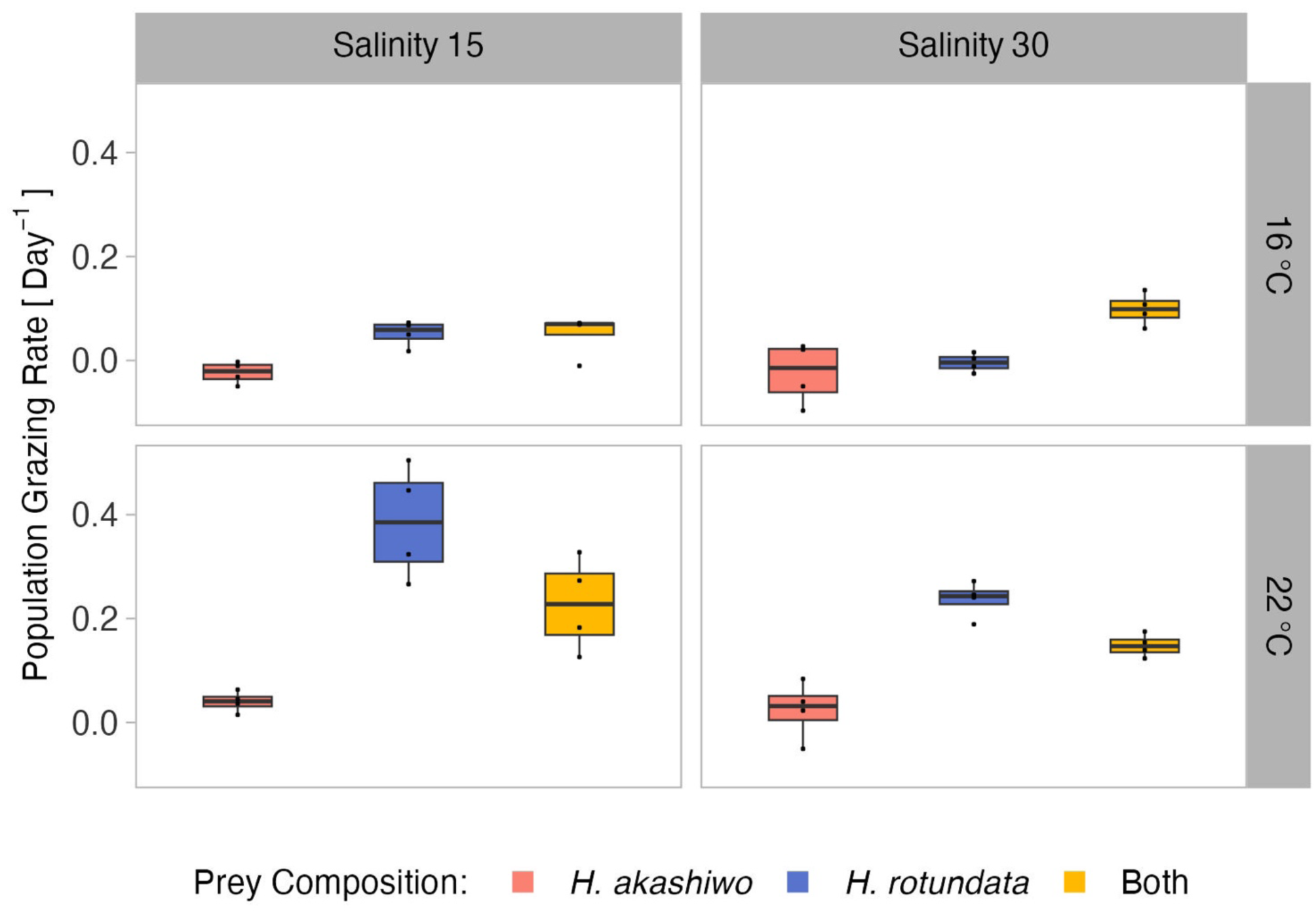
*A tonsa* nauplii relative population grazing rate day^-1^at the different prey algae compositions *H. akashiwo*, *H. rotundata* and both algae in combination (indicated by color) across two levels of temperature (16 °C and 22 °C) and salinity (15 and 30) represented by facet grid rows and columns, respectively.

**Figure 4:**
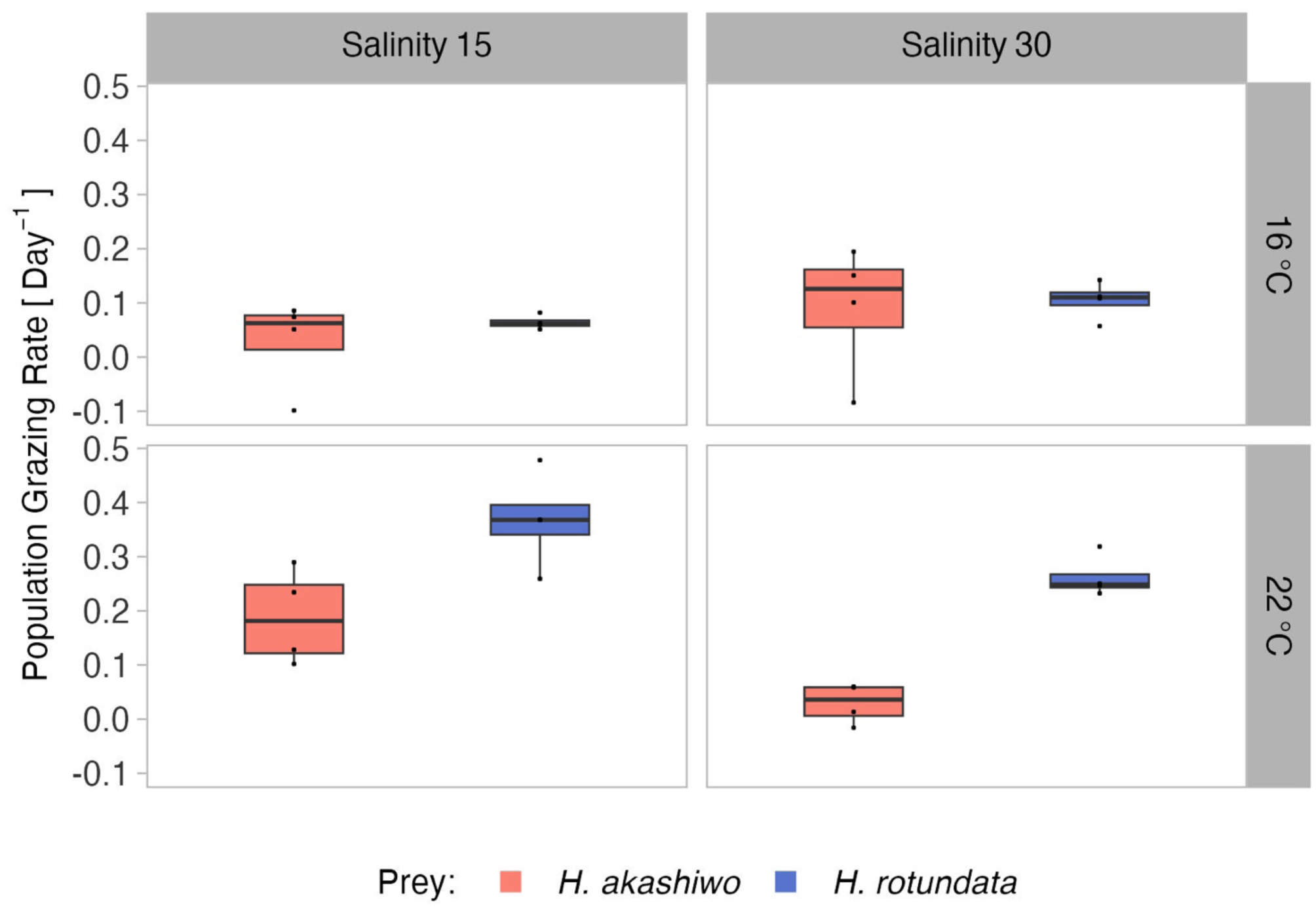
Relative population grazing rates of the selective feeding *A tonsa* nauplii in the polyculture treatments separated by prey algae species (indicated by color) across two levels of temperature (16 °C and 22 °C) and salinity (15 and 30) represented by facet grid rows and columns, respectively.

**Table 4:** ANOVA results testing differences among grazing rates in the polyculture treatments as a response to algae species identity, temperature and salinity for *A. tonsa* and *B. plicatilis*. Degrees of freedom (df), F and p-values are given for each effect. Values marked with an asterisk (*) indicate significant effects (p < 0.05).

| Effect | *A. tosa* Grazing Rate<br>(polyculture) |  |  | *B. plicatilis* Grazing Rate<br>(polyculture) |  |  |
| --- | --- | --- | --- | --- | --- | --- |
|  | Df | F | p | Df | F | p |
| Prey | 1 | 20.2 | <0.001 * | 1 | 4.1 | 0.079 |
| Temp. | 1 | 29.6 | <0.001 * | - | - | - |
| Salinity | 1 | 2.5 | 0.128 | 1 | 2.4 | 0.160 |
| Prey × Temp. | 1 | 12.4 | 0.002 * | - | - | - |
| Prey × Salinity | 1 | 0.1 | 0.761 | 1 | 16.1 | 0.004 * |
| Temp. × Salinity | 1 | 12.9 | 0.001 * | - | - | - |
| Prey × Temp. × Salinity | 1 | 0.5 | 0.475 | - | - | - |

Dissolved nutrient concentrations indicated no macronutrient depletion of phosphorous or nitrogen during the experiment (Fig. S7, S8). C:N ratios of the phytoplankton ranged between 6.7 and 8.2 (Fig. S9).

### 4.3 Rotifer grazing assay

The second grazing assay was conducted with the rotifer *B. plicatilis* at only one temperature (16°C), but both salinity levels which were also used in the nauplii grazing assay. Here, rotifer mortality was significantly affected by prey composition and time; both factors depending on salinity (two-fold interactive effects of both factors with salinity, Fig. 5, Table 5).

**Figure 5:**
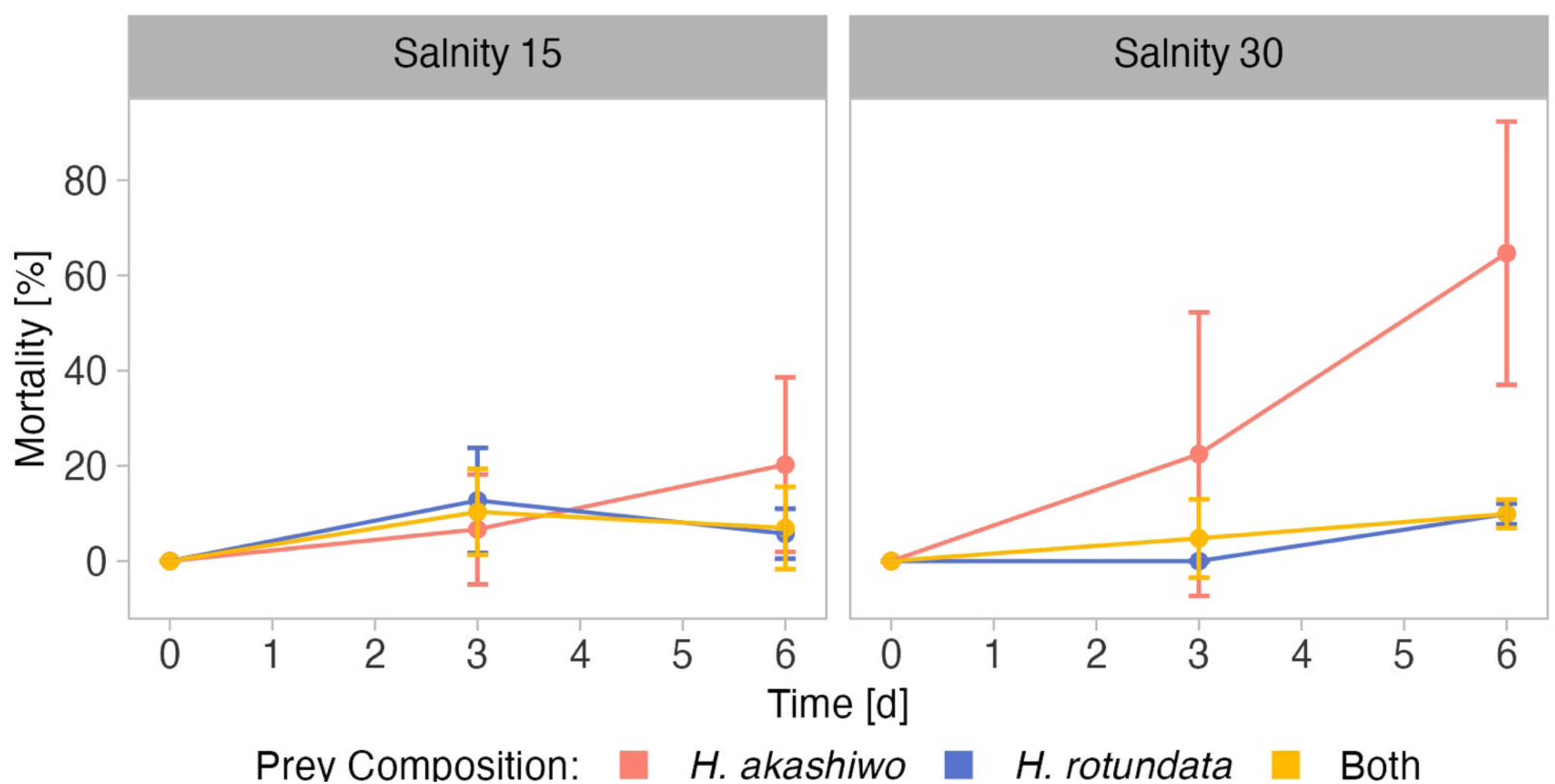
Mortality of *B. plicatilis* over time when exposed to *H. akashiwo*, *H. rotundata* or both algae species in combination (indicated by color) across two levels of salinity (15 and 30) represented by facet grid columns.

**Table 5:** ANOVA results testing the effects of salinity and prey composition on *B. plicatilis* mortality and population grazing rate. Degrees of freedom (df), F and p-values are given for each effect. Values marked with an asterisk (*) indicate significant effects (p < 0.05).

| Effect | Mortality (AOV) |  |  |  | Grazing Rate (AOV) |  |  |  |
| --- | --- | --- | --- | --- | --- | --- | --- | --- |
|  | Df | *F* | *p* |  | Df | *F* | *p* |  |
| Salinity | 1 | 6.5 | 0.025 | * | 1 | 36.2 | <0.001 | * |
| Prey Composition | 2 | 11.6 | 0.002 | * | 2 | 1.4 | 0.290 |  |
| Salinity × Prey Composition | 2 | 4.1 | 0.043 | * | 2 | 8.1 | 0.006 | * |

As in the nauplii grazing assay, grazer mortality at the last sampling day was higher in *H. akashiwo* monoculture treatments compared to the *H. rotundata* monoculture or polyculture treatments (p < 0.004, Table S10). Within the *H. akashiwo* treatments, the highest mortality was found at the high salinity level (p = 0.023) with a mean mortality of 65% at the end of the experiment, which was notably higher than the mortality at the low salinity level (20%). Rotifer mortality in treatments containing the non-harmful prey *H. rotundata* in monoculture or in polyculture with *H. akashiwo* did not differ from each other and did not exceed a mean mortality of 10%.

The relative population grazing rates were very low across all treatments and included negative values, indicating that for some treatments phytoplankton growth rates were even higher in the presence of grazers compared to the non-grazed controls (Fig. 6). However, the relative population grazing rate of *B. plicatilis* differed with salinity as well as interactively with salinity and prey composition (Table 5). This was reflected in similar grazing rates across different prey compositions at the low salinity level, but significant differences at the higher salinity level (Table S11). However, despite these significant differences, the overall grazing impact of the rotifers and therefore also the effect size was very low.

**Figure 6:**
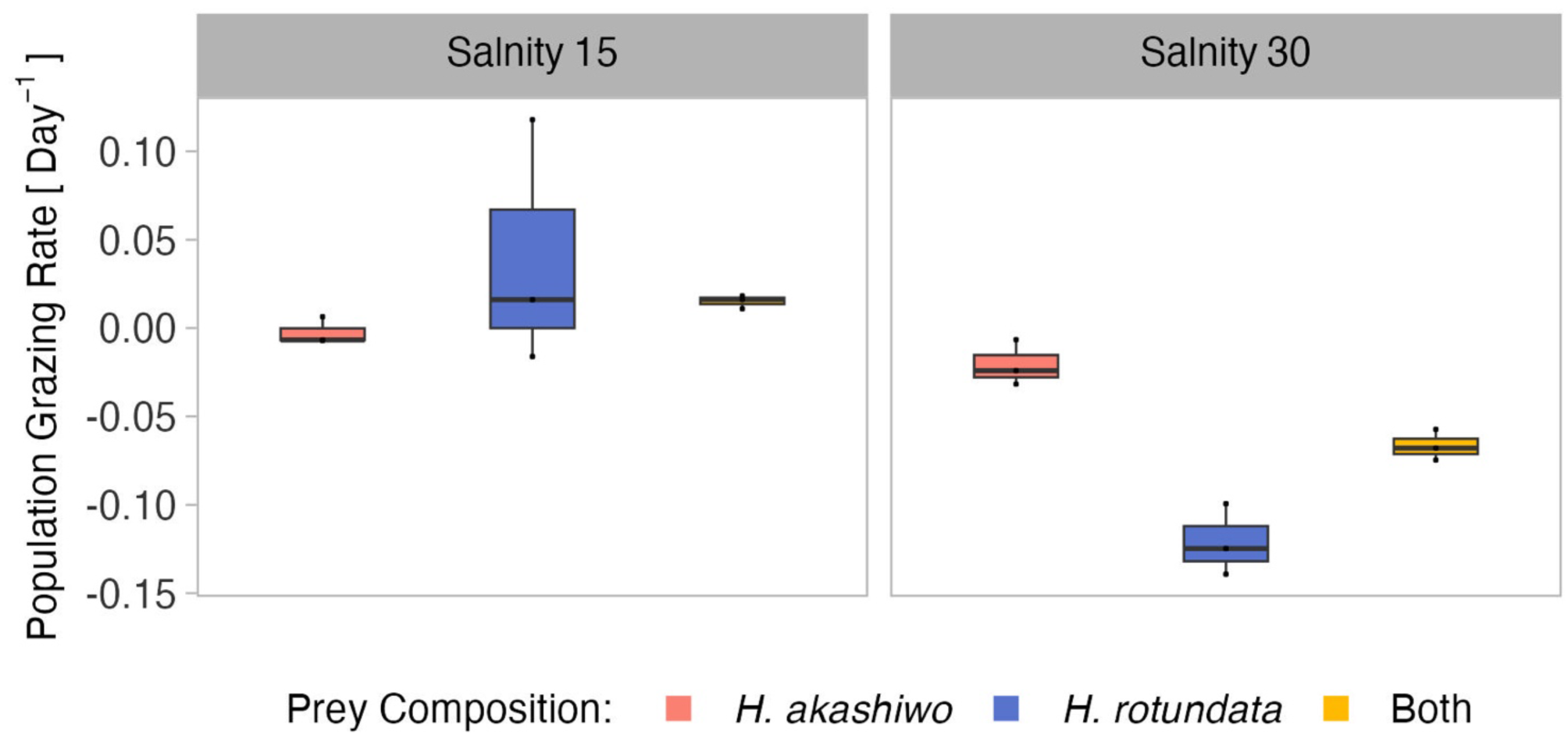
*B. plicatilis* relative population grazing rate day^-1^at the different prey algae compositions *H. akashiwo*, *H. rotundata* and both algae in combination (indicated by color) across two levels of salinity (15 and 30) represented by facet grid columns.

In polycultures, grazing rates on *H. rotundata* were higher than for *H. akashiwo* at the lower salinity level (p = 0.012, Table 4, S12). Yet, differences between the species-specific relative grazing rates were also relatively small (< 0.1) and are likely an artefact of the enhanced growth of *H. akashiwo* in this treatment since *B. plicatilis* is known to have a non-selective feeding behavior (Fig. S10).

Comparing the mortality of *B. plicatilis* and *A. tonsa* nauplii in the 16°C treatments, grazer mortality was significantly and interactively affected by prey composition and time (significant main factors and interaction), while salinity effects were only important in relation to time (Table S13). Grazer mortality did not differ notably between the two consumer species, indicating similar sensitivities of both grazers in response to the harmful algae. However, salinity affected grazer mortality in *H. akashiwo* monocultures in contrasting ways, with *B. plicatilis* exhibiting higher mortality at high salinity and nauplii showing higher mortality at the low salinity level (Fig. 2, Fig. 5, Table S14).

Population grazing rates of the two grazers differed from each other and were interactively affected by salinity and prey composition (significant main effects and two-way interactions, Table S13). The grazing rates of the two consumers did not exceed an average of 0.1 day^-1^ in the lower temperature level. The observed interactive effect of consumer identity and prey composition indicated that the presence of non-harmful prey led to higher grazing rates for the *A. tonsa* nauplii. In contrast, the presence of non-harmful prey did not result in increased grazing for *B. plicatilis* based on its overall minor grazing impact across all factorial combinations (Fig. 3, Fig. 6, Table S15).

## 5 Discussion

As expected, based on field observations, *H. akashiwo* was promoted at high temperature – low salinity conditions, while *H. rotundata* proliferated at the contrary low temperature – high salinity conditions. In competition, *H. akashiwo* dominated the system at the higher temperature regime irrespective of salinity. At low temperature, however, salinity determined the competitive outcome, promoting *H. akashiwo* over *H. rotundata* at low salinity and vice versa at high salinity. Both studied grazers were negatively affected by *H. akashiwo* in monoculture; however, salinity levels altered these effects in different ways, leading to higher mortality at low salinity for *A. tonsa* nauplii and at high salinity for the rotifer *B. plicatilis*. Furthermore, for both grazers, the presence of non-harmful prey significantly alleviated the harmful effects caused by *H. akashiwo.* In contrast to the non-selective filter feeder *B. plicatilis*, the highly selective *A. tonsa* nauplii strongly avoided *H. akashiwo* in the polyculture treatments.

The observed growth preferences of *H. akashiwo* match the findings of Lemley et al. (2018b), who identified optimum growth conditions for *H. akashiwo* at elevated water temperatures of 20 – 26 °C and a salinity preference towards brackish conditions (4–24) in the Sundays Estuary. Similar patterns could be observed in the Golden Horn Estuary in the northeast Sea of Marmara, Turkey, where a short-term increase in water temperature from 15.7°C to 20.2°C in combination with a decrease in salinity from 18.7to 16.4 triggered a *H. akashiwo* HAB with a density of 10^3^ cells ml^-1^ (Dursun et al., 2016). However, despite the identification of its optimum growth conditions in the present study, *H. akashiwo* also exhibited its previously reported high tolerance against broad ranges of temperature and salinity regimes outside of its optimum growth window (Strom et al., 2013, Martinez et al., 2010, Allaf and Trick, 2019). This was evidenced by only minor performance differences across treatments in monoculture as well as by dominating three out of four temperature – salinity combinations in competition.

The performance of the dinoflagellate competitor *H. rotundata* in competition with *H. akashiwo* was similar to its respective monoculture growth controls, although *H. rotundata* was shown to decline in the presence of *H. akashiwo* in the field (2018a). Our laboratory study implies a low suppressive potential of the studied South African *H. akashiwo* strain against its competitor under the selected experimental conditions. Yamasaki et al. (2007) indicated that the harmful substances produced by *H. akashiwo* may be highly species-specific. In their study, *H. akashiwo* substantially suppressed the growth of the harmful diatom species *Chaetoceros muelleri* and *Skeletonema costatum*, while having no harmful effect on cultures of the dinoflagellate *Prorocentrum cordatum.* Another possible explanation for the lack of a negative impact of *H. akashiwo* on *H. rotundata* in our study could be the higher growth rate of *H. rotundata* compared to *H. akashiwo*, resulting in a distinct time lag for the two species to reach their carrying capacity. Consequently, this could have led to *H. rotundata* already being naturally in decline by the time *H. akashiwo* reached cell densities at which allelopathy became relevant. This advantageous high growth rate could represent a protection strategy to avoid elevated concentrations of allelopathic or other harmful substances (Arzula et al., 1999). This idea matches the results of Yamasaki et al. (2007), who demonstrated that substantial inhibitory effects by *H. akashiwo* on its competitors did not come into effect before the end of their exponential growth phase, indicating that the extent of the harmful effect is not only dependent on cell concentration, but also on the specific growth phase of *H. akashiwo*.

Interestingly in this study, however, the negative effect of *H. rotundata* on *H. akashiwo* growth under high salinity – low temperature conditions point towards an inhibitory potential of *H. rotundata*. Although it is generally known that *H. rotundata* can form red tides in terms of large monospecific bloom events, this species has not been associated with direct harmful effects on competitors or consumers so far. Therefore, the suppressed growth of *H. akashiwo* observed in mixed culture with *H. rotundata* in this treatment is more likely based on a competitive advantage of the dinoflagellate under these environmental conditions. Given the fact that the total biovolume in polyculture was twice as high as in the monocultures based on the additive experimental design, it is likely that interspecific competition in polycultures constrained populations dynamics before intraspecific competition did in monocultures.

Since *H. akashiwo* is a globally occurring species, experimental evidence for growth preferences of a certain strain from a particular habitat raises the question to which degree these findings can be transferred to other strains in other systems, i.e., what is the extent of intraspecific trait variation for this species. Martinez et al. (2010) investigated the growth responses of six different strains of *H. akashiwo* originating from a broad latitudinal range at different levels of temperature, salinity, and light intensity and demonstrated an across-strain preference for the higher experimental temperature of 23°C (vs. 17°C). Salinity preferences depended on strain identity, yet all strains showed a fast adaptive capability to different salinity regimes. The high halotolerance of *H. akashiwo* was intensively studied by Strom et al. (2013), who indicated that this tolerance is likely to be an evolutionary defense strategy developed by this species against marine micrograzers that allows it to escape by inhabiting low salinity surface waters that are unsuitable for marine grazers.

Copepod nauplii exposed to *H. akashiwo* monocultures in this study showed increased mortality as well as inhibited grazing activity, implying a high sensitivity of these grazers to the harmful effects of *H. akashiwo*. This was also confirmed in our pilot grazing assay in which the negative effects of *H. akashiwo* on *A. tonsa* nauplii far exceeded those of pure starvation in the filtered seawater control. Furthermore, the selective feeding behavior indicated that the nauplii were able to recognize the prey as harmful and actively avoid it. Since the C:N ratios of *H. akashiwo* were only marginally higher than the Redfield C:N ratio, this grazer deterrence effect was likely not based on low algal food quality, although unfortunately no particulate organic phosphorous data could be evaluated to further support this finding.

As for the *A. tonsa* nauplii in our study, Colin and Dam (2002) demonstrated suppressed ingestion rates, as well as reduced survival and egg production in adult *A. tonsa* copepods when restricted to *H. akashiwo* monoculture prey. In the present study, nauplii mortality was highest in high temperature – low salinity treatments, thus coinciding with the optimum growth conditions of *H. akashiwo*. These stronger negative effects of *H. akashiwo* at high versus low temperature are contrary to previous studies that investigated brevetoxin production (Ono et al., 2000) and cell permeability as a proxy for cytotoxicity of *H. akashiwo* (Allaf and Trick, 2019) at different levels of temperature, salinity, and light. Ono et al. (2000) found a maximum toxicity of a *H. akashiwo* strain originating from the Seto Inland Sea, Japan, at 20°C (vs. 25°C), whereas Allaf and Trick (2019) found the highest cell permeability at 15°C (vs. 20 and 25°C) in a *H. akashiwo* strain isolated from Clam Bay, USA. Thus, both studies demonstrated an increasing cytotoxicity of *H. akashiwo* with decreasing temperature. Yet, both studies did not test the actual adverse effects on primary consumers at the studied conditions, therefore neglecting how different environmental conditions could alter the susceptibility of grazers towards the harmful substances produced by *H. akashiwo*. Furthermore, the discrepancies to the present study could potentially be explained by strain-specific differences in the predominant mode of harmfulness of this species. For instance, John et al. (2015) demonstrated that within a toxigenic population of the dinoflagellate *Alexandrium fundyense*, single strains were capable of allelopathy to protect the population from grazing, while other strains did not show any allelopathic capabilities, but produced potent saxitoxins. In the present study, neither toxin production nor cell permeability were determined. However, based on observations during the experiments, mucus production by *H. akashiwo* likely played a major role in causing the lethal effects on both grazers in this study, as after 24h, for instance, the majority of nauplii and rotifers were trapped in mucus and were strongly immobilized. Similar observations were made by Xie et al. (2007) who also found an enhanced immobilization of rotifers due to mucus produced by *H. akashiwo*. As harmful effects of *H. akashiwo* on the nauplii exceeded those of pure starvation, it is likely that other harmful substances were involved in the high consumer mortality observed here, which are either excreted together with the mucus or are located on the cell surface of *H. akashiwo*, causing harmful effects upon cell contact (Yamasaki et al., 2009).

The fact that the strongest effects on nauplii mortality in this study were found in the *H. akashiwo* monocultures at high temperature conditions could be based on increased metabolic rates of *H. akashiwo*, resulting in increased net mucus production at higher temperature. Additionally, the immobile nauplii in the *H. akashiwo* monocultures were likely to use up their energy reserves more rapidly at higher temperature due to increased metabolic rates, but also due to their increased physical efforts to detach themselves from the mucus. This also supports the pattern that the temperature effect on mortality in *H. akashiwo* monocultures was much larger than the salinity effect, which was in contrast to that observed for *B. plicatilis*, indicating that salinity affected consumer fitness rather than the harmfulness of *H. akashiwo*.

In contrast, nauplii mortality was much lower in treatments containing *H. rotundata*, either in monoculture or in polyculture with *H. akashiwo*, which indicates that the presence of non-harmful prey can substantially buffer the adverse effects caused by *H. akashiwo*. This further matches the findings of Colin and Dam (2002), who demonstrated that reduced survival, ingestion and egg production rates of adult *A. tonsa* individuals induced by *H. akashiwo* were alleviated when non-harmful prey was present as an alternative food source. In addition to providing an alternative high-quality food source, the adhesive nature of potential allelochemicals (Branco et al., 2014) may cause them to attach to the non-harmful algae and thus reduce the ambient concentration of these substances for primary consumers. Similar suggestions were made by Prince et al. (2008), who found growth inhibitory effects caused by allelopathic compounds exuded by the red-tide-forming dinoflagellate *Karenia brevis* to be reduced in the presence of the diatom species *Skeletonema costatum*.

The observed lethal effects of *H. akashiwo* on *B. plicatilis* match the findings of Xie et al. (2007), who also showed a population decrease of *B. plicatilis* when exposed to *H. akashiwo* monocultures. Moreover, the presence of the dinoflagellate *Alexandrium tamarense* in polyculture with *H. akashiwo* alleviated the population decrease in a similar way as the presence of *H. rotundata* reduced the mortality of the rotifers in this study. Comparing the sensitivity of *A. tonsa* nauplii and *B. plicatilis* to *H. akashiwo* exposure, similar harmful effects were found for both tested metazoan grazers. Interestingly, salinity had opposite effects on the mortality of the two grazers, resulting in the highest mortality of *A. tonsa* nauplii at the low salinity level, while *B. plicatilis* experienced the highest mortality at the higher salinity level. This interactive effect of prey composition, salinity, and consumer identity indicates that although both grazers can be found in estuarine ecosystems (Paffenhöfer and Stearns, 1998), *A. tonsa* generally prefers a higher salinity level, whereas *B. plicatilis* is promoted by a lower salinity level. Consequently, when exposed to non-optimum salinities, the grazers need to spend more energy on osmoregulatory processes (Goolish and Burton, 1989), which consequently might reduce their fitness and lead to higher mortality when exposed to unfavorable prey.

Overall, the present study highlighted the growth dynamics of two dominant bloom-forming phytoplankton species in estuarine ecosystems, and demonstrated their preference for contrary temperature and salinity regimes. The observed growth optima for the two species match field observations and contribute to the explanation of alternating bloom occurrence patterns of *H. akashiwo* and *H. rotundata* in the field (Rothenberger et al., 2009, Lemley et al., 2018b, Hilmer and Bate, 1991). Moreover, *H. akashiwo* could only be outcompeted by *H. rotundata* at high salinity – low temperature conditions but dominated at low salinity – low temperature and at high temperature irrespective of salinity, emphasizing the importance of the interplay of natural abiotic conditions and their variability in estuarine ecosystems for determining the bloom dynamics of this species. Environmental variability in estuaries can be induced by weather changes or openings and closings of the river mouth, representing environmental resetting events which are necessary to constrain monospecific *H. akashiwo* blooms and thus strengthen ecosystem resilience. However, anthropogenic regulating interventions like the construction of barrages and water extraction for agricultural purposes reduce flow velocity and lead to permanently closed river mouth states, which in turn favor optimum bloom conditions of *H. akashiwo* (Lemley et al., 2018b, Dalu et al., 2018). This impeded abiotic variability inducing the formation and persistence of *H. akashiwo* blooms may increase even further in the future, especially under global warming additionally promoting the harmful raphidophyte, and can only be controlled by other measures that have the potential to reduce HAB events, such as the reduction of nutrient input to the ecosystem. In future studies, it is therefore crucial to consider the effects of inorganic and organic nutrient pollution in concert with other environmental factors to derive a comprehensive understanding of the bloom dynamics of *H. akashiwo* and *H. rotundata* in estuarine systems. Furthermore, this study demonstrated negative effects of *H. akashiwo* on copepod nauplii as well as rotifers, which likely have been caused by mucus production and potentially associated harmful substances of *H. akashiwo.* The poorly characterized nature of this mucus opens up an important research perspective to identify its chemical composition and quantify its production under different environmental conditions as well as its immobilizing and lethal effects on different targets. In an ecosystem context, however, not only the direct adverse effects on zooplankton need to be considered, but also the consequences of a potential decline in key estuarine primary consumers for higher trophic levels, as well as interactive effects with the temporally and spatially highly variable environment.

## Data accessibility

All data are provided in the electronic supplementary material and can be found in a GitHub repository (https://github.com/jakobgiesler/Heterosigma)

## Acknowledgements

We thank Heike Rickels (ICBM Terramare, Wilhelmshaven, Germany) for analysis of dissolved and particulate nutrient samples as well as technical support in the lab. Furthermore, we also thank Jan-Claas Dajka (HIFMB, Oldenburg, Germany) for advice in data analysis and Cedric Meunier (AWI, Helgoland, Germany) for providing the *A. tonsa* eggs.

## Authors’ contributions

SM, JA and DL conceived the idea and designed the study. JG conducted the experiments, performed the statistical analyses and drafted the manuscript with input from all co-authors. All authors contributed to the article and approved the submitted version.

