## Supplementary material for "Interactive effects of salinity, temperature and food web configuration on performance and harmfulness of the raphidophyte *Heterosigma akashiwo*"

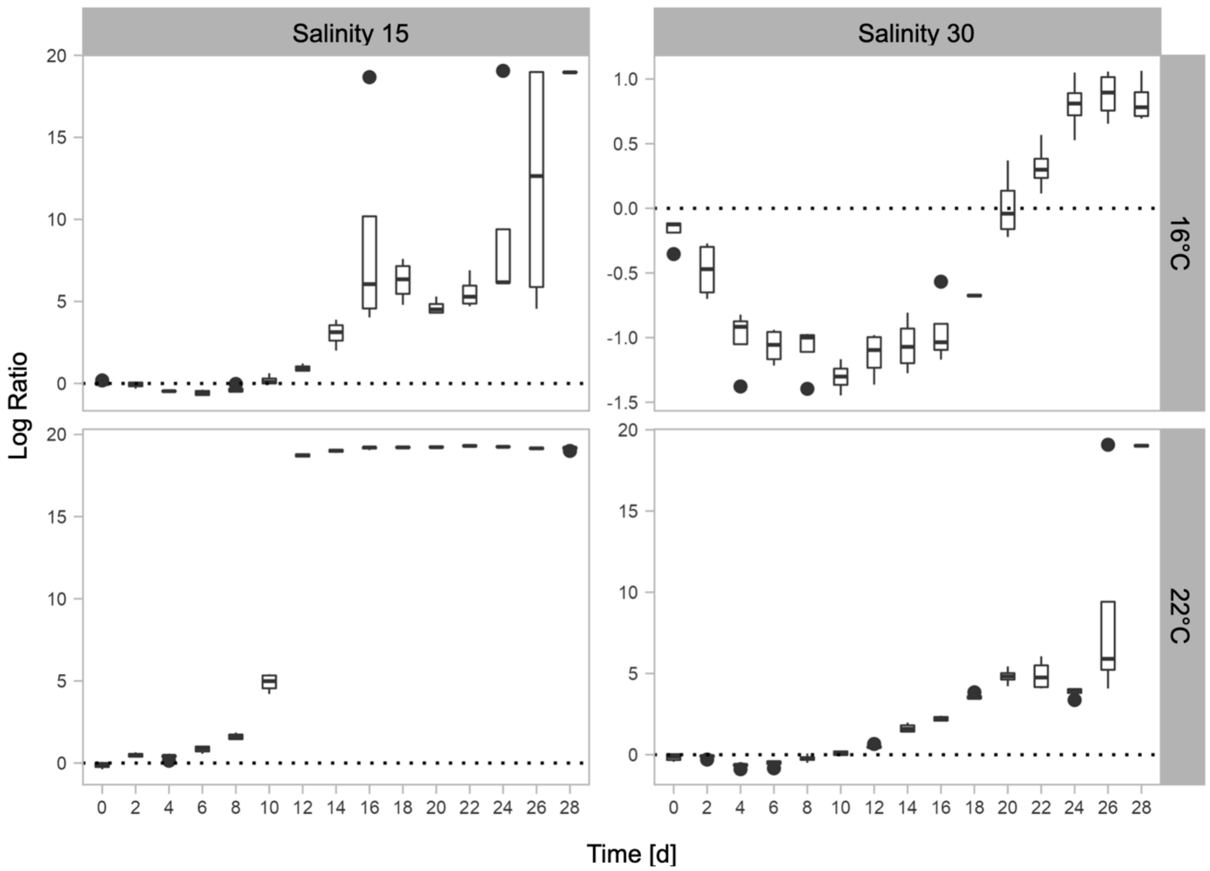


**Figure S1:** Log ratios of the polycultures calculated from *H. akashiwo* biovolumes divided by *H. rotundata* biovolumes at two levels of temperature (facet columns) in combination with two levels of salinity (facet rows) over time. Values > 0 (above dashed line) represent *H. akashiwo* dominance, values < 0 (below red dashed line) represent *H. rotundata* dominance.

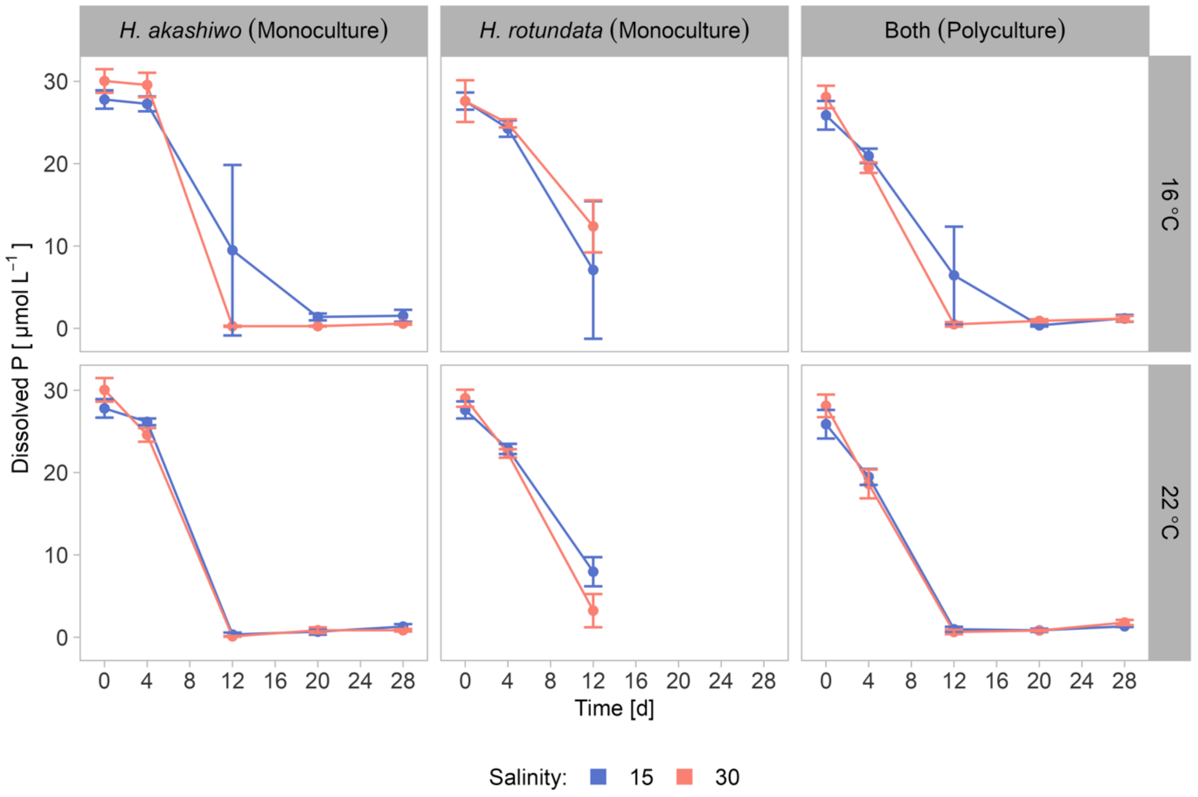


**Figure S2**: Dissolved phosphate concentrations [$\mu$mol L$^{-1}$] ± SD over time in the competition experiment comprising two levels of temperature (line type) in combination with two levels of salinity (facet columns) across the three species compositions (*H. akashiwo* monoculture, *H. rotundata* monoculture as well as both species together in polyculture).

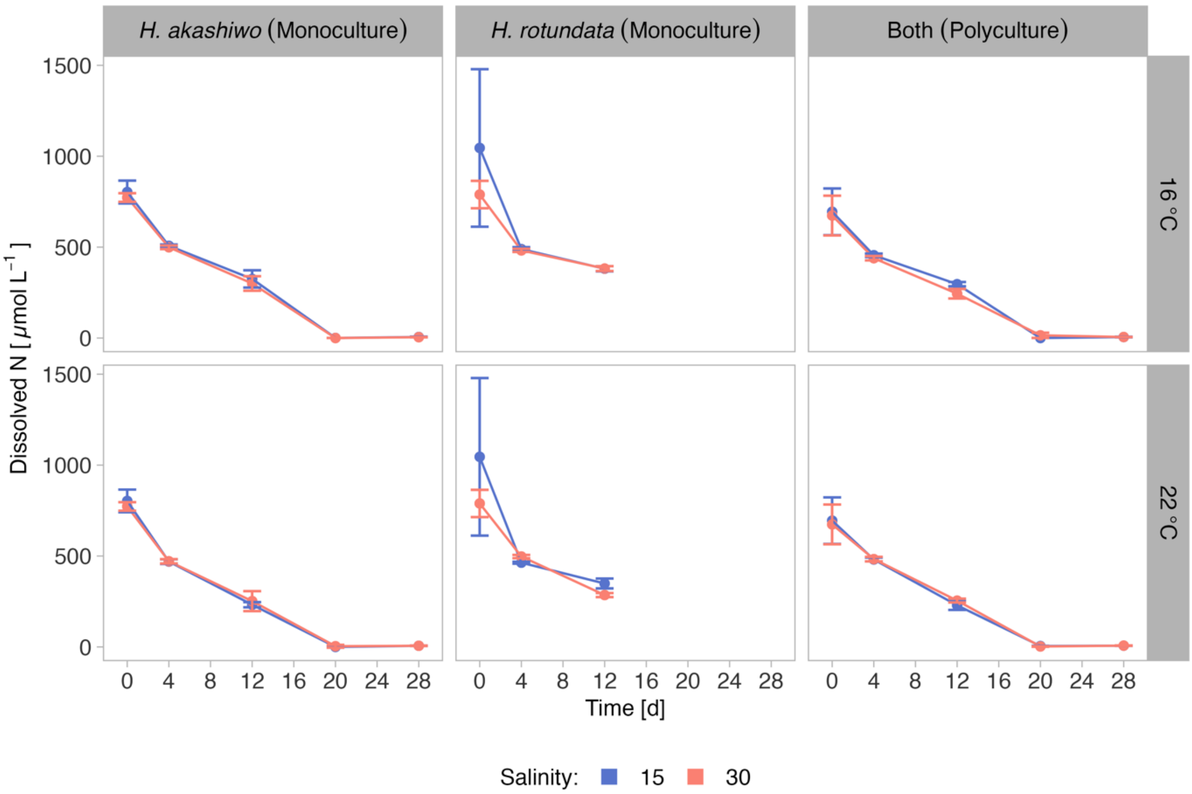


**Figure S3**: Dissolved nitrate and nitrite concentrations [μmol L$^{-1}$] ± SD over time in the competition experiment comprising two levels of temperature (line type) in combination with two levels of salinity (facet columns) across the three species compositions (*H. akashiwo* monoculture, *H. rotundata* monoculture and both species together in polyculture).

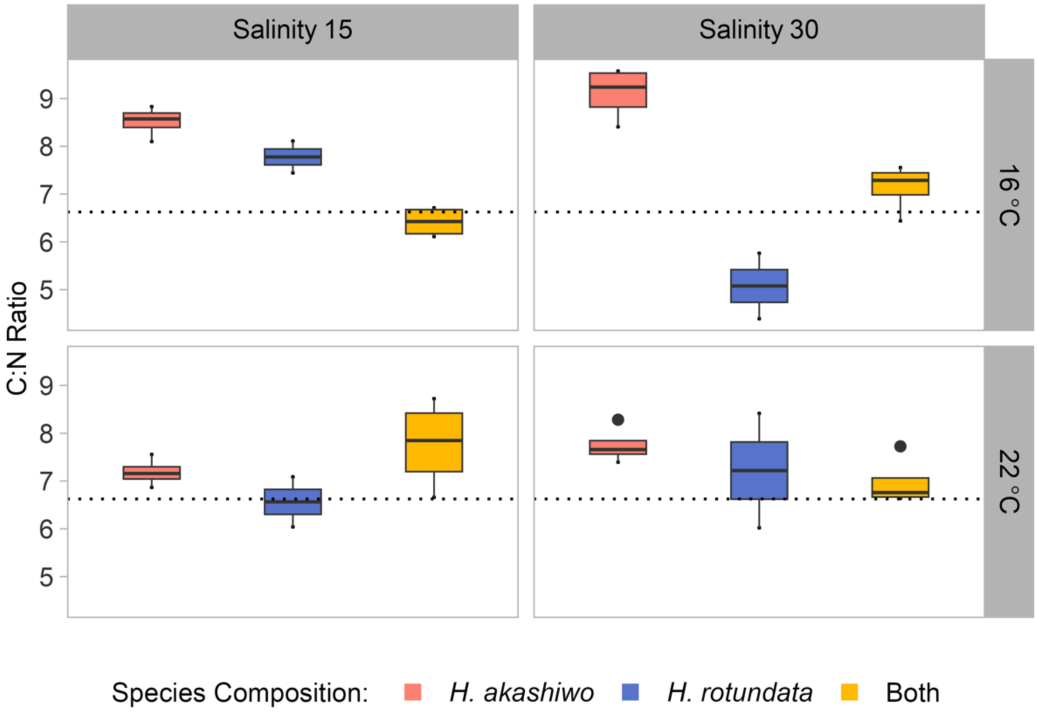


**Figure S4:** Particulate C:N ratio of *H. akashiwo*, *H. rotundata* and both species in polyculture across two levels of temperature (facet rows) in combination with two levels of salinity (facet columns) at day 28 of the competition experiment. The red field ratio is indicated by the dashed line.

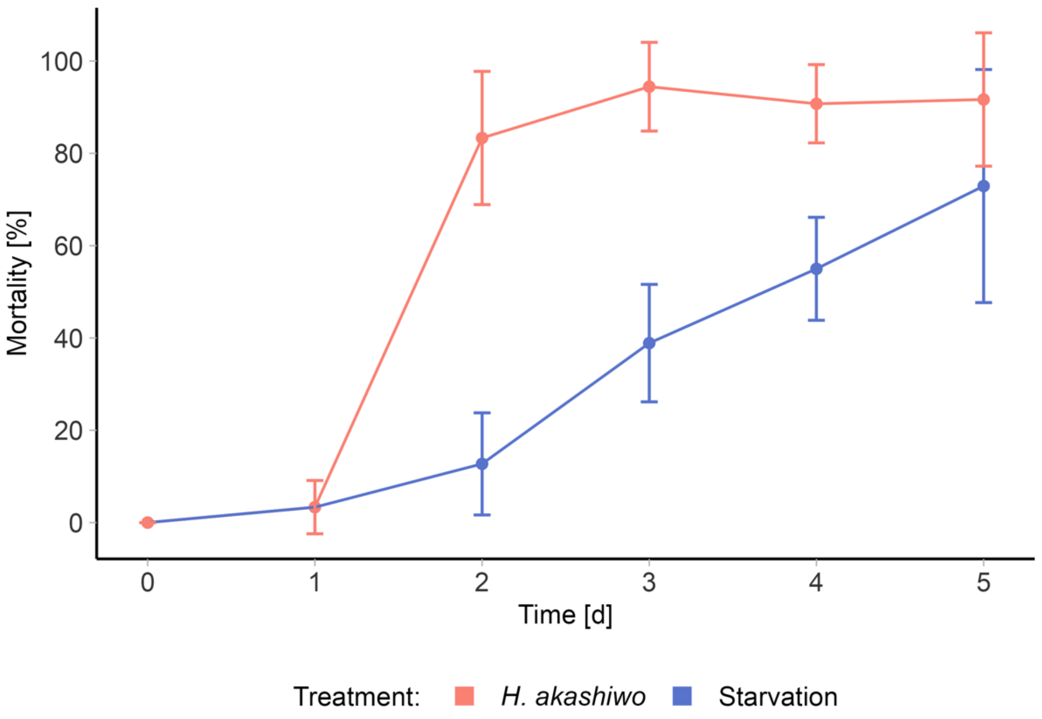


**Figure S5:** Results of *A. tonsa* nauplii pilot study: Nauplii mortality [%] ± SD over time [d] when exposed to 40.000 cells $ml^{-1}$ *H. akashiwo* culture (red) and filtered seawater (blue).

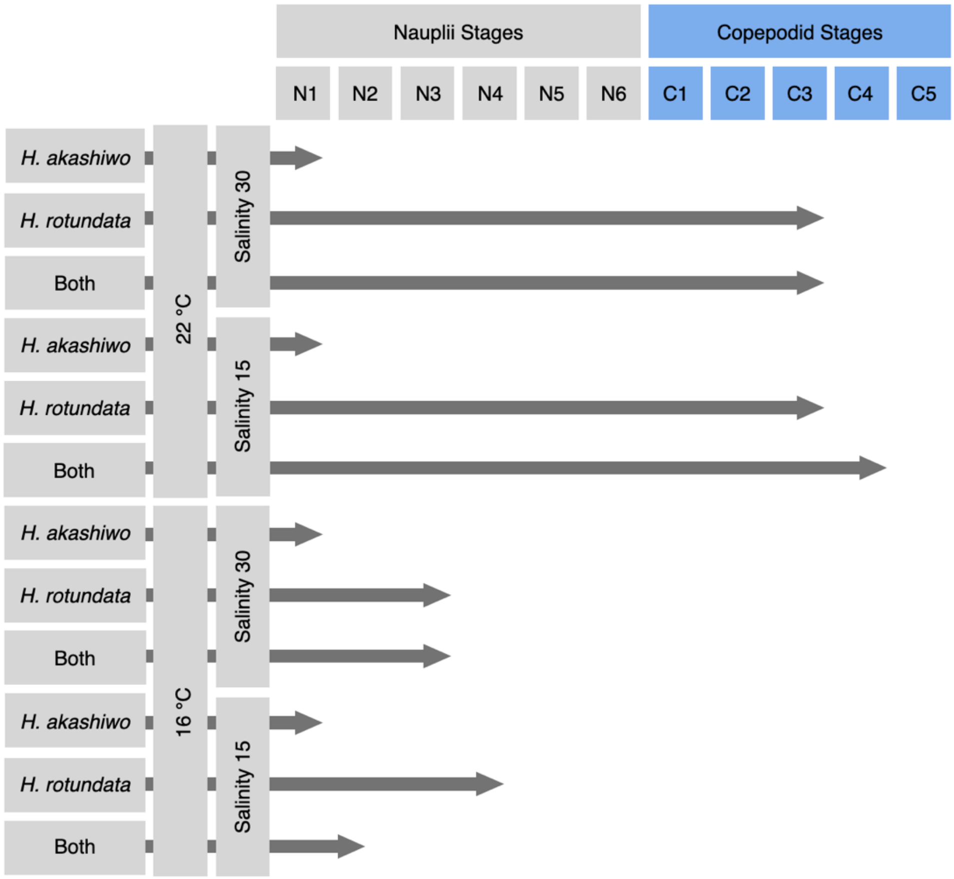


**Figure S6:** *A. tonsa* developmental stages at final sampling day of the full-factorial grazing assay across three different prey compositions, two levels of temperature and two levels of salinity.

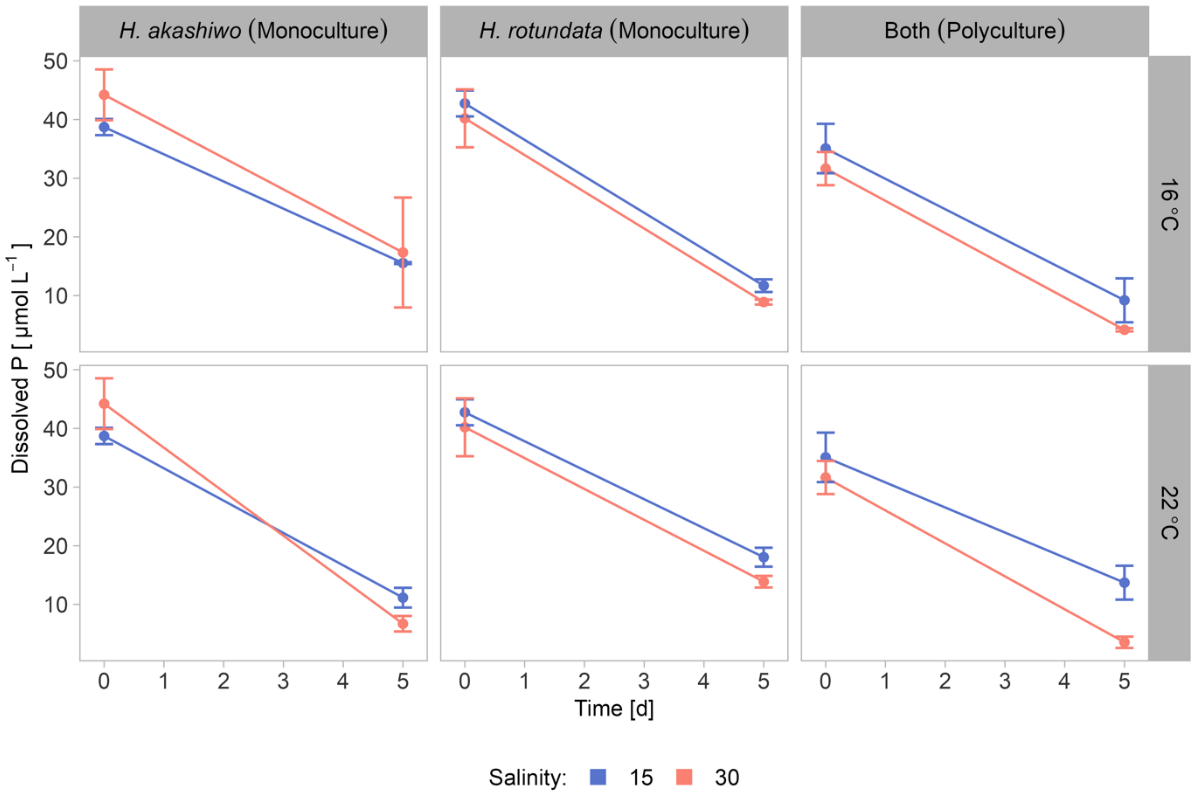


**Figure S7**: Dissolved phosphate concentrations [$\mu$mol L$^{-1}$] ± SD over time in the full-factorial nauplii grazing assay comprising two levels of temperature (facet rows) in combination with two levels of salinity (color) across the three prey compositions (*H. akashiwo* monoculture, *H. rotundata* monoculture as well as both species together in a polyculture).

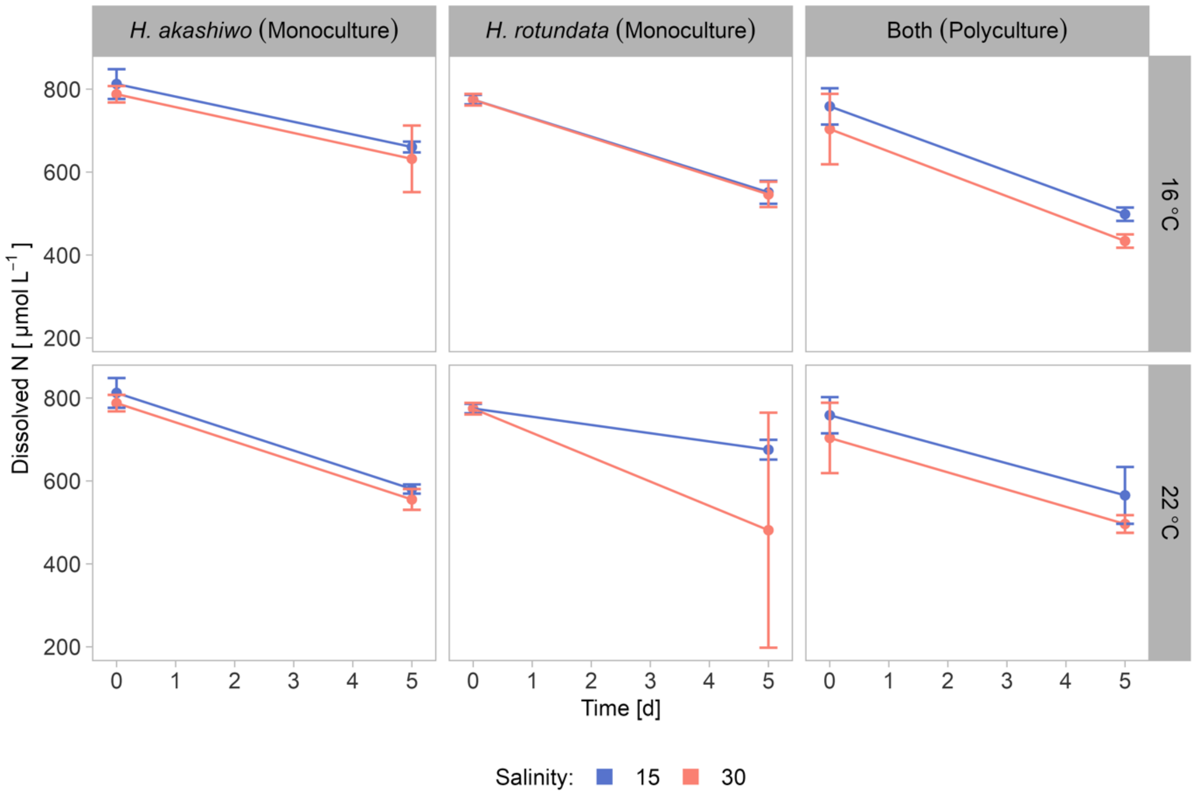


**Figure S8**: Dissolved nitrate and nitrite concentrations [$\mu$mol L$^{-1}$] ± SD over time in the full-factorial nauplii grazing assay comprising two levels of temperature (facet rows) in combination with two levels of salinity (color) across the three prey compositions (*H. akashiwo* monoculture, *H. rotundata* monoculture as well as both species together in a polyculture).

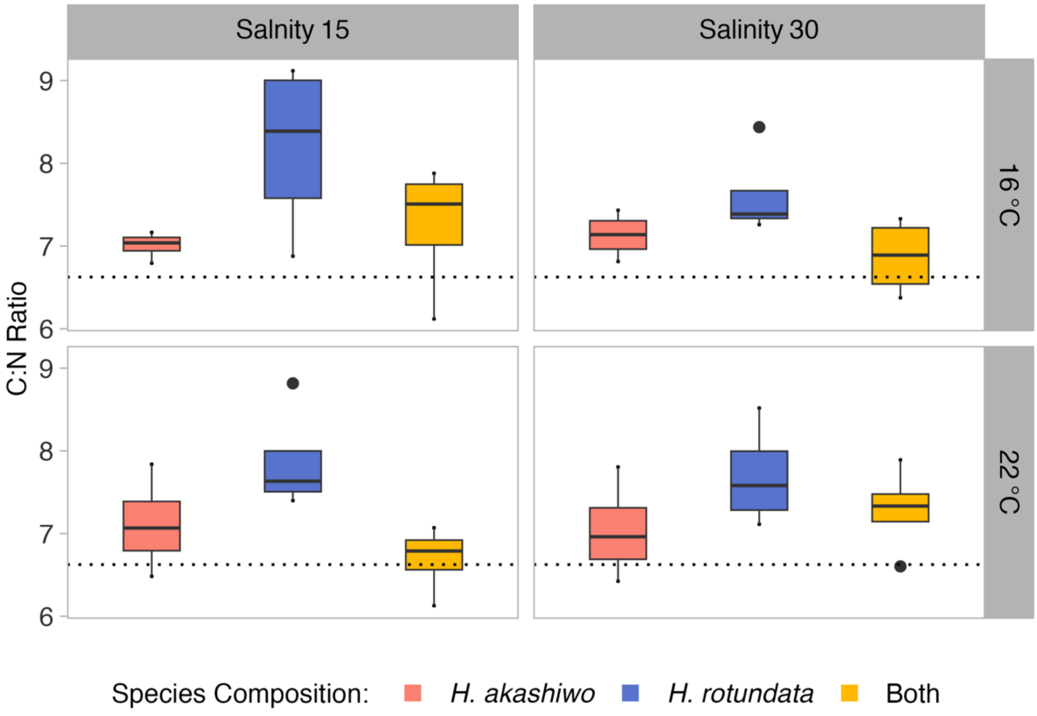


**Figure S9:** Particulate C:N ratio of *H. akashiwo*, *H. rotundata* and both species in polyculture across two levels of temperature (facet rows) in combination with two levels of salinity (facet columns) at the end of the full factorial *A. tonsa* grazing assay. The red field ratio is indicated by the dashed line.

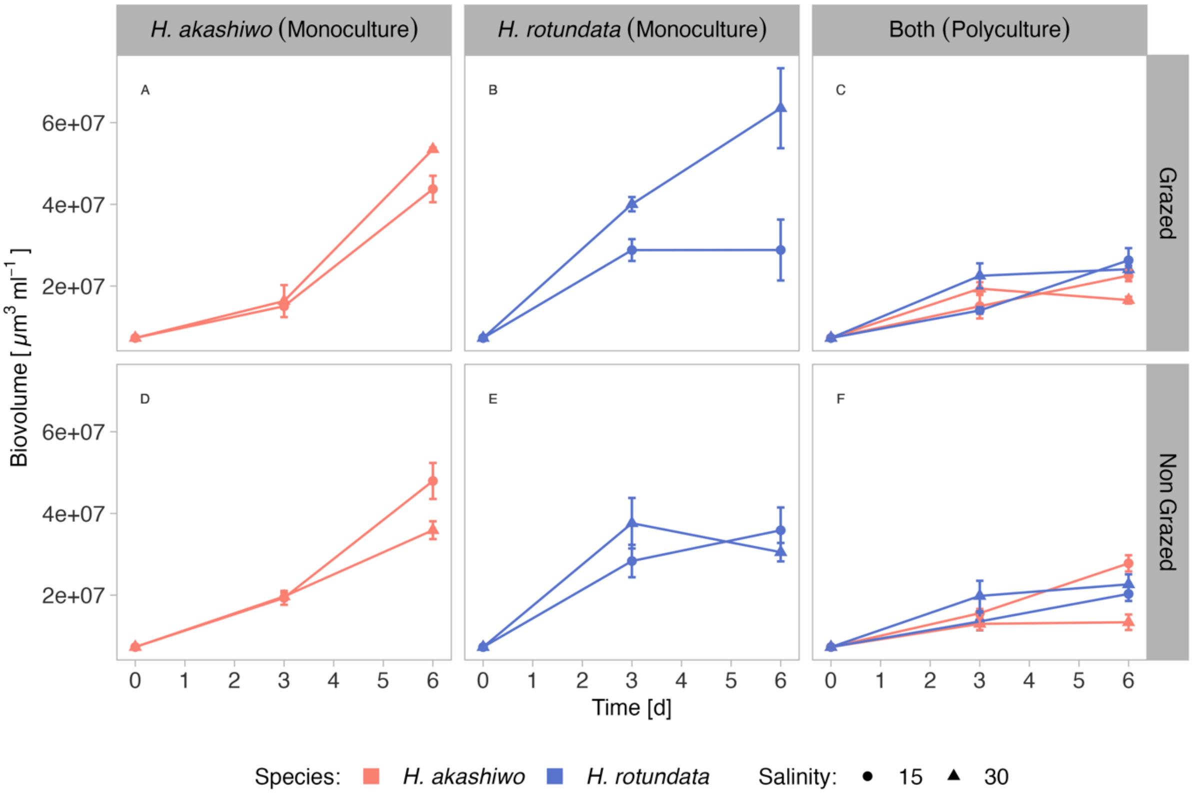


**Figure S10:** Mean biovolume ml $^{-1}$ of *H. akashiwo* and *H. rotundata* (indicated by color) over time ± SD incubated in monocultures and polycultures (facet columns) at two levels of salinity (indicated by shape) in rotifer grazed and non-grazed treatments represented by facet rows

**Table S1:** Post-hoc results for all significant main effects and interactions (response: maximum growth rate for *H. akashiwo* in competition experiment)

```
  Tukey multiple comparisons of means
    95% family-wise confidence level

Fit: aov(formula = r_max ~ salinity * temperature * competitor, data =
H.aka)

Tukey_rmax_H.aka$Salinity[Tukey_rmax_H.aka$Salinity[,4]<0.05,]
                          diff          lwr          upr        p adj
30-15               -0.1334375 -0.1578370844 -0.109037916 4.399470e-11

Tukey_rmax_H.aka$Temp[Tukey_rmax_H.aka$Temp[,4]<0.05,]
                          diff          lwr          upr        p adj
22-16                0.1149375  0.0905379156  0.139337084 8.495601e-10

Tukey_rmax_H.aka$Competitor[Tukey_rmax_H.aka$Competitor[,4]<0.05,]
                          diff          lwr          upr        p adj
yes-no              -0.0679375 -0.0923370844 -0.043537916 6.372610e-06

Tukey_rmax_H.aka$Salinity:Temp[Tukey_rmax_H.aka$Salinity:Temp[,4]<0.05,]
                          diff          lwr          upr        p adj
30:16-15:16         -0.1175000 -0.1636210546 -0.071378945 1.653388e-06
15:22-15:16          0.1308750  0.0847539454  0.176996055 2.669456e-07
15:22-30:16          0.2483750  0.2022539454  0.294496055 8.010259e-13
30:22-30:16          0.0990000  0.0528789454  0.145121055 2.332759e-05
30:22-15:22         -0.1493750 -0.1954960546 -0.103253945 2.456050e-08

Tukey_rmax_H.aka$Salinity:Competitor[Tukey_rmax_H.aka$Salinity:Competitor[,
4]<0.05,]
                          diff          lwr          upr        p adj
30:no-15:no         -0.1126250 -0.1587460546 -0.066503945 3.277793e-06
15:yes-15:no        -0.0471250 -0.0932460546 -0.001003945 4.395498e-02
30:yes-15:no        -0.2013750 -0.2474960546 -0.155253945 6.791201e-11
15:yes-30:no         0.0655000  0.0193789454  0.111621055 3.391323e-03
30:yes-30:no        -0.0887500 -0.1348710546 -0.042628945 1.061381e-04
30:yes-15:yes       -0.1542500 -0.2003710546 -0.108128945 1.344832e-08

Tukey_rmax_H.aka$Temp:Competitor[Tukey_rmax_H.aka$Temp:Competitor[,4]<0.05,
]
                          diff          lwr          upr        p adj
22:no-16:no          0.0961250  0.0500039454  0.142246055 3.558294e-05
16:yes-16:no        -0.0867500 -0.1328710546 -0.040628945 1.430207e-04
22:yes-16:no         0.0470000  0.0008789454  0.093121055 4.466966e-02
16:yes-22:no        -0.1828750 -0.2289960546 -0.136753945 4.848977e-10
22:yes-22:no        -0.0491250 -0.0952460546 -0.003003945 3.385139e-02
22:yes-16:yes        0.1337500  0.0876289454  0.179871055 1.823240e-07

Tukey_rmax_H.aka$Salinity:Temp:Competitor[Tukey_rmax_H.aka$Salinity:Temp:Co
mpetitor[,4]<0.05,]
                          diff          lwr          upr        p adj
15:22:no-15:16:no    0.1700000  0.0916925635  0.248307437 4.920187e-06
30:16:yes-15:16:no  -0.2042500 -0.2825574365 -0.125942563 2.015830e-07
15:22:yes-15:16:no   0.0837500  0.0054425635  0.162057437 3.012298e-02
15:22:no-30:16:no    0.2087500  0.1304425635  0.287057437 1.351066e-07
30:16:yes-30:16:no  -0.1655000 -0.2438074365 -0.087192563 7.630212e-06
15:22:yes-30:16:no   0.1225000  0.0441925635  0.200807437 5.984960e-04
30:22:no-15:22:no   -0.1865000 -0.2648074365 -0.108192563 1.021756e-06
15:16:yes-15:22:no  -0.1780000 -0.2563074365 -0.099692563 2.279229e-06
30:16:yes-15:22:no  -0.3742500 -0.4525574365 -0.295942563 9.195977e-13
15:22:yes-15:22:no  -0.0862500 -0.1645574365 -0.007942563 2.372406e-02
```

| | | | | |
|---|---|---|---|---|
| 30:22:yes-15:22:no | -0.1985000 | -0.2768074365 | -0.120192563 | 3.383799e-07 |
| 30:16:yes-30:22:no | -0.1877500 | -0.2660574365 | -0.109442563 | 9.092748e-07 |
| 15:22:yes-30:22:no | 0.1002500 | 0.0219425635 | 0.178557437 | 5.922041e-03 |
| 30:16:yes-15:16:yes | -0.1962500 | -0.2745574365 | -0.117942563 | 4.152560e-07 |
| 15:22:yes-15:16:yes | 0.0917500 | 0.0134425635 | 0.170057437 | 1.387635e-02 |
| 15:22:yes-30:16:yes | 0.2880000 | 0.2096925635 | 0.366307437 | 2.410184e-10 |
| 30:22:yes-30:16:yes | 0.1757500 | 0.0974425635 | 0.254057437 | 2.826030e-06 |
| 30:22:yes-15:22:yes | -0.1122500 | -0.1905574365 | -0.033942563 | 1.730205e-03 |

**Table S2:** Post-hoc results for all significant main effects and interactions (response: carrying capacity for *H. akashiwo* in the competition experiment)

```
  Tukey multiple comparisons of means
    95% family-wise confidence level

Fit: aov(formula = capacity ~ salinity * temperature * competitor, data =
H.aka)

Tukey_capacity_H.aka$Salinity[Tukey_capacity_H.aka$Salinity[,4]<0.05,]
                          diff          lwr        upr        p adj
30-15                -60368890  -72252742.4  -48485038 1.936291e-10

Tukey_capacity_H.aka$Temp[Tukey_capacity_H.aka$Temp[,4]<0.05,]
                          diff          lwr        upr        p adj
22-16                 39664072   27780219.7   51547924 4.003462e-07

Tukey_capacity_H.aka$Competitor[Tukey_capacity_H.aka$Competitor[,4]<0.05,]
                          diff          lwr        upr        p adj
yes-no               -56221014  -68104865.7  -44337161 7.819529e-10

Tukey_capacity_H.aka$Salinity:Temp[Tukey_capacity_H.aka$Salinity:Temp[,4]<0
.05,]
                          diff          lwr        upr        p adj
30:16-15:16          -85964747 -108428071.6  -63501422 9.854677e-10
15:22-30:16          100032962   77569637.3  122496287 4.510092e-11
30:22-30:16           65259928   42796603.4   87723253 1.766477e-07
30:22-15:22          -34773034  -57236358.7  -12309709 1.419641e-03

Tukey_capacity_H.aka$Salinity:Competitor[Tukey_capacity_H.aka$Salinity:Comp
etitor[,4]<0.05,]
                          diff          lwr        upr        p adj
30:no-15:no          -23470070  -45933394.5   -1006745 3.828486e-02
30:yes-15:no        -116589904 -139053228.7  -94126579 1.749045e-12
30:yes-30:no         -93119834 -115583159.1  -70656509 1.971419e-10
30:yes-15:yes        -97267711 -119731035.9  -74804386 8.064893e-11

Tukey_capacity_H.aka$Temp:Competitor[Tukey_capacity_H.aka$Temp:Competitor[,
4]<0.05,]

                          diff          lwr        upr        p adj
16:yes-16:no        -116797298 -139260622.6  -94333973 1.683986e-12
16:yes-22:no         -95885085 -118348410.2  -73421761 1.083152e-10
22:yes-16:yes        100240356   77777031.1  122703681 4.319678e-11

Tukey_capacity_H.aka$Salinity:Temp:Competitor[Tukey_capacity_H.aka$Salinity
:Temp:Competitor[,4]<0.05,]
                          diff          lwr        upr        p adj
30:22:no-15:16:no    -44382282  -82522029.7   -6242534 1.475910e-02
15:16:yes-15:16:no   -53991530  -92131277.7  -15851782 1.998534e-03
30:16:yes-15:16:no  -202762044 -240901792.2 -164622297 1.110223e-13
30:22:yes-15:16:no   -51018885  -89158632.7  -12879137 3.743375e-03
30:16:yes-30:16:no  -179603066 -217742813.2 -141463318 1.252776e-12
30:16:yes-15:22:no  -182160923 -220300670.4 -144021175 9.320322e-13
30:16:yes-30:22:no  -158379762 -196519510.2 -120240015 1.848266e-11
15:22:yes-30:22:no    39128305     988556.8   77268052 4.148509e-02
30:16:yes-15:16:yes -148770514 -186910262.2 -110630767 7.028456e-11
15:22:yes-15:16:yes   48737552   10597804.8   86877300 6.033235e-03
15:22:yes-30:16:yes  197508067  159368319.3  235647815 1.814104e-13
30:22:yes-30:16:yes  151743160  113603411.8  189882907 4.619027e-11
30:22:yes-15:22:yes  -45764907  -83904655.2   -7625160 1.114257e-02
```

**Table S3:** Post-hoc results for all significant main effects and interactions (response: maximum growth rate for *H. rotundata* in the competition experiment)

```
  Tukey multiple comparisons of means
    95% family-wise confidence level

Fit: aov(formula = r_max ~ salinity * temperature * competitor, data =
H.rot)

Tukey_rmax_H.rot$Salinity[Tukey_rmax_H.rot$Salinity[,4]<0.05,]
                          diff        lwr        upr        p adj
30-15                0.1781875  0.09663709  0.2597379 1.445299e-04

Tukey_rmax_H.rot$Competitor[Tukey_rmax_H.rot$Competitor[,4]<0.05,]
                          diff        lwr        upr        p adj
yes-no               0.1794375  0.09788709  0.2609879 1.333524e-04

Tukey_rmax_H.rot$Salinity:Temp[Tukey_rmax_H.rot$Salinity:Temp[,4]<0.05,]
                          diff        lwr        upr        p adj
15:22-15:16         -0.2015000 -0.35564980 -0.0473502 7.224830e-03
15:22-30:16         -0.2412500 -0.39539980 -0.0871002 1.262951e-03
30:22-15:22          0.3166250  0.16247520  0.4707748 4.369079e-05

Tukey_rmax_H.rot$Salinity:Competitor[Tukey_rmax_H.rot$Salinity:Competitor[,
4]<0.05,]
                          diff        lwr        upr        p adj
30:yes-15:no         0.3576250  0.20347520  0.5117748 7.307876e-06
30:yes-30:no         0.3113750  0.15722520  0.4655248 5.511292e-05
30:yes-15:yes        0.3101250  0.15597520  0.4642748 5.825153e-05

Tukey_rmax_H.rot$Salinity:Competitor[Tukey_rmax_H.rot$Salinity:Competitor[,
4]<0.05,]
                          diff        lwr        upr        p adj
16:yes-16:no         0.1732500  0.01910020  0.3273998 2.355402e-02
16:yes-22:no         0.2425000  0.08835020  0.3966498 1.194538e-03
22:yes-22:no         0.1856250  0.03147520  0.3397748 1.415372e-02

Tukey_rmax_H.rot$Salinity:Temp:Competitor[Tukey_rmax_H.rot$Salinity:Temp:Co
mpetitor[,4]<0.05,]
                          diff        lwr        upr        p adj
30:22:yes-15:16:no   0.3252500  0.06352407  0.5869759 7.967517e-03
30:16:yes-30:16:no   0.2745000  0.01277407  0.5362259 3.509891e-02
30:22:yes-30:16:no   0.3867500  0.12502407  0.6484759 1.209098e-03
30:16:yes-15:22:no   0.3900000  0.12827407  0.6517259 1.093261e-03
30:22:yes-15:22:no   0.5022500  0.24052407  0.7639759 3.468233e-05
30:22:yes-30:22:no   0.3482500  0.08652407  0.6099759 3.962258e-03
15:22:yes-30:16:yes -0.3670000 -0.62872593 -0.1052741 2.226868e-03
30:22:yes-15:22:yes  0.4792500  0.21752407  0.7409759 6.965698e-05
```

**Table S4:** Post-hoc results for all significant main effects and interactions (response: carrying capacity rate for *H. rotundata* in the competition experiment)

```
  Tukey multiple comparisons of means
    95% family-wise confidence level

Fit: aov(formula = capacity ~ salinity * temperature * competitor, data =
H.rot)

Tukey_capacity_H.rot$Salinity[Tukey_capacity_H.rot$Salinity[,4]<0.05,]
                          diff         lwr        upr        p adj
30-15                0.8068785  0.75577652  0.8579803 2.098322e-14

Tukey_capacity_H.rot$Temp[Tukey_capacity_H.rot$Temp[,4]<0.05,]
                          diff         lwr        upr        p adj
22-16               -0.5022071 -0.55330898 -0.4511051 2.120526e-14

Tukey_capacity_H.rot$Temp[Tukey_capacity_H.rot$Temp[,4]<0.05,]
                          diff         lwr        upr        p adj
yes-no              -0.1257134 -0.17681533 -0.0746114 3.417830e-05

Tukey_capacity_H.rot$Salinity:Temp[Tukey_capacity_H.rot$Salinity:Temp[,4]<0
.05,]
                          diff         lwr        upr        p adj
30:16-15:16          0.7454868  0.64889190  0.8420816 2.120526e-14
15:22-15:16         -0.5635987 -0.66019360 -0.4670038 1.609823e-13
30:22-15:16          0.3046714  0.20807652  0.4012662 4.009868e-08
15:22-30:16         -1.3090855 -1.40568038 -1.2124906 2.098322e-14
30:22-30:16         -0.4408154 -0.53741027 -0.3442205 2.703437e-11
30:22-15:22          0.8682701  0.77167524  0.9648649 2.109424e-14

Tukey_capacity_H.rot$Salinity:Competitor[Tukey_capacity_H.rot$Salinity:Comp
etitor[,4]<0.05,]
                          diff         lwr        upr        p adj
30:no-15:no          0.7356292  0.63903433  0.8322240 2.131628e-14
15:yes-15:no        -0.1969626 -0.29355752 -0.1003677 4.837610e-05
30:yes-15:no         0.6811651  0.58457017  0.7777599 2.298162e-14
15:yes-30:no        -0.9325918 -1.02918673 -0.8359969 2.098322e-14
30:yes-15:yes        0.8781277  0.78153281  0.9747225 2.098322e-14
22:no-16:no         -0.4859283 -0.58252315 -0.3893333 3.412159e-12

Tukey_capacity_H.rot$Temp:Competitor[Tukey_capacity_H.rot$Temp:Competitor[,
4]<0.05,]
                          diff         lwr        upr        p adj
16:yes-16:no        -0.1094346 -0.20602949 -0.0128397 2.226125e-02
22:yes-16:no        -0.6279205 -0.72451533 -0.5313255 3.419487e-14
16:yes-22:no         0.3764937  0.27989878  0.4730885 6.839616e-10
22:yes-22:no        -0.1419922 -0.23858706 -0.0453973 2.419227e-03
22:yes-16:yes       -0.5184858 -0.61508072 -0.4218909 8.584244e-13

Tukey_capacity_H.rot$Salinity:Temp:Competitor[Tukey_capacity_H.rot$Salinity
:Temp:Competitor[,4]<0.05,]
                          diff         lwr        upr        p adj
30:16:no-15:16:no    0.6905011  0.52649581  0.8545064 1.373968e-11
15:22:no-15:16:no   -0.5310564 -0.69506167 -0.3670510 3.210872e-09
30:22:no-15:16:no    0.2497009  0.08569564  0.4137062 8.404504e-04
15:16:yes-15:16:no  -0.1644203 -0.32842559 -0.0004149 4.910331e-02
30:16:yes-15:16:no   0.6360522  0.47204687  0.8000574 7.939471e-11
15:22:yes-15:16:no  -0.7605614 -0.92456666 -0.5965560 1.730727e-12
30:22:yes-15:16:no   0.1952216  0.03121626  0.3592268 1.200626e-02
15:22:no-30:16:no   -1.2215575 -1.38556278 -1.0575521 2.109424e-14
30:22:no-30:16:no   -0.4408002 -0.60480547 -0.2767948 1.161026e-07
15:16:yes-30:16:no  -0.8549214 -1.01892670 -0.6909160 1.598721e-13
```

| | | | | |
|---|---|---|---|---|
| 15:22:yes-30:16:no | -1.4510625 | -1.61506777 | -1.2870571 | 2.098322e-14 |
| 30:22:yes-30:16:no | -0.4952795 | -0.65928485 | -0.3312742 | 1.268358e-08 |
| 30:22:no-15:22:no | 0.7807573 | 0.61675201 | 0.9447626 | 9.977574e-13 |
| 15:16:yes-15:22:no | 0.3666361 | 0.20263078 | 0.5306413 | 3.020107e-06 |
| 30:16:yes-15:22:no | 1.1671085 | 1.00310324 | 1.3311138 | 2.109424e-14 |
| 15:22:yes-15:22:no | -0.2295050 | -0.39351029 | -0.0654996 | 2.278778e-03 |
| 30:22:yes-15:22:no | 0.7262779 | 0.56227263 | 0.8902832 | 4.631517e-12 |
| 15:16:yes-30:22:no | -0.4141212 | -0.57812653 | -0.2501159 | 3.628900e-07 |
| 30:16:yes-30:22:no | 0.3863512 | 0.22234593 | 0.5503565 | 1.235666e-06 |
| 15:22:yes-30:22:no | -1.0102623 | -1.17426760 | -0.8462569 | 2.364775e-14 |
| 30:16:yes-15:16:yes | 0.8004725 | 0.63646716 | 0.9644777 | 5.953016e-13 |
| 15:22:yes-15:16:yes | -0.5961411 | -0.76014637 | -0.4321357 | 3.073256e-10 |
| 30:22:yes-15:16:yes | 0.3596419 | 0.19563655 | 0.5236471 | 4.166024e-06 |
| 15:22:yes-30:16:yes | -1.3966135 | -1.56061883 | -1.2326082 | 2.098322e-14 |
| 30:22:yes-30:16:yes | -0.4408306 | -0.60483591 | -0.2768253 | 1.159542e-07 |
| 30:22:yes-15:22:yes | 0.9557829 | 0.79177762 | 1.1197882 | 3.153033e-14 |

**Table S5:** Post-hoc results for pairwise comparisons of *H. akashiwo* / *H. rotundata* biovolume log-ratios in the polycultures (competition experiment) between sampling days (data separated by salinity-temperature treatment combinations). Red marked values represent log-ratio comparisons against day zero (used in the text).

Pairwise comparisons using Wilcoxon rank sum exact test

data: s30t16$log_ratio and s30t16$day

| | 0 | 2 | 4 | 6 | 8 | 10 | 12 | 14 | 16 | 18 | 20 | 22 | 24 | 26 | 28 |
|---|---|---|---|---|---|---|---|---|---|---|---|---|---|---|---|
| 2 | 0.114 | - | - | - | - | - | - | - | - | - | - | - | - | - | - |
| 4 | 0.029 | 0.029 | - | - | - | - | - | - | - | - | - | - | - | - | - |
| 6 | 0.029 | 0.029 | 0.486 | - | - | - | - | - | - | - | - | - | - | - | - |
| 8 | 0.029 | 0.029 | 0.200 | 0.686 | - | - | - | - | - | - | - | - | - | - | - |
| 10 | 0.029 | 0.029 | 0.200 | 0.057 | 0.200 | - | - | - | - | - | - | - | - | - | - |
| 12 | 0.029 | 0.029 | 0.343 | 0.486 | 0.886 | 0.343 | - | - | - | - | - | - | - | - | - |
| 14 | 0.029 | 0.029 | 0.886 | 0.886 | 0.686 | 0.200 | 0.486 | - | - | - | - | - | - | - | - |
| 16 | 0.029 | 0.114 | 0.886 | 0.886 | 1.000 | 0.057 | 0.686 | 0.686 | - | - | - | - | - | - | - |
| 18 | 0.029 | 0.343 | 0.029 | 0.029 | 0.029 | 0.029 | 0.029 | 0.029 | 0.343 | - | - | - | - | - | - |
| 20 | 0.686 | 0.029 | 0.029 | 0.029 | 0.029 | 0.029 | 0.029 | 0.029 | 0.029 | 0.029 | - | - | - | - | - |
| 22 | 0.029 | 0.029 | 0.029 | 0.029 | 0.029 | 0.029 | 0.029 | 0.029 | 0.029 | 0.029 | 0.200 | - | - | - | - |
| 24 | 0.029 | 0.029 | 0.029 | 0.029 | 0.029 | 0.029 | 0.029 | 0.029 | 0.029 | 0.029 | 0.029 | 0.057 | - | - | - |
| 26 | 0.029 | 0.029 | 0.029 | 0.029 | 0.029 | 0.029 | 0.029 | 0.029 | 0.029 | 0.029 | 0.029 | 0.029 | 0.686 | - | - |
| 28 | 0.029 | 0.029 | 0.029 | 0.029 | 0.029 | 0.029 | 0.029 | 0.029 | 0.029 | 0.029 | 0.029 | 0.029 | 0.886 | 1.000 | - |

data: s15t16$log_ratio and s15t16$day

| | 0 | 2 | 4 | 6 | 8 | 10 | 12 | 14 | 16 | 18 | 20 | 22 | 24 | 26 | 28 |
|---|---|---|---|---|---|---|---|---|---|---|---|---|---|---|---|
| 2 | 0.686 | - | - | - | - | - | - | - | - | - | - | - | - | - | - |
| 4 | 0.029 | 0.029 | - | - | - | - | - | - | - | - | - | - | - | - | - |
| 6 | 0.029 | 0.029 | 0.486 | - | - | - | - | - | - | - | - | - | - | - | - |
| 8 | 0.114 | 0.114 | 0.886 | 0.486 | - | - | - | - | - | - | - | - | - | - | - |
| 10 | 0.114 | 0.343 | 0.029 | 0.029 | 0.029 | - | - | - | - | - | - | - | - | - | - |
| 12 | 0.029 | 0.029 | 0.029 | 0.029 | 0.029 | 0.029 | - | - | - | - | - | - | - | - | - |
| 14 | 0.029 | 0.029 | 0.029 | 0.029 | 0.029 | 0.029 | 0.029 | - | - | - | - | - | - | - | - |
| 16 | 0.029 | 0.029 | 0.029 | 0.029 | 0.029 | 0.029 | 0.029 | 0.029 | - | - | - | - | - | - | - |
| 18 | 0.029 | 0.029 | 0.029 | 0.029 | 0.029 | 0.029 | 0.029 | 0.029 | 0.886 | - | - | - | - | - | - |
| 20 | 0.029 | 0.029 | 0.029 | 0.029 | 0.029 | 0.029 | 0.029 | 0.029 | 0.486 | 0.057 | - | - | - | - | - |
| 22 | 0.029 | 0.029 | 0.029 | 0.029 | 0.029 | 0.029 | 0.029 | 0.029 | 0.886 | 0.343 | 0.114 | - | - | - | - |
| 24 | 0.029 | 0.029 | 0.029 | 0.029 | 0.029 | 0.029 | 0.029 | 0.029 | 0.686 | 0.686 | 0.029 | 0.200 | - | - | - |
| 26 | 0.029 | 0.029 | 0.029 | 0.029 | 0.029 | 0.029 | 0.029 | 0.029 | 0.486 | 0.686 | 0.114 | 0.486 | 0.886 | - | - |
| 28 | 0.029 | 0.029 | 0.029 | 0.029 | 0.029 | 0.029 | 0.029 | 0.029 | 0.029 | 0.029 | 0.029 | 0.029 | 0.343 | 0.486 | - |

data: s30t22$log_ratio and s30t22$day

| | 0 | 2 | 4 | 6 | 8 | 10 | 12 | 14 | 16 | 18 | 20 | 22 | 24 | 26 | 28 |
|---|---|---|---|---|---|---|---|---|---|---|---|---|---|---|---|
| 2 | 0.886 | - | - | - | - | - | - | - | - | - | - | - | - | - | - |
| 4 | 0.029 | 0.029 | - | - | - | - | - | - | - | - | - | - | - | - | - |
| 6 | 0.114 | 0.029 | 0.191 | - | - | - | - | - | - | - | - | - | - | - | - |
| 8 | 1.000 | 0.486 | 0.059 | 0.200 | - | - | - | - | - | - | - | - | - | - | - |
| 10 | 0.343 | 0.114 | 0.029 | 0.029 | 0.114 | - | - | - | - | - | - | - | - | - | - |
| 12 | 0.029 | 0.029 | 0.029 | 0.029 | 0.029 | 0.029 | - | - | - | - | - | - | - | - | - |
| 14 | 0.029 | 0.029 | 0.029 | 0.029 | 0.029 | 0.029 | 0.029 | - | - | - | - | - | - | - | - |
| 16 | 0.029 | 0.029 | 0.029 | 0.029 | 0.029 | 0.029 | 0.029 | 0.029 | - | - | - | - | - | - | - |
| 18 | 0.029 | 0.029 | 0.029 | 0.029 | 0.029 | 0.029 | 0.029 | 0.029 | 0.029 | - | - | - | - | - | - |
| 20 | 0.029 | 0.029 | 0.029 | 0.029 | 0.029 | 0.029 | 0.029 | 0.029 | 0.029 | 0.029 | - | - | - | - | - |
| 22 | 0.029 | 0.029 | 0.029 | 0.029 | 0.029 | 0.029 | 0.029 | 0.029 | 0.029 | 0.029 | 0.886 | - | - | - | - |
| 24 | 0.029 | 0.029 | 0.029 | 0.029 | 0.029 | 0.029 | 0.029 | 0.029 | 0.029 | 0.343 | 0.029 | 0.029 | - | - | - |
| 26 | 0.029 | 0.029 | 0.029 | 0.029 | 0.029 | 0.029 | 0.029 | 0.029 | 0.029 | 0.029 | 0.343 | 0.486 | 0.029 | - | - |
| 28 | 0.029 | 0.029 | 0.029 | 0.029 | 0.029 | 0.029 | 0.029 | 0.029 | 0.029 | 0.029 | 0.029 | 0.029 | 0.029 | 0.200 | - |

data: s15t22$log_ratio and s15t22$day

| | 0 | 2 | 4 | 6 | 8 | 10 | 12 | 14 | 16 | 18 | 20 | 22 | 24 | 26 | 28 |
|---|---|---|---|---|---|---|---|---|---|---|---|---|---|---|---|
| 2 | 0.029 | - | - | - | - | - | - | - | - | - | - | - | - | - | - |
| 4 | 0.029 | 0.663 | - | - | - | - | - | - | - | - | - | - | - | - | - |
| 6 | 0.029 | 0.057 | 0.029 | - | - | - | - | - | - | - | - | - | - | - | - |
| 8 | 0.029 | 0.029 | 0.029 | 0.029 | - | - | - | - | - | - | - | - | - | - | - |
| 10 | 0.029 | 0.029 | 0.029 | 0.029 | 0.029 | - | - | - | - | - | - | - | - | - | - |
| 12 | 0.029 | 0.029 | 0.029 | 0.029 | 0.029 | 0.029 | - | - | - | - | - | - | - | - | - |
| 14 | 0.029 | 0.029 | 0.028 | 0.029 | 0.029 | 0.029 | 0.029 | - | - | - | - | - | - | - | - |
| 16 | 0.029 | 0.029 | 0.029 | 0.029 | 0.029 | 0.029 | 0.029 | 0.110 | - | - | - | - | - | - | - |
| 18 | 0.029 | 0.029 | 0.029 | 0.029 | 0.029 | 0.029 | 0.029 | 0.029 | 0.886 | - | - | - | - | - | - |
| 20 | 0.029 | 0.029 | 0.029 | 0.029 | 0.029 | 0.029 | 0.029 | 0.029 | 0.343 | 0.686 | - | - | - | - | - |
| 22 | 0.029 | 0.029 | 0.029 | 0.029 | 0.029 | 0.029 | 0.029 | 0.029 | 0.029 | 0.114 | 0.200 | - | - | - | - |
| 24 | 0.029 | 0.029 | 0.029 | 0.029 | 0.029 | 0.029 | 0.029 | 0.029 | 0.029 | 0.343 | 0.686 | 0.343 | - | - | - |
| 26 | 0.029 | 0.029 | 0.029 | 0.029 | 0.029 | 0.029 | 0.029 | 0.029 | 0.686 | 0.200 | 0.146 | 0.029 | 0.029 | - | - |
| 28 | 0.029 | 0.029 | 0.029 | 0.029 | 0.029 | 0.029 | 0.029 | 0.110 | 0.561 | 0.686 | 0.343 | 0.029 | 0.029 | 0.886 | - |

**Table S6:** ANOVA results testing the effects of temperature, salinity and species composition on the particulate organic C:N ratios at the end of the competition experiment. Degrees of freedom (Df), F and p-values are given for each effect. Values marked with an asterisk (*) indicate significant effects ($p < 0.05$).

| | particulate C:N ratio | | | |
|---|---|---|---|---|
| Effect | *Df* | *F* | *p* | |
| Species Composition | 2 | 19.2 | <0.001 | * |
| Temperature | 1 | 1.2 | 0.292 | |
| Salinity | 1 | <0.1 | 0.951 | |
| Species Composition × Temperature | 2 | 11.2 | <0.001 | * |
| Species Composition × Salinity | 2 | 4.5 | 0.021 | * |
| Temperature × Salinity | 1 | <0.1 | 0.901 | |
| Species Composition × Temperature × Salinity | 2 | 10.3 | <0.001 | * |

**Table S7:** Post-hoc results for all significant pairwise comparisons (response: *A. tonsa* nauplii mortality in the full-factorial grazing assay across three different prey composition treatments, two levels of salinity, two levels of temperature and 5 sampling days).

```
emmeans(mortality, pairwise ~ day * prey * sal * temp)
$emmeans

Degrees-of-freedom method: kenward-roger
P value adjustment: tukey method for comparing a family of 72 estimates
```

| contrast | estimate | SE | df | t.ratio | p.value |
|---|---|---|---|---|---|
| day0 both sal15 temp16 - day5 h.aka sal15 temp16 | -64,881 | 7,85863 | 213,4551 | -8,25601 | 4,07E-11 |
| day0 both sal15 temp16 - day5 h.aka sal30 temp16 | -41,6667 | 7,85863 | 213,4551 | -5,30203 | 0,000624 |
| day0 both sal15 temp16 - day2 h.aka sal15 temp22 | -54,5833 | 7,85863 | 213,4551 | -6,94566 | 1,13E-07 |
| day0 both sal15 temp16 - day3 h.aka sal15 temp22 | -90 | 7,85863 | 213,4551 | -11,4524 | 2,87E-13 |
| day0 both sal15 temp16 - day4 h.aka sal15 temp22 | -100 | 7,85863 | 213,4551 | -12,7249 | 9,56E-14 |
| day0 both sal15 temp16 - day5 h.aka sal15 temp22 | -93,75 | 7,85863 | 213,4551 | -11,9296 | 1,43E-13 |
| day0 both sal15 temp16 - day3 h.aka sal30 temp22 | -68,75 | 7,85863 | 213,4551 | -8,74834 | 2,44E-12 |
| day0 both sal15 temp16 - day4 h.aka sal30 temp22 | -66,6667 | 7,85863 | 213,4551 | -8,48324 | 1,01E-11 |
| day0 both sal15 temp16 - day5 h.aka sal30 temp22 | -91,6667 | 7,85863 | 213,4551 | -11,6645 | 2,1E-13 |
| day1 both sal15 temp16 - day5 h.aka sal15 temp16 | -64,881 | 7,85863 | 213,4551 | -8,25601 | 4,07E-11 |
| day1 both sal15 temp16 - day5 h.aka sal30 temp16 | -41,6667 | 7,85863 | 213,4551 | -5,30203 | 0,000624 |
| day1 both sal15 temp16 - day2 h.aka sal15 temp22 | -54,5833 | 7,85863 | 213,4551 | -6,94566 | 1,13E-07 |
| day1 both sal15 temp16 - day3 h.aka sal15 temp22 | -90 | 7,85863 | 213,4551 | -11,4524 | 2,87E-13 |
| day1 both sal15 temp16 - day4 h.aka sal15 temp22 | -100 | 7,85863 | 213,4551 | -12,7249 | 9,56E-14 |
| day1 both sal15 temp16 - day5 h.aka sal15 temp22 | -93,75 | 7,85863 | 213,4551 | -11,9296 | 1,43E-13 |
| day1 both sal15 temp16 - day3 h.aka sal30 temp22 | -68,75 | 7,85863 | 213,4551 | -8,74834 | 2,44E-12 |
| day1 both sal15 temp16 - day4 h.aka sal30 temp22 | -66,6667 | 7,85863 | 213,4551 | -8,48324 | 1,01E-11 |
| day1 both sal15 temp16 - day5 h.aka sal30 temp22 | -91,6667 | 7,85863 | 213,4551 | -11,6645 | 2,1E-13 |
| day2 both sal15 temp16 - day5 h.aka sal15 temp16 | -64,881 | 7,85863 | 213,4551 | -8,25601 | 4,07E-11 |
| day2 both sal15 temp16 - day5 h.aka sal30 temp16 | -41,6667 | 7,85863 | 213,4551 | -5,30203 | 0,000624 |
| day2 both sal15 temp16 - day2 h.aka sal15 temp22 | -54,5833 | 7,85863 | 213,4551 | -6,94566 | 1,13E-07 |
| day2 both sal15 temp16 - day3 h.aka sal15 temp22 | -90 | 7,85863 | 213,4551 | -11,4524 | 2,87E-13 |
| day2 both sal15 temp16 - day4 h.aka sal15 temp22 | -100 | 7,85863 | 213,4551 | -12,7249 | 9,56E-14 |
| day2 both sal15 temp16 - day5 h.aka sal15 temp22 | -93,75 | 7,85863 | 213,4551 | -11,9296 | 1,43E-13 |
| day2 both sal15 temp16 - day3 h.aka sal30 temp22 | -68,75 | 7,85863 | 213,4551 | -8,74834 | 2,44E-12 |
| day2 both sal15 temp16 - day4 h.aka sal30 temp22 | -66,6667 | 7,85863 | 213,4551 | -8,48324 | 1,01E-11 |
| day2 both sal15 temp16 - day5 h.aka sal30 temp22 | -91,6667 | 7,85863 | 213,4551 | -11,6645 | 2,1E-13 |
| day3 both sal15 temp16 - day5 h.aka sal15 temp16 | -56,5476 | 7,85863 | 213,4551 | -7,19561 | 2,63E-08 |
| day3 both sal15 temp16 - day5 h.aka sal30 temp16 | -33,3333 | 7,85863 | 213,4551 | -4,24162 | 0,047643 |
| day3 both sal15 temp16 - day2 h.aka sal15 temp22 | -46,25 | 7,85863 | 213,4551 | -5,88525 | 3,62E-05 |
| day3 both sal15 temp16 - day3 h.aka sal15 temp22 | -81,6667 | 7,85863 | 213,4551 | -10,392 | 7,11E-13 |
| day3 both sal15 temp16 - day4 h.aka sal15 temp22 | -91,6667 | 7,85863 | 213,4551 | -11,6645 | 2,1E-13 |
| day3 both sal15 temp16 - day5 h.aka sal15 temp22 | -85,4167 | 7,85863 | 213,4551 | -10,8692 | 5,63E-13 |
| day3 both sal15 temp16 - day3 h.aka sal30 temp22 | -60,4167 | 7,85863 | 213,4551 | -7,68794 | 1,37E-09 |
| day3 both sal15 temp16 - day4 h.aka sal30 temp22 | -58,3333 | 7,85863 | 213,4551 | -7,42284 | 6,82E-09 |
| day3 both sal15 temp16 - day5 h.aka sal30 temp22 | -83,3333 | 7,85863 | 213,4551 | -10,6041 | 6,96E-13 |
| day4 both sal15 temp16 - day5 h.aka sal15 temp16 | -59,881 | 7,85863 | 213,4551 | -7,61977 | 2,08E-09 |
| day4 both sal15 temp16 - day5 h.aka sal30 temp16 | -36,6667 | 7,85863 | 213,4551 | -4,66578 | 0,009819 |
| day4 both sal15 temp16 - day2 h.aka sal15 temp22 | -49,5833 | 7,85863 | 213,4551 | -6,30941 | 3,91E-06 |
| day4 both sal15 temp16 - day3 h.aka sal15 temp22 | -85 | 7,85863 | 213,4551 | -10,8161 | 5,98E-13 |
| day4 both sal15 temp16 - day4 h.aka sal15 temp22 | -95 | 7,85863 | 213,4551 | -12,0886 | 1,33E-13 |
| day4 both sal15 temp16 - day5 h.aka sal15 temp22 | -88,75 | 7,85863 | 213,4551 | -11,2933 | 3,51E-13 |
| day4 both sal15 temp16 - day3 h.aka sal30 temp22 | -63,75 | 7,85863 | 213,4551 | -8,1121 | 9,97E-11 |
| day4 both sal15 temp16 - day4 h.aka sal30 temp22 | -61,6667 | 7,85863 | 213,4551 | -7,847 | 5,17E-10 |
| day4 both sal15 temp16 - day5 h.aka sal30 temp22 | -86,6667 | 7,85863 | 213,4551 | -11,0282 | 4,67E-13 |
| day5 both sal15 temp16 - day5 h.aka sal15 temp16 | -56,5476 | 7,85863 | 213,4551 | -7,19561 | 2,63E-08 |
| day5 both sal15 temp16 - day5 h.aka sal30 temp16 | -33,3333 | 7,85863 | 213,4551 | -4,24162 | 0,047643 |
| day5 both sal15 temp16 - day2 h.aka sal15 temp22 | -46,25 | 7,85863 | 213,4551 | -5,88525 | 3,62E-05 |
| day5 both sal15 temp16 - day3 h.aka sal15 temp22 | -81,6667 | 7,85863 | 213,4551 | -10,392 | 7,11E-13 |
| day5 both sal15 temp16 - day4 h.aka sal15 temp22 | -91,6667 | 7,85863 | 213,4551 | -11,6645 | 2,1E-13 |
| day5 both sal15 temp16 - day5 h.aka sal15 temp22 | -85,4167 | 7,85863 | 213,4551 | -10,8692 | 5,63E-13 |
| day5 both sal15 temp16 - day3 h.aka sal30 temp22 | -60,4167 | 7,85863 | 213,4551 | -7,68794 | 1,37E-09 |
| day5 both sal15 temp16 - day4 h.aka sal30 temp22 | -58,3333 | 7,85863 | 213,4551 | -7,42284 | 6,82E-09 |
| day5 both sal15 temp16 - day5 h.aka sal30 temp22 | -83,3333 | 7,85863 | 213,4551 | -10,6041 | 6,96E-13 |
| day0 h.aka sal15 temp16 - day5 h.aka sal15 temp16 | -64,881 | 7,664356 | 180 | -8,46528 | 2,29E-11 |
| day0 h.aka sal15 temp16 - day5 h.aka sal30 temp16 | -41,6667 | 7,85863 | 213,4551 | -5,30203 | 0,000624 |
| day0 h.aka sal15 temp16 - day2 h.aka sal15 temp22 | -54,5833 | 7,85863 | 213,4551 | -6,94566 | 1,13E-07 |
| day0 h.aka sal15 temp16 - day3 h.aka sal15 temp22 | -90 | 7,85863 | 213,4551 | -11,4524 | 2,87E-13 |
| day0 h.aka sal15 temp16 - day4 h.aka sal15 temp22 | -100 | 7,85863 | 213,4551 | -12,7249 | 9,56E-14 |
| day0 h.aka sal15 temp16 - day5 h.aka sal15 temp22 | -93,75 | 7,85863 | 213,4551 | -11,9296 | 1,43E-13 |
| day0 h.aka sal15 temp16 - day3 h.aka sal30 temp22 | -68,75 | 7,85863 | 213,4551 | -8,74834 | 2,44E-12 |
| day0 h.aka sal15 temp16 - day4 h.aka sal30 temp22 | -66,6667 | 7,85863 | 213,4551 | -8,48324 | 1,01E-11 |
| day0 h.aka sal15 temp16 - day5 h.aka sal30 temp22 | -91,6667 | 7,85863 | 213,4551 | -11,6645 | 2,1E-13 |
| day1 h.aka sal15 temp16 - day5 h.aka sal15 temp16 | -64,881 | 7,664356 | 180 | -8,46528 | 2,29E-11 |
| day1 h.aka sal15 temp16 - day5 h.aka sal30 temp16 | -41,6667 | 7,85863 | 213,4551 | -5,30203 | 0,000624 |
| day1 h.aka sal15 temp16 - day2 h.aka sal15 temp22 | -54,5833 | 7,85863 | 213,4551 | -6,94566 | 1,13E-07 |
| day1 h.aka sal15 temp16 - day3 h.aka sal15 temp22 | -90 | 7,85863 | 213,4551 | -11,4524 | 2,87E-13 |
| day1 h.aka sal15 temp16 - day4 h.aka sal15 temp22 | -100 | 7,85863 | 213,4551 | -12,7249 | 9,56E-14 |
| day1 h.aka sal15 temp16 - day5 h.aka sal15 temp22 | -93,75 | 7,85863 | 213,4551 | -11,9296 | 1,43E-13 |

| | | | | | |
|---|---|---|---|---|---|
| day1 h.aka sal15 temp16 - day3 h.aka sal30 temp22 | -68,75 | 7,85863 | 213,4551 | -8,74834 | 2,44E-12 |
| day1 h.aka sal15 temp16 - day4 h.aka sal30 temp22 | -66,6667 | 7,85863 | 213,4551 | -8,48324 | 1,01E-11 |
| day1 h.aka sal15 temp16 - day5 h.aka sal30 temp22 | -91,6667 | 7,85863 | 213,4551 | -11,6645 | 2,1E-13 |
| day2 h.aka sal15 temp16 - day5 h.aka sal15 temp16 | -64,881 | 7,664356 | 180 | -8,46528 | 2,29E-11 |
| day2 h.aka sal15 temp16 - day5 h.aka sal30 temp16 | -41,6667 | 7,85863 | 213,4551 | -5,30203 | 0,000624 |
| day2 h.aka sal15 temp16 - day2 h.aka sal15 temp22 | -54,5833 | 7,85863 | 213,4551 | -6,94566 | 1,13E-07 |
| day2 h.aka sal15 temp16 - day3 h.aka sal15 temp22 | -90 | 7,85863 | 213,4551 | -11,4524 | 2,87E-13 |
| day2 h.aka sal15 temp16 - day4 h.aka sal15 temp22 | -100 | 7,85863 | 213,4551 | -12,7249 | 9,56E-14 |
| day2 h.aka sal15 temp16 - day5 h.aka sal15 temp22 | -93,75 | 7,85863 | 213,4551 | -11,9296 | 1,43E-13 |
| day2 h.aka sal15 temp16 - day3 h.aka sal30 temp22 | -68,75 | 7,85863 | 213,4551 | -8,74834 | 2,44E-12 |
| day2 h.aka sal15 temp16 - day4 h.aka sal30 temp22 | -66,6667 | 7,85863 | 213,4551 | -8,48324 | 1,01E-11 |
| day2 h.aka sal15 temp16 - day5 h.aka sal30 temp22 | -91,6667 | 7,85863 | 213,4551 | -11,6645 | 2,1E-13 |
| day3 h.aka sal15 temp16 - day5 h.aka sal15 temp16 | -41,5476 | 7,664356 | 180 | -5,42089 | 0,000416 |
| day3 h.aka sal15 temp16 - day3 h.aka sal15 temp22 | -66,6667 | 7,85863 | 213,4551 | -8,48324 | 1,01E-11 |
| day3 h.aka sal15 temp16 - day4 h.aka sal15 temp22 | -76,6667 | 7,85863 | 213,4551 | -9,75573 | 8E-13 |
| day3 h.aka sal15 temp16 - day5 h.aka sal15 temp22 | -70,4167 | 7,85863 | 213,4551 | -8,96043 | 1,18E-12 |
| day3 h.aka sal15 temp16 - day3 h.aka sal30 temp22 | -45,4167 | 7,85863 | 213,4551 | -5,77921 | 6,2E-05 |
| day3 h.aka sal15 temp16 - day4 h.aka sal30 temp22 | -43,3333 | 7,85863 | 213,4551 | -5,51411 | 0,000229 |
| day3 h.aka sal15 temp16 - day5 h.aka sal30 temp22 | -68,3333 | 7,85863 | 213,4551 | -8,69532 | 3,15E-12 |
| day4 h.aka sal15 temp16 - day5 h.aka sal15 temp16 | -33,3333 | 7,664356 | 180 | -4,34914 | 0,034389 |
| day4 h.aka sal15 temp16 - day3 h.aka sal15 temp22 | -58,4524 | 7,85863 | 213,4551 | -7,43799 | 6,23E-09 |
| day4 h.aka sal15 temp16 - day4 h.aka sal15 temp22 | -68,4524 | 7,85863 | 213,4551 | -8,71047 | 2,92E-12 |
| day4 h.aka sal15 temp16 - day5 h.aka sal15 temp22 | -62,2024 | 7,85863 | 213,4551 | -7,91517 | 3,39E-10 |
| day4 h.aka sal15 temp16 - day3 h.aka sal30 temp22 | -37,2024 | 7,85863 | 213,4551 | -4,73395 | 0,007459 |
| day4 h.aka sal15 temp16 - day4 h.aka sal30 temp22 | -35,119 | 7,85863 | 213,4551 | -4,46885 | 0,021053 |
| day4 h.aka sal15 temp16 - day5 h.aka sal30 temp22 | -60,119 | 7,85863 | 213,4551 | -7,65007 | 1,73E-09 |
| day5 h.aka sal15 temp16 - day0 h.rot sal15 temp16 | 64,88095 | 7,85863 | 213,4551 | 8,256013 | 4,07E-11 |
| day5 h.aka sal15 temp16 - day1 h.rot sal15 temp16 | 56,54762 | 7,85863 | 213,4551 | 7,195608 | 2,63E-08 |
| day5 h.aka sal15 temp16 - day2 h.rot sal15 temp16 | 64,88095 | 7,85863 | 213,4551 | 8,256013 | 4,07E-11 |
| day5 h.aka sal15 temp16 - day3 h.rot sal15 temp16 | 64,88095 | 7,85863 | 213,4551 | 8,256013 | 4,07E-11 |
| day5 h.aka sal15 temp16 - day4 h.rot sal15 temp16 | 56,54762 | 7,85863 | 213,4551 | 7,195608 | 2,63E-08 |
| day5 h.aka sal15 temp16 - day5 h.rot sal15 temp16 | 55,71429 | 7,85863 | 213,4551 | 7,089568 | 4,89E-08 |
| day5 h.aka sal15 temp16 - day0 both sal30 temp16 | 64,88095 | 7,85863 | 213,4551 | 8,256013 | 4,07E-11 |
| day5 h.aka sal15 temp16 - day1 both sal30 temp16 | 64,88095 | 7,85863 | 213,4551 | 8,256013 | 4,07E-11 |
| day5 h.aka sal15 temp16 - day2 both sal30 temp16 | 64,88095 | 7,85863 | 213,4551 | 8,256013 | 4,07E-11 |
| day5 h.aka sal15 temp16 - day3 both sal30 temp16 | 64,88095 | 7,85863 | 213,4551 | 8,256013 | 4,07E-11 |
| day5 h.aka sal15 temp16 - day4 both sal30 temp16 | 56,54762 | 7,85863 | 213,4551 | 7,195608 | 2,63E-08 |
| day5 h.aka sal15 temp16 - day5 both sal30 temp16 | 58,63095 | 7,85863 | 213,4551 | 7,460709 | 5,44E-09 |
| day5 h.aka sal15 temp16 - day0 h.aka sal30 temp16 | 64,88095 | 7,85863 | 213,4551 | 8,256013 | 4,07E-11 |
| day5 h.aka sal15 temp16 - day1 h.aka sal30 temp16 | 64,88095 | 7,85863 | 213,4551 | 8,256013 | 4,07E-11 |
| day5 h.aka sal15 temp16 - day2 h.aka sal30 temp16 | 64,88095 | 7,85863 | 213,4551 | 8,256013 | 4,07E-11 |
| day5 h.aka sal15 temp16 - day3 h.aka sal30 temp16 | 53,42262 | 7,85863 | 213,4551 | 6,797956 | 2,62E-07 |
| day5 h.aka sal15 temp16 - day4 h.aka sal30 temp16 | 61,30952 | 7,85863 | 213,4551 | 7,801554 | 6,84E-10 |
| day5 h.aka sal15 temp16 - day0 h.rot sal30 temp16 | 64,88095 | 7,85863 | 213,4551 | 8,256013 | 4,07E-11 |
| day5 h.aka sal15 temp16 - day1 h.rot sal30 temp16 | 64,88095 | 7,85863 | 213,4551 | 8,256013 | 4,07E-11 |
| day5 h.aka sal15 temp16 - day2 h.rot sal30 temp16 | 64,88095 | 7,85863 | 213,4551 | 8,256013 | 4,07E-11 |
| day5 h.aka sal15 temp16 - day3 h.rot sal30 temp16 | 64,88095 | 7,85863 | 213,4551 | 8,256013 | 4,07E-11 |
| day5 h.aka sal15 temp16 - day4 h.rot sal30 temp16 | 64,88095 | 7,85863 | 213,4551 | 8,256013 | 4,07E-11 |
| day5 h.aka sal15 temp16 - day5 h.rot sal30 temp16 | 64,88095 | 7,85863 | 213,4551 | 8,256013 | 4,07E-11 |
| day5 h.aka sal15 temp16 - day0 both sal15 temp22 | 64,88095 | 7,85863 | 213,4551 | 8,256013 | 4,07E-11 |
| day5 h.aka sal15 temp16 - day1 both sal15 temp22 | 64,88095 | 7,85863 | 213,4551 | 8,256013 | 4,07E-11 |
| day5 h.aka sal15 temp16 - day2 both sal15 temp22 | 64,88095 | 7,85863 | 213,4551 | 8,256013 | 4,07E-11 |
| day5 h.aka sal15 temp16 - day3 both sal15 temp22 | 47,38095 | 7,85863 | 213,4551 | 6,029162 | 1,72E-05 |
| day5 h.aka sal15 temp16 - day4 both sal15 temp22 | 39,88095 | 7,85863 | 213,4551 | 5,074797 | 0,001749 |
| day5 h.aka sal15 temp16 - day5 both sal15 temp22 | 58,63095 | 7,85863 | 213,4551 | 7,460709 | 5,44E-09 |
| day5 h.aka sal15 temp16 - day0 h.aka sal15 temp22 | 64,88095 | 7,85863 | 213,4551 | 8,256013 | 4,07E-11 |
| day5 h.aka sal15 temp16 - day1 h.aka sal15 temp22 | 56,54762 | 7,85863 | 213,4551 | 7,195608 | 2,63E-08 |
| day5 h.aka sal15 temp16 - day4 h.aka sal15 temp22 | -35,119 | 7,85863 | 213,4551 | -4,46885 | 0,021053 |
| day5 h.aka sal15 temp16 - day0 h.rot sal15 temp22 | 64,88095 | 7,85863 | 213,4551 | 8,256013 | 4,07E-11 |
| day5 h.aka sal15 temp16 - day1 h.rot sal15 temp22 | 64,88095 | 7,85863 | 213,4551 | 8,256013 | 4,07E-11 |
| day5 h.aka sal15 temp16 - day2 h.rot sal15 temp22 | 58,18452 | 7,85863 | 213,4551 | 7,403902 | 7,64E-09 |
| day5 h.aka sal15 temp16 - day3 h.rot sal15 temp22 | 64,88095 | 7,85863 | 213,4551 | 8,256013 | 4,07E-11 |
| day5 h.aka sal15 temp16 - day4 h.rot sal15 temp22 | 64,88095 | 7,85863 | 213,4551 | 8,256013 | 4,07E-11 |
| day5 h.aka sal15 temp16 - day5 h.rot sal15 temp22 | 55,05952 | 7,85863 | 213,4551 | 7,00625 | 7,93E-08 |
| day5 h.aka sal15 temp16 - day0 both sal30 temp22 | 64,88095 | 7,85863 | 213,4551 | 8,256013 | 4,07E-11 |
| day5 h.aka sal15 temp16 - day1 both sal30 temp22 | 64,88095 | 7,85863 | 213,4551 | 8,256013 | 4,07E-11 |
| day5 h.aka sal15 temp16 - day2 both sal30 temp22 | 64,88095 | 7,85863 | 213,4551 | 8,256013 | 4,07E-11 |
| day5 h.aka sal15 temp16 - day3 both sal30 temp22 | 59,25595 | 7,85863 | 213,4551 | 7,54024 | 3,37E-09 |
| day5 h.aka sal15 temp16 - day4 both sal30 temp22 | 64,88095 | 7,85863 | 213,4551 | 8,256013 | 4,07E-11 |
| day5 h.aka sal15 temp16 - day5 both sal30 temp22 | 56,54762 | 7,85863 | 213,4551 | 7,195608 | 2,63E-08 |
| day5 h.aka sal15 temp16 - day0 h.aka sal30 temp22 | 64,88095 | 7,85863 | 213,4551 | 8,256013 | 4,07E-11 |
| day5 h.aka sal15 temp16 - day1 h.aka sal30 temp22 | 64,88095 | 7,85863 | 213,4551 | 8,256013 | 4,07E-11 |
| day5 h.aka sal15 temp16 - day2 h.aka sal30 temp22 | 43,63095 | 7,85863 | 213,4551 | 5,55198 | 0,00019 |
| day5 h.aka sal15 temp16 - day0 h.rot sal30 temp22 | 64,88095 | 7,85863 | 213,4551 | 8,256013 | 4,07E-11 |
| day5 h.aka sal15 temp16 - day1 h.rot sal30 temp22 | 64,88095 | 7,85863 | 213,4551 | 8,256013 | 4,07E-11 |
| day5 h.aka sal15 temp16 - day2 h.rot sal30 temp22 | 60,71429 | 7,85863 | 213,4551 | 7,725811 | 1,09E-09 |
| day5 h.aka sal15 temp16 - day3 h.rot sal30 temp22 | 56,13095 | 7,85863 | 213,4551 | 7,142588 | 3,59E-08 |
| day5 h.aka sal15 temp16 - day4 h.rot sal30 temp22 | 64,88095 | 7,85863 | 213,4551 | 8,256013 | 4,07E-11 |
| day5 h.aka sal15 temp16 - day5 h.rot sal30 temp22 | 55,71429 | 7,85863 | 213,4551 | 7,089568 | 4,89E-08 |
| day0 h.rot sal15 temp16 - day5 h.aka sal30 temp16 | -41,6667 | 7,85863 | 213,4551 | -5,30203 | 0,000624 |
| day0 h.rot sal15 temp16 - day2 h.aka sal15 temp22 | -54,5833 | 7,85863 | 213,4551 | -6,94566 | 1,13E-07 |
| day0 h.rot sal15 temp16 - day3 h.aka sal15 temp22 | -90 | 7,85863 | 213,4551 | -11,4524 | 2,87E-13 |
| day0 h.rot sal15 temp16 - day4 h.aka sal15 temp22 | -100 | 7,85863 | 213,4551 | -12,7249 | 9,56E-14 |
| day0 h.rot sal15 temp16 - day5 h.aka sal15 temp22 | -93,75 | 7,85863 | 213,4551 | -11,9296 | 1,43E-13 |

| | | | | | |
|---|---|---|---|---|---|
| day0 h.rot sal15 temp16 - day3 h.aka sal30 temp22 | -68,75 | 7,85863 | 213,4551 | -8,74834 | 2,44E-12 |
| day0 h.rot sal15 temp16 - day4 h.aka sal30 temp22 | -66,6667 | 7,85863 | 213,4551 | -8,48324 | 1,01E-11 |
| day0 h.rot sal15 temp16 - day5 h.aka sal30 temp22 | -91,6667 | 7,85863 | 213,4551 | -11,6645 | 2,1E-13 |
| day1 h.rot sal15 temp16 - day5 h.aka sal30 temp16 | -33,3333 | 7,85863 | 213,4551 | -4,24162 | 0,047643 |
| day1 h.rot sal15 temp16 - day2 h.aka sal15 temp22 | -46,25 | 7,85863 | 213,4551 | -5,88525 | 3,62E-05 |
| day1 h.rot sal15 temp16 - day3 h.aka sal15 temp22 | -81,6667 | 7,85863 | 213,4551 | -10,392 | 7,11E-13 |
| day1 h.rot sal15 temp16 - day4 h.aka sal15 temp22 | -91,6667 | 7,85863 | 213,4551 | -11,6645 | 2,1E-13 |
| day1 h.rot sal15 temp16 - day5 h.aka sal15 temp22 | -85,4167 | 7,85863 | 213,4551 | -10,8692 | 5,63E-13 |
| day1 h.rot sal15 temp16 - day3 h.aka sal30 temp22 | -60,4167 | 7,85863 | 213,4551 | -7,68794 | 1,37E-09 |
| day1 h.rot sal15 temp16 - day4 h.aka sal30 temp22 | -58,3333 | 7,85863 | 213,4551 | -7,42284 | 6,82E-09 |
| day1 h.rot sal15 temp16 - day5 h.aka sal30 temp22 | -83,3333 | 7,85863 | 213,4551 | -10,6041 | 6,96E-13 |
| day2 h.rot sal15 temp16 - day5 h.aka sal30 temp16 | -41,6667 | 7,85863 | 213,4551 | -5,30203 | 0,000624 |
| day2 h.rot sal15 temp16 - day2 h.aka sal15 temp22 | -54,5833 | 7,85863 | 213,4551 | -6,94566 | 1,13E-07 |
| day2 h.rot sal15 temp16 - day3 h.aka sal15 temp22 | -90 | 7,85863 | 213,4551 | -11,4524 | 2,87E-13 |
| day2 h.rot sal15 temp16 - day4 h.aka sal15 temp22 | -100 | 7,85863 | 213,4551 | -12,7249 | 9,56E-14 |
| day2 h.rot sal15 temp16 - day5 h.aka sal15 temp22 | -93,75 | 7,85863 | 213,4551 | -11,9296 | 1,43E-13 |
| day2 h.rot sal15 temp16 - day3 h.aka sal30 temp22 | -68,75 | 7,85863 | 213,4551 | -8,74834 | 2,44E-12 |
| day2 h.rot sal15 temp16 - day4 h.aka sal30 temp22 | -66,6667 | 7,85863 | 213,4551 | -8,48324 | 1,01E-11 |
| day2 h.rot sal15 temp16 - day5 h.aka sal30 temp22 | -91,6667 | 7,85863 | 213,4551 | -11,6645 | 2,1E-13 |
| day3 h.rot sal15 temp16 - day5 h.aka sal30 temp16 | -41,6667 | 7,85863 | 213,4551 | -5,30203 | 0,000624 |
| day3 h.rot sal15 temp16 - day2 h.aka sal15 temp22 | -54,5833 | 7,85863 | 213,4551 | -6,94566 | 1,13E-07 |
| day3 h.rot sal15 temp16 - day3 h.aka sal15 temp22 | -90 | 7,85863 | 213,4551 | -11,4524 | 2,87E-13 |
| day3 h.rot sal15 temp16 - day4 h.aka sal15 temp22 | -100 | 7,85863 | 213,4551 | -12,7249 | 9,56E-14 |
| day3 h.rot sal15 temp16 - day5 h.aka sal15 temp22 | -93,75 | 7,85863 | 213,4551 | -11,9296 | 1,43E-13 |
| day3 h.rot sal15 temp16 - day3 h.aka sal30 temp22 | -68,75 | 7,85863 | 213,4551 | -8,74834 | 2,44E-12 |
| day3 h.rot sal15 temp16 - day4 h.aka sal30 temp22 | -66,6667 | 7,85863 | 213,4551 | -8,48324 | 1,01E-11 |
| day3 h.rot sal15 temp16 - day5 h.aka sal30 temp22 | -91,6667 | 7,85863 | 213,4551 | -11,6645 | 2,1E-13 |
| day4 h.rot sal15 temp16 - day5 h.aka sal30 temp16 | -33,3333 | 7,85863 | 213,4551 | -4,24162 | 0,047643 |
| day4 h.rot sal15 temp16 - day2 h.aka sal15 temp22 | -46,25 | 7,85863 | 213,4551 | -5,88525 | 3,62E-05 |
| day4 h.rot sal15 temp16 - day3 h.aka sal15 temp22 | -81,6667 | 7,85863 | 213,4551 | -10,392 | 7,11E-13 |
| day4 h.rot sal15 temp16 - day4 h.aka sal15 temp22 | -91,6667 | 7,85863 | 213,4551 | -11,6645 | 2,1E-13 |
| day4 h.rot sal15 temp16 - day5 h.aka sal15 temp22 | -85,4167 | 7,85863 | 213,4551 | -10,8692 | 5,63E-13 |
| day4 h.rot sal15 temp16 - day3 h.aka sal30 temp22 | -60,4167 | 7,85863 | 213,4551 | -7,68794 | 1,37E-09 |
| day4 h.rot sal15 temp16 - day4 h.aka sal30 temp22 | -58,3333 | 7,85863 | 213,4551 | -7,42284 | 6,82E-09 |
| day4 h.rot sal15 temp16 - day5 h.aka sal30 temp22 | -83,3333 | 7,85863 | 213,4551 | -10,6041 | 6,96E-13 |
| day5 h.rot sal15 temp16 - day2 h.aka sal15 temp22 | -45,4167 | 7,85863 | 213,4551 | -5,77921 | 6,2E-05 |
| day5 h.rot sal15 temp16 - day3 h.aka sal15 temp22 | -80,8333 | 7,85863 | 213,4551 | -10,2859 | 7,39E-13 |
| day5 h.rot sal15 temp16 - day4 h.aka sal15 temp22 | -90,8333 | 7,85863 | 213,4551 | -11,5584 | 2,44E-13 |
| day5 h.rot sal15 temp16 - day5 h.aka sal15 temp22 | -84,5833 | 7,85863 | 213,4551 | -10,7631 | 6E-13 |
| day5 h.rot sal15 temp16 - day3 h.aka sal30 temp22 | -59,5833 | 7,85863 | 213,4551 | -7,5819 | 2,62E-09 |
| day5 h.rot sal15 temp16 - day4 h.aka sal30 temp22 | -57,5 | 7,85863 | 213,4551 | -7,3168 | 1,28E-08 |
| day5 h.rot sal15 temp16 - day5 h.aka sal30 temp22 | -82,5 | 7,85863 | 213,4551 | -10,498 | 7,08E-13 |
| day0 both sal30 temp16 - day5 h.aka sal30 temp16 | -41,6667 | 7,85863 | 213,4551 | -5,30203 | 0,000624 |
| day0 both sal30 temp16 - day2 h.aka sal15 temp22 | -54,5833 | 7,85863 | 213,4551 | -6,94566 | 1,13E-07 |
| day0 both sal30 temp16 - day3 h.aka sal15 temp22 | -90 | 7,85863 | 213,4551 | -11,4524 | 2,87E-13 |
| day0 both sal30 temp16 - day4 h.aka sal15 temp22 | -100 | 7,85863 | 213,4551 | -12,7249 | 9,56E-14 |
| day0 both sal30 temp16 - day5 h.aka sal15 temp22 | -93,75 | 7,85863 | 213,4551 | -11,9296 | 1,43E-13 |
| day0 both sal30 temp16 - day3 h.aka sal30 temp22 | -68,75 | 7,85863 | 213,4551 | -8,74834 | 2,44E-12 |
| day0 both sal30 temp16 - day4 h.aka sal30 temp22 | -66,6667 | 7,85863 | 213,4551 | -8,48324 | 1,01E-11 |
| day0 both sal30 temp16 - day5 h.aka sal30 temp22 | -91,6667 | 7,85863 | 213,4551 | -11,6645 | 2,1E-13 |
| day1 both sal30 temp16 - day5 h.aka sal30 temp16 | -41,6667 | 7,85863 | 213,4551 | -5,30203 | 0,000624 |
| day1 both sal30 temp16 - day2 h.aka sal15 temp22 | -54,5833 | 7,85863 | 213,4551 | -6,94566 | 1,13E-07 |
| day1 both sal30 temp16 - day3 h.aka sal15 temp22 | -90 | 7,85863 | 213,4551 | -11,4524 | 2,87E-13 |
| day1 both sal30 temp16 - day4 h.aka sal15 temp22 | -100 | 7,85863 | 213,4551 | -12,7249 | 9,56E-14 |
| day1 both sal30 temp16 - day5 h.aka sal15 temp22 | -93,75 | 7,85863 | 213,4551 | -11,9296 | 1,43E-13 |
| day1 both sal30 temp16 - day3 h.aka sal30 temp22 | -68,75 | 7,85863 | 213,4551 | -8,74834 | 2,44E-12 |
| day1 both sal30 temp16 - day4 h.aka sal30 temp22 | -66,6667 | 7,85863 | 213,4551 | -8,48324 | 1,01E-11 |
| day1 both sal30 temp16 - day5 h.aka sal30 temp22 | -91,6667 | 7,85863 | 213,4551 | -11,6645 | 2,1E-13 |
| day2 both sal30 temp16 - day5 h.aka sal30 temp16 | -41,6667 | 7,85863 | 213,4551 | -5,30203 | 0,000624 |
| day2 both sal30 temp16 - day2 h.aka sal15 temp22 | -54,5833 | 7,85863 | 213,4551 | -6,94566 | 1,13E-07 |
| day2 both sal30 temp16 - day3 h.aka sal15 temp22 | -90 | 7,85863 | 213,4551 | -11,4524 | 2,87E-13 |
| day2 both sal30 temp16 - day4 h.aka sal15 temp22 | -100 | 7,85863 | 213,4551 | -12,7249 | 9,56E-14 |
| day2 both sal30 temp16 - day5 h.aka sal15 temp22 | -93,75 | 7,85863 | 213,4551 | -11,9296 | 1,43E-13 |
| day2 both sal30 temp16 - day3 h.aka sal30 temp22 | -68,75 | 7,85863 | 213,4551 | -8,74834 | 2,44E-12 |
| day2 both sal30 temp16 - day4 h.aka sal30 temp22 | -66,6667 | 7,85863 | 213,4551 | -8,48324 | 1,01E-11 |
| day2 both sal30 temp16 - day5 h.aka sal30 temp22 | -91,6667 | 7,85863 | 213,4551 | -11,6645 | 2,1E-13 |
| day3 both sal30 temp16 - day5 h.aka sal30 temp16 | -41,6667 | 7,85863 | 213,4551 | -5,30203 | 0,000624 |
| day3 both sal30 temp16 - day2 h.aka sal15 temp22 | -54,5833 | 7,85863 | 213,4551 | -6,94566 | 1,13E-07 |
| day3 both sal30 temp16 - day3 h.aka sal15 temp22 | -90 | 7,85863 | 213,4551 | -11,4524 | 2,87E-13 |
| day3 both sal30 temp16 - day4 h.aka sal15 temp22 | -100 | 7,85863 | 213,4551 | -12,7249 | 9,56E-14 |
| day3 both sal30 temp16 - day5 h.aka sal15 temp22 | -93,75 | 7,85863 | 213,4551 | -11,9296 | 1,43E-13 |
| day3 both sal30 temp16 - day3 h.aka sal30 temp22 | -68,75 | 7,85863 | 213,4551 | -8,74834 | 2,44E-12 |
| day3 both sal30 temp16 - day4 h.aka sal30 temp22 | -66,6667 | 7,85863 | 213,4551 | -8,48324 | 1,01E-11 |
| day3 both sal30 temp16 - day5 h.aka sal30 temp22 | -91,6667 | 7,85863 | 213,4551 | -11,6645 | 2,1E-13 |
| day4 both sal30 temp16 - day5 h.aka sal30 temp16 | -33,3333 | 7,85863 | 213,4551 | -4,24162 | 0,047643 |
| day4 both sal30 temp16 - day2 h.aka sal15 temp22 | -46,25 | 7,85863 | 213,4551 | -5,88525 | 3,62E-05 |
| day4 both sal30 temp16 - day3 h.aka sal15 temp22 | -81,6667 | 7,85863 | 213,4551 | -10,392 | 7,11E-13 |
| day4 both sal30 temp16 - day4 h.aka sal15 temp22 | -91,6667 | 7,85863 | 213,4551 | -11,6645 | 2,1E-13 |
| day4 both sal30 temp16 - day5 h.aka sal15 temp22 | -85,4167 | 7,85863 | 213,4551 | -10,8692 | 5,63E-13 |
| day4 both sal30 temp16 - day3 h.aka sal30 temp22 | -60,4167 | 7,85863 | 213,4551 | -7,68794 | 1,37E-09 |
| day4 both sal30 temp16 - day4 h.aka sal30 temp22 | -58,3333 | 7,85863 | 213,4551 | -7,42284 | 6,82E-09 |
| day4 both sal30 temp16 - day5 h.aka sal30 temp22 | -83,3333 | 7,85863 | 213,4551 | -10,6041 | 6,96E-13 |
| day5 both sal30 temp16 - day5 h.aka sal30 temp16 | -35,4167 | 7,85863 | 213,4551 | -4,50672 | 0,018249 |
| day5 both sal30 temp16 - day2 h.aka sal15 temp22 | -48,3333 | 7,85863 | 213,4551 | -6,15035 | 9,13E-06 |

| | | | | | |
|---|---|---|---|---|---|
| day5 both sal30 temp16 - day3 h.aka sal15 temp22 | -83,75 | 7,85863 | 213,4551 | -10,6571 | 6,43E-13 |
| day5 both sal30 temp16 - day4 h.aka sal15 temp22 | -93,75 | 7,85863 | 213,4551 | -11,9296 | 1,43E-13 |
| day5 both sal30 temp16 - day5 h.aka sal15 temp22 | -87,5 | 7,85863 | 213,4551 | -11,1343 | 4,35E-13 |
| day5 both sal30 temp16 - day3 h.aka sal30 temp22 | -62,5 | 7,85863 | 213,4551 | -7,95304 | 2,68E-10 |
| day5 both sal30 temp16 - day4 h.aka sal30 temp22 | -60,4167 | 7,85863 | 213,4551 | -7,68794 | 1,37E-09 |
| day5 both sal30 temp16 - day5 h.aka sal30 temp22 | -85,4167 | 7,85863 | 213,4551 | -10,8692 | 5,63E-13 |
| day0 h.aka sal30 temp16 - day5 h.aka sal30 temp16 | -41,6667 | 7,664356 | 180 | -5,43642 | 0,000387 |
| day0 h.aka sal30 temp16 - day2 h.aka sal15 temp22 | -54,5833 | 7,85863 | 213,4551 | -6,94566 | 1,13E-07 |
| day0 h.aka sal30 temp16 - day3 h.aka sal15 temp22 | -90 | 7,85863 | 213,4551 | -11,4524 | 2,87E-13 |
| day0 h.aka sal30 temp16 - day4 h.aka sal15 temp22 | -100 | 7,85863 | 213,4551 | -12,7249 | 9,56E-14 |
| day0 h.aka sal30 temp16 - day5 h.aka sal15 temp22 | -93,75 | 7,85863 | 213,4551 | -11,9296 | 1,43E-13 |
| day0 h.aka sal30 temp16 - day3 h.aka sal30 temp22 | -68,75 | 7,85863 | 213,4551 | -8,74834 | 2,44E-12 |
| day0 h.aka sal30 temp16 - day4 h.aka sal30 temp22 | -66,6667 | 7,85863 | 213,4551 | -8,48324 | 1,01E-11 |
| day0 h.aka sal30 temp16 - day5 h.aka sal30 temp22 | -91,6667 | 7,85863 | 213,4551 | -11,6645 | 2,1E-13 |
| day1 h.aka sal30 temp16 - day5 h.aka sal30 temp16 | -41,6667 | 7,664356 | 180 | -5,43642 | 0,000387 |
| day1 h.aka sal30 temp16 - day2 h.aka sal15 temp22 | -54,5833 | 7,85863 | 213,4551 | -6,94566 | 1,13E-07 |
| day1 h.aka sal30 temp16 - day3 h.aka sal15 temp22 | -90 | 7,85863 | 213,4551 | -11,4524 | 2,87E-13 |
| day1 h.aka sal30 temp16 - day4 h.aka sal15 temp22 | -100 | 7,85863 | 213,4551 | -12,7249 | 9,56E-14 |
| day1 h.aka sal30 temp16 - day5 h.aka sal15 temp22 | -93,75 | 7,85863 | 213,4551 | -11,9296 | 1,43E-13 |
| day1 h.aka sal30 temp16 - day3 h.aka sal30 temp22 | -68,75 | 7,85863 | 213,4551 | -8,74834 | 2,44E-12 |
| day1 h.aka sal30 temp16 - day4 h.aka sal30 temp22 | -66,6667 | 7,85863 | 213,4551 | -8,48324 | 1,01E-11 |
| day1 h.aka sal30 temp16 - day5 h.aka sal30 temp22 | -91,6667 | 7,85863 | 213,4551 | -11,6645 | 2,1E-13 |
| day2 h.aka sal30 temp16 - day5 h.aka sal30 temp16 | -41,6667 | 7,664356 | 180 | -5,43642 | 0,000387 |
| day2 h.aka sal30 temp16 - day2 h.aka sal15 temp22 | -54,5833 | 7,85863 | 213,4551 | -6,94566 | 1,13E-07 |
| day2 h.aka sal30 temp16 - day3 h.aka sal15 temp22 | -90 | 7,85863 | 213,4551 | -11,4524 | 2,87E-13 |
| day2 h.aka sal30 temp16 - day4 h.aka sal15 temp22 | -100 | 7,85863 | 213,4551 | -12,7249 | 9,56E-14 |
| day2 h.aka sal30 temp16 - day5 h.aka sal15 temp22 | -93,75 | 7,85863 | 213,4551 | -11,9296 | 1,43E-13 |
| day2 h.aka sal30 temp16 - day3 h.aka sal30 temp22 | -68,75 | 7,85863 | 213,4551 | -8,74834 | 2,44E-12 |
| day2 h.aka sal30 temp16 - day4 h.aka sal30 temp22 | -66,6667 | 7,85863 | 213,4551 | -8,48324 | 1,01E-11 |
| day2 h.aka sal30 temp16 - day5 h.aka sal30 temp22 | -91,6667 | 7,85863 | 213,4551 | -11,6645 | 2,1E-13 |
| day3 h.aka sal30 temp16 - day2 h.aka sal15 temp22 | -43,125 | 7,85863 | 213,4551 | -5,4876 | 0,00026 |
| day3 h.aka sal30 temp16 - day3 h.aka sal15 temp22 | -78,5417 | 7,85863 | 213,4551 | -9,99432 | 8,03E-13 |
| day3 h.aka sal30 temp16 - day4 h.aka sal15 temp22 | -88,5417 | 7,85863 | 213,4551 | -11,2668 | 3,61E-13 |
| day3 h.aka sal30 temp16 - day5 h.aka sal15 temp22 | -82,2917 | 7,85863 | 213,4551 | -10,4715 | 7,09E-13 |
| day3 h.aka sal30 temp16 - day3 h.aka sal30 temp22 | -57,2917 | 7,85863 | 213,4551 | -7,29029 | 1,5E-08 |
| day3 h.aka sal30 temp16 - day4 h.aka sal30 temp22 | -55,2083 | 7,85863 | 213,4551 | -7,02519 | 7,11E-08 |
| day3 h.aka sal30 temp16 - day5 h.aka sal30 temp22 | -80,2083 | 7,85863 | 213,4551 | -10,2064 | 7,53E-13 |
| day4 h.aka sal30 temp16 - day5 h.aka sal30 temp16 | -38,0952 | 7,664356 | 180 | -4,97044 | 0,003064 |
| day4 h.aka sal30 temp16 - day2 h.aka sal15 temp22 | -51,0119 | 7,85863 | 213,4551 | -6,4912 | 1,45E-06 |
| day4 h.aka sal30 temp16 - day3 h.aka sal15 temp22 | -86,4286 | 7,85863 | 213,4551 | -10,9979 | 4,8E-13 |
| day4 h.aka sal30 temp16 - day4 h.aka sal15 temp22 | -96,4286 | 7,85863 | 213,4551 | -12,2704 | 1,09E-13 |
| day4 h.aka sal30 temp16 - day5 h.aka sal15 temp22 | -90,1786 | 7,85863 | 213,4551 | -11,4751 | 2,86E-13 |
| day4 h.aka sal30 temp16 - day3 h.aka sal30 temp22 | -65,1786 | 7,85863 | 213,4551 | -8,29389 | 3,21E-11 |
| day4 h.aka sal30 temp16 - day4 h.aka sal30 temp22 | -63,0952 | 7,85863 | 213,4551 | -8,02878 | 1,67E-10 |
| day4 h.aka sal30 temp16 - day5 h.aka sal30 temp22 | -88,0952 | 7,85863 | 213,4551 | -11,21 | 3,96E-13 |
| day5 h.aka sal30 temp16 - day0 h.rot sal30 temp16 | 41,66667 | 7,85863 | 213,4551 | 5,302027 | 0,000624 |
| day5 h.aka sal30 temp16 - day1 h.rot sal30 temp16 | 41,66667 | 7,85863 | 213,4551 | 5,302027 | 0,000624 |
| day5 h.aka sal30 temp16 - day2 h.rot sal30 temp16 | 41,66667 | 7,85863 | 213,4551 | 5,302027 | 0,000624 |
| day5 h.aka sal30 temp16 - day3 h.rot sal30 temp16 | 41,66667 | 7,85863 | 213,4551 | 5,302027 | 0,000624 |
| day5 h.aka sal30 temp16 - day4 h.rot sal30 temp16 | 41,66667 | 7,85863 | 213,4551 | 5,302027 | 0,000624 |
| day5 h.aka sal30 temp16 - day5 h.rot sal30 temp16 | 41,66667 | 7,85863 | 213,4551 | 5,302027 | 0,000624 |
| day5 h.aka sal30 temp16 - day0 both sal15 temp22 | 41,66667 | 7,85863 | 213,4551 | 5,302027 | 0,000624 |
| day5 h.aka sal30 temp16 - day1 both sal15 temp22 | 41,66667 | 7,85863 | 213,4551 | 5,302027 | 0,000624 |
| day5 h.aka sal30 temp16 - day2 both sal15 temp22 | 41,66667 | 7,85863 | 213,4551 | 5,302027 | 0,000624 |
| day5 h.aka sal30 temp16 - day5 both sal15 temp22 | 35,41667 | 7,85863 | 213,4551 | 4,506723 | 0,018249 |
| day5 h.aka sal30 temp16 - day0 h.aka sal15 temp22 | 41,66667 | 7,85863 | 213,4551 | 5,302027 | 0,000624 |
| day5 h.aka sal30 temp16 - day1 h.aka sal15 temp22 | 33,33333 | 7,85863 | 213,4551 | 4,241622 | 0,047643 |
| day5 h.aka sal30 temp16 - day3 h.aka sal15 temp22 | -48,3333 | 7,85863 | 213,4551 | -6,15035 | 9,13E-06 |
| day5 h.aka sal30 temp16 - day4 h.aka sal15 temp22 | -58,3333 | 7,85863 | 213,4551 | -7,42284 | 6,82E-09 |
| day5 h.aka sal30 temp16 - day5 h.aka sal15 temp22 | -52,0833 | 7,85863 | 213,4551 | -6,62753 | 6,83E-07 |
| day5 h.aka sal30 temp16 - day0 h.rot sal15 temp22 | 41,66667 | 7,85863 | 213,4551 | 5,302027 | 0,000624 |
| day5 h.aka sal30 temp16 - day1 h.rot sal15 temp22 | 41,66667 | 7,85863 | 213,4551 | 5,302027 | 0,000624 |
| day5 h.aka sal30 temp16 - day2 h.rot sal15 temp22 | 34,97024 | 7,85863 | 213,4551 | 4,449916 | 0,022597 |
| day5 h.aka sal30 temp16 - day3 h.rot sal15 temp22 | 41,66667 | 7,85863 | 213,4551 | 5,302027 | 0,000624 |
| day5 h.aka sal30 temp16 - day4 h.rot sal15 temp22 | 41,66667 | 7,85863 | 213,4551 | 5,302027 | 0,000624 |
| day5 h.aka sal30 temp16 - day0 both sal30 temp22 | 41,66667 | 7,85863 | 213,4551 | 5,302027 | 0,000624 |
| day5 h.aka sal30 temp16 - day1 both sal30 temp22 | 41,66667 | 7,85863 | 213,4551 | 5,302027 | 0,000624 |
| day5 h.aka sal30 temp16 - day2 both sal30 temp22 | 41,66667 | 7,85863 | 213,4551 | 5,302027 | 0,000624 |
| day5 h.aka sal30 temp16 - day3 both sal30 temp22 | 36,04167 | 7,85863 | 213,4551 | 4,586253 | 0,013438 |
| day5 h.aka sal30 temp16 - day4 both sal30 temp22 | 41,66667 | 7,85863 | 213,4551 | 5,302027 | 0,000624 |
| day5 h.aka sal30 temp16 - day5 both sal30 temp22 | 33,33333 | 7,85863 | 213,4551 | 4,241622 | 0,047643 |
| day5 h.aka sal30 temp16 - day0 h.aka sal30 temp22 | 41,66667 | 7,85863 | 213,4551 | 5,302027 | 0,000624 |
| day5 h.aka sal30 temp16 - day1 h.aka sal30 temp22 | 41,66667 | 7,85863 | 213,4551 | 5,302027 | 0,000624 |
| day5 h.aka sal30 temp16 - day5 h.aka sal30 temp22 | -50 | 7,85863 | 213,4551 | -6,36243 | 2,93E-06 |
| day5 h.aka sal30 temp16 - day0 h.rot sal30 temp22 | 41,66667 | 7,85863 | 213,4551 | 5,302027 | 0,000624 |
| day5 h.aka sal30 temp16 - day1 h.rot sal30 temp22 | 41,66667 | 7,85863 | 213,4551 | 5,302027 | 0,000624 |
| day5 h.aka sal30 temp16 - day2 h.rot sal30 temp22 | 37,5 | 7,85863 | 213,4551 | 4,771824 | 0,006388 |
| day5 h.aka sal30 temp16 - day4 h.rot sal30 temp22 | 41,66667 | 7,85863 | 213,4551 | 5,302027 | 0,000624 |
| day0 h.rot sal30 temp16 - day2 h.aka sal15 temp22 | -54,5833 | 7,85863 | 213,4551 | -6,94566 | 1,13E-07 |
| day0 h.rot sal30 temp16 - day3 h.aka sal15 temp22 | -90 | 7,85863 | 213,4551 | -11,4524 | 2,87E-13 |
| day0 h.rot sal30 temp16 - day4 h.aka sal15 temp22 | -100 | 7,85863 | 213,4551 | -12,7249 | 9,56E-14 |
| day0 h.rot sal30 temp16 - day5 h.aka sal15 temp22 | -93,75 | 7,85863 | 213,4551 | -11,9296 | 1,43E-13 |
| day0 h.rot sal30 temp16 - day3 h.aka sal30 temp22 | -68,75 | 7,85863 | 213,4551 | -8,74834 | 2,44E-12 |
| day0 h.rot sal30 temp16 - day4 h.aka sal30 temp22 | -66,6667 | 7,85863 | 213,4551 | -8,48324 | 1,01E-11 |

| | | | | | |
|---|---|---|---|---|---|
| day0 h.rot sal30 temp16 - day5 h.aka sal30 temp22 | -91,6667 | 7,85863 | 213,4551 | -11,6645 | 2,1E-13 |
| day1 h.rot sal30 temp16 - day2 h.aka sal15 temp22 | -54,5833 | 7,85863 | 213,4551 | -6,94566 | 1,13E-07 |
| day1 h.rot sal30 temp16 - day3 h.aka sal15 temp22 | -90 | 7,85863 | 213,4551 | -11,4524 | 2,87E-13 |
| day1 h.rot sal30 temp16 - day4 h.aka sal15 temp22 | -100 | 7,85863 | 213,4551 | -12,7249 | 9,56E-14 |
| day1 h.rot sal30 temp16 - day5 h.aka sal15 temp22 | -93,75 | 7,85863 | 213,4551 | -11,9296 | 1,43E-13 |
| day1 h.rot sal30 temp16 - day3 h.aka sal30 temp22 | -68,75 | 7,85863 | 213,4551 | -8,74834 | 2,44E-12 |
| day1 h.rot sal30 temp16 - day4 h.aka sal30 temp22 | -66,6667 | 7,85863 | 213,4551 | -8,48324 | 1,01E-11 |
| day1 h.rot sal30 temp16 - day5 h.aka sal30 temp22 | -91,6667 | 7,85863 | 213,4551 | -11,6645 | 2,1E-13 |
| day2 h.rot sal30 temp16 - day2 h.aka sal15 temp22 | -54,5833 | 7,85863 | 213,4551 | -6,94566 | 1,13E-07 |
| day2 h.rot sal30 temp16 - day3 h.aka sal15 temp22 | -90 | 7,85863 | 213,4551 | -11,4524 | 2,87E-13 |
| day2 h.rot sal30 temp16 - day4 h.aka sal15 temp22 | -100 | 7,85863 | 213,4551 | -12,7249 | 9,56E-14 |
| day2 h.rot sal30 temp16 - day5 h.aka sal15 temp22 | -93,75 | 7,85863 | 213,4551 | -11,9296 | 1,43E-13 |
| day2 h.rot sal30 temp16 - day3 h.aka sal30 temp22 | -68,75 | 7,85863 | 213,4551 | -8,74834 | 2,44E-12 |
| day2 h.rot sal30 temp16 - day4 h.aka sal30 temp22 | -66,6667 | 7,85863 | 213,4551 | -8,48324 | 1,01E-11 |
| day2 h.rot sal30 temp16 - day5 h.aka sal30 temp22 | -91,6667 | 7,85863 | 213,4551 | -11,6645 | 2,1E-13 |
| day3 h.rot sal30 temp16 - day2 h.aka sal15 temp22 | -54,5833 | 7,85863 | 213,4551 | -6,94566 | 1,13E-07 |
| day3 h.rot sal30 temp16 - day3 h.aka sal15 temp22 | -90 | 7,85863 | 213,4551 | -11,4524 | 2,87E-13 |
| day3 h.rot sal30 temp16 - day4 h.aka sal15 temp22 | -100 | 7,85863 | 213,4551 | -12,7249 | 9,56E-14 |
| day3 h.rot sal30 temp16 - day5 h.aka sal15 temp22 | -93,75 | 7,85863 | 213,4551 | -11,9296 | 1,43E-13 |
| day3 h.rot sal30 temp16 - day3 h.aka sal30 temp22 | -68,75 | 7,85863 | 213,4551 | -8,74834 | 2,44E-12 |
| day3 h.rot sal30 temp16 - day4 h.aka sal30 temp22 | -66,6667 | 7,85863 | 213,4551 | -8,48324 | 1,01E-11 |
| day3 h.rot sal30 temp16 - day5 h.aka sal30 temp22 | -91,6667 | 7,85863 | 213,4551 | -11,6645 | 2,1E-13 |
| day4 h.rot sal30 temp16 - day2 h.aka sal15 temp22 | -54,5833 | 7,85863 | 213,4551 | -6,94566 | 1,13E-07 |
| day4 h.rot sal30 temp16 - day3 h.aka sal15 temp22 | -90 | 7,85863 | 213,4551 | -11,4524 | 2,87E-13 |
| day4 h.rot sal30 temp16 - day4 h.aka sal15 temp22 | -100 | 7,85863 | 213,4551 | -12,7249 | 9,56E-14 |
| day4 h.rot sal30 temp16 - day5 h.aka sal15 temp22 | -93,75 | 7,85863 | 213,4551 | -11,9296 | 1,43E-13 |
| day4 h.rot sal30 temp16 - day3 h.aka sal30 temp22 | -68,75 | 7,85863 | 213,4551 | -8,74834 | 2,44E-12 |
| day4 h.rot sal30 temp16 - day4 h.aka sal30 temp22 | -66,6667 | 7,85863 | 213,4551 | -8,48324 | 1,01E-11 |
| day4 h.rot sal30 temp16 - day5 h.aka sal30 temp22 | -91,6667 | 7,85863 | 213,4551 | -11,6645 | 2,1E-13 |
| day5 h.rot sal30 temp16 - day2 h.aka sal15 temp22 | -54,5833 | 7,85863 | 213,4551 | -6,94566 | 1,13E-07 |
| day5 h.rot sal30 temp16 - day3 h.aka sal15 temp22 | -90 | 7,85863 | 213,4551 | -11,4524 | 2,87E-13 |
| day5 h.rot sal30 temp16 - day4 h.aka sal15 temp22 | -100 | 7,85863 | 213,4551 | -12,7249 | 9,56E-14 |
| day5 h.rot sal30 temp16 - day5 h.aka sal15 temp22 | -93,75 | 7,85863 | 213,4551 | -11,9296 | 1,43E-13 |
| day5 h.rot sal30 temp16 - day3 h.aka sal30 temp22 | -68,75 | 7,85863 | 213,4551 | -8,74834 | 2,44E-12 |
| day5 h.rot sal30 temp16 - day4 h.aka sal30 temp22 | -66,6667 | 7,85863 | 213,4551 | -8,48324 | 1,01E-11 |
| day5 h.rot sal30 temp16 - day5 h.aka sal30 temp22 | -91,6667 | 7,85863 | 213,4551 | -11,6645 | 2,1E-13 |
| day0 both sal15 temp22 - day2 h.aka sal15 temp22 | -54,5833 | 7,85863 | 213,4551 | -6,94566 | 1,13E-07 |
| day0 both sal15 temp22 - day3 h.aka sal15 temp22 | -90 | 7,85863 | 213,4551 | -11,4524 | 2,87E-13 |
| day0 both sal15 temp22 - day4 h.aka sal15 temp22 | -100 | 7,85863 | 213,4551 | -12,7249 | 9,56E-14 |
| day0 both sal15 temp22 - day5 h.aka sal15 temp22 | -93,75 | 7,85863 | 213,4551 | -11,9296 | 1,43E-13 |
| day0 both sal15 temp22 - day3 h.aka sal30 temp22 | -68,75 | 7,85863 | 213,4551 | -8,74834 | 2,44E-12 |
| day0 both sal15 temp22 - day4 h.aka sal30 temp22 | -66,6667 | 7,85863 | 213,4551 | -8,48324 | 1,01E-11 |
| day0 both sal15 temp22 - day5 h.aka sal30 temp22 | -91,6667 | 7,85863 | 213,4551 | -11,6645 | 2,1E-13 |
| day1 both sal15 temp22 - day2 h.aka sal15 temp22 | -54,5833 | 7,85863 | 213,4551 | -6,94566 | 1,13E-07 |
| day1 both sal15 temp22 - day3 h.aka sal15 temp22 | -90 | 7,85863 | 213,4551 | -11,4524 | 2,87E-13 |
| day1 both sal15 temp22 - day4 h.aka sal15 temp22 | -100 | 7,85863 | 213,4551 | -12,7249 | 9,56E-14 |
| day1 both sal15 temp22 - day5 h.aka sal15 temp22 | -93,75 | 7,85863 | 213,4551 | -11,9296 | 1,43E-13 |
| day1 both sal15 temp22 - day3 h.aka sal30 temp22 | -68,75 | 7,85863 | 213,4551 | -8,74834 | 2,44E-12 |
| day1 both sal15 temp22 - day4 h.aka sal30 temp22 | -66,6667 | 7,85863 | 213,4551 | -8,48324 | 1,01E-11 |
| day1 both sal15 temp22 - day5 h.aka sal30 temp22 | -91,6667 | 7,85863 | 213,4551 | -11,6645 | 2,1E-13 |
| day2 both sal15 temp22 - day2 h.aka sal15 temp22 | -54,5833 | 7,85863 | 213,4551 | -6,94566 | 1,13E-07 |
| day2 both sal15 temp22 - day3 h.aka sal15 temp22 | -90 | 7,85863 | 213,4551 | -11,4524 | 2,87E-13 |
| day2 both sal15 temp22 - day4 h.aka sal15 temp22 | -100 | 7,85863 | 213,4551 | -12,7249 | 9,56E-14 |
| day2 both sal15 temp22 - day5 h.aka sal15 temp22 | -93,75 | 7,85863 | 213,4551 | -11,9296 | 1,43E-13 |
| day2 both sal15 temp22 - day3 h.aka sal30 temp22 | -68,75 | 7,85863 | 213,4551 | -8,74834 | 2,44E-12 |
| day2 both sal15 temp22 - day4 h.aka sal30 temp22 | -66,6667 | 7,85863 | 213,4551 | -8,48324 | 1,01E-11 |
| day2 both sal15 temp22 - day5 h.aka sal30 temp22 | -91,6667 | 7,85863 | 213,4551 | -11,6645 | 2,1E-13 |
| day3 both sal15 temp22 - day2 h.aka sal15 temp22 | -37,0833 | 7,85863 | 213,4551 | -4,7188 | 0,007932 |
| day3 both sal15 temp22 - day3 h.aka sal15 temp22 | -72,5 | 7,85863 | 213,4551 | -9,22553 | 8,38E-13 |
| day3 both sal15 temp22 - day4 h.aka sal15 temp22 | -82,5 | 7,85863 | 213,4551 | -10,498 | 7,08E-13 |
| day3 both sal15 temp22 - day5 h.aka sal15 temp22 | -76,25 | 7,85863 | 213,4551 | -9,70271 | 7,56E-13 |
| day3 both sal15 temp22 - day3 h.aka sal30 temp22 | -51,25 | 7,85863 | 213,4551 | -6,52149 | 1,23E-06 |
| day3 both sal15 temp22 - day4 h.aka sal30 temp22 | -49,1667 | 7,85863 | 213,4551 | -6,25639 | 5,2E-06 |
| day3 both sal15 temp22 - day5 h.aka sal30 temp22 | -74,1667 | 7,85863 | 213,4551 | -9,43761 | 7,96E-13 |
| day4 both sal15 temp22 - day3 h.aka sal15 temp22 | -65 | 7,85863 | 213,4551 | -8,27116 | 3,7E-11 |
| day4 both sal15 temp22 - day4 h.aka sal15 temp22 | -75 | 7,85863 | 213,4551 | -9,54365 | 7,96E-13 |
| day4 both sal15 temp22 - day5 h.aka sal15 temp22 | -68,75 | 7,85863 | 213,4551 | -8,74834 | 2,44E-12 |
| day4 both sal15 temp22 - day3 h.aka sal30 temp22 | -43,75 | 7,85863 | 213,4551 | -5,56713 | 0,000177 |
| day4 both sal15 temp22 - day4 h.aka sal30 temp22 | -41,6667 | 7,85863 | 213,4551 | -5,30203 | 0,000624 |
| day4 both sal15 temp22 - day5 h.aka sal30 temp22 | -66,6667 | 7,85863 | 213,4551 | -8,48324 | 1,01E-11 |
| day5 both sal15 temp22 - day2 h.aka sal15 temp22 | -48,3333 | 7,85863 | 213,4551 | -6,15035 | 9,13E-06 |
| day5 both sal15 temp22 - day3 h.aka sal15 temp22 | -83,75 | 7,85863 | 213,4551 | -10,6571 | 6,43E-13 |
| day5 both sal15 temp22 - day4 h.aka sal15 temp22 | -93,75 | 7,85863 | 213,4551 | -11,9296 | 1,43E-13 |
| day5 both sal15 temp22 - day5 h.aka sal15 temp22 | -87,5 | 7,85863 | 213,4551 | -11,1343 | 4,35E-13 |
| day5 both sal15 temp22 - day3 h.aka sal30 temp22 | -62,5 | 7,85863 | 213,4551 | -7,95304 | 2,68E-10 |
| day5 both sal15 temp22 - day4 h.aka sal30 temp22 | -60,4167 | 7,85863 | 213,4551 | -7,68794 | 1,37E-09 |
| day5 both sal15 temp22 - day5 h.aka sal30 temp22 | -85,4167 | 7,85863 | 213,4551 | -10,8692 | 5,63E-13 |
| day0 h.aka sal15 temp22 - day2 h.aka sal15 temp22 | -54,5833 | 7,664356 | 180 | -7,12171 | 6,19E-08 |
| day0 h.aka sal15 temp22 - day3 h.aka sal15 temp22 | -90 | 7,664356 | 180 | -11,7427 | 2,34E-14 |
| day0 h.aka sal15 temp22 - day4 h.aka sal15 temp22 | -100 | 7,664356 | 180 | -13,0474 | 0 |
| day0 h.aka sal15 temp22 - day5 h.aka sal15 temp22 | -93,75 | 7,664356 | 180 | -12,2319 | 0 |
| day0 h.aka sal15 temp22 - day3 h.aka sal30 temp22 | -68,75 | 7,85863 | 213,4551 | -8,74834 | 2,44E-12 |
| day0 h.aka sal15 temp22 - day4 h.aka sal30 temp22 | -66,6667 | 7,85863 | 213,4551 | -8,48324 | 1,01E-11 |
| day0 h.aka sal15 temp22 - day5 h.aka sal30 temp22 | -91,6667 | 7,85863 | 213,4551 | -11,6645 | 2,1E-13 |

| | | | | | |
|---|---|---|---|---|---|
| day1 h.aka sal15 temp22 - day2 h.aka sal15 temp22 | -46,25 | 7,664356 | 180 | -6,03443 | 2,12E-05 |
| day1 h.aka sal15 temp22 - day3 h.aka sal15 temp22 | -81,6667 | 7,664356 | 180 | -10,6554 | 4,49E-13 |
| day1 h.aka sal15 temp22 - day4 h.aka sal15 temp22 | -91,6667 | 7,664356 | 180 | -11,9601 | 0 |
| day1 h.aka sal15 temp22 - day5 h.aka sal15 temp22 | -85,4167 | 7,664356 | 180 | -11,1447 | 2,42E-13 |
| day1 h.aka sal15 temp22 - day3 h.aka sal30 temp22 | -60,4167 | 7,85863 | 213,4551 | -7,68794 | 1,37E-09 |
| day1 h.aka sal15 temp22 - day4 h.aka sal30 temp22 | -58,3333 | 7,85863 | 213,4551 | -7,42284 | 6,82E-09 |
| day1 h.aka sal15 temp22 - day5 h.aka sal30 temp22 | -83,3333 | 7,85863 | 213,4551 | -10,6041 | 6,96E-13 |
| day2 h.aka sal15 temp22 - day3 h.aka sal15 temp22 | -35,4167 | 7,664356 | 180 | -4,62096 | 0,012611 |
| day2 h.aka sal15 temp22 - day4 h.aka sal15 temp22 | -45,4167 | 7,664356 | 180 | -5,9257 | 3,66E-05 |
| day2 h.aka sal15 temp22 - day5 h.aka sal15 temp22 | -39,1667 | 7,664356 | 180 | -5,11024 | 0,001681 |
| day2 h.aka sal15 temp22 - day0 h.rot sal15 temp22 | 54,58333 | 7,85863 | 213,4551 | 6,945655 | 1,13E-07 |
| day2 h.aka sal15 temp22 - day1 h.rot sal15 temp22 | 54,58333 | 7,85863 | 213,4551 | 6,945655 | 1,13E-07 |
| day2 h.aka sal15 temp22 - day2 h.rot sal15 temp22 | 47,8869 | 7,85863 | 213,4551 | 6,093544 | 1,23E-05 |
| day2 h.aka sal15 temp22 - day3 h.rot sal15 temp22 | 54,58333 | 7,85863 | 213,4551 | 6,945655 | 1,13E-07 |
| day2 h.aka sal15 temp22 - day4 h.rot sal15 temp22 | 54,58333 | 7,85863 | 213,4551 | 6,945655 | 1,13E-07 |
| day2 h.aka sal15 temp22 - day5 h.rot sal15 temp22 | 44,7619 | 7,85863 | 213,4551 | 5,695892 | 9,4E-05 |
| day2 h.aka sal15 temp22 - day0 both sal30 temp22 | 54,58333 | 7,85863 | 213,4551 | 6,945655 | 1,13E-07 |
| day2 h.aka sal15 temp22 - day1 both sal30 temp22 | 54,58333 | 7,85863 | 213,4551 | 6,945655 | 1,13E-07 |
| day2 h.aka sal15 temp22 - day2 both sal30 temp22 | 54,58333 | 7,85863 | 213,4551 | 6,945655 | 1,13E-07 |
| day2 h.aka sal15 temp22 - day3 both sal30 temp22 | 48,95833 | 7,85863 | 213,4551 | 6,229882 | 5,99E-06 |
| day2 h.aka sal15 temp22 - day4 both sal30 temp22 | 54,58333 | 7,85863 | 213,4551 | 6,945655 | 1,13E-07 |
| day2 h.aka sal15 temp22 - day5 both sal30 temp22 | 46,25 | 7,85863 | 213,4551 | 5,88525 | 3,62E-05 |
| day2 h.aka sal15 temp22 - day0 h.aka sal30 temp22 | 54,58333 | 7,85863 | 213,4551 | 6,945655 | 1,13E-07 |
| day2 h.aka sal15 temp22 - day1 h.aka sal30 temp22 | 54,58333 | 7,85863 | 213,4551 | 6,945655 | 1,13E-07 |
| day2 h.aka sal15 temp22 - day2 h.aka sal30 temp22 | 33,33333 | 7,85863 | 213,4551 | 4,241622 | 0,047643 |
| day2 h.aka sal15 temp22 - day5 h.aka sal30 temp22 | -37,0833 | 7,85863 | 213,4551 | -4,7188 | 0,007932 |
| day2 h.aka sal15 temp22 - day0 h.rot sal30 temp22 | 54,58333 | 7,85863 | 213,4551 | 6,945655 | 1,13E-07 |
| day2 h.aka sal15 temp22 - day1 h.rot sal30 temp22 | 54,58333 | 7,85863 | 213,4551 | 6,945655 | 1,13E-07 |
| day2 h.aka sal15 temp22 - day2 h.rot sal30 temp22 | 50,41667 | 7,85863 | 213,4551 | 6,415453 | 2,2E-06 |
| day2 h.aka sal15 temp22 - day3 h.rot sal30 temp22 | 45,83333 | 7,85863 | 213,4551 | 5,83223 | 4,74E-05 |
| day2 h.aka sal15 temp22 - day4 h.rot sal30 temp22 | 54,58333 | 7,85863 | 213,4551 | 6,945655 | 1,13E-07 |
| day2 h.aka sal15 temp22 - day5 h.rot sal30 temp22 | 45,41667 | 7,85863 | 213,4551 | 5,779209 | 6,2E-05 |
| day3 h.aka sal15 temp22 - day0 h.rot sal15 temp22 | 90 | 7,85863 | 213,4551 | 11,45238 | 2,87E-13 |
| day3 h.aka sal15 temp22 - day1 h.rot sal15 temp22 | 90 | 7,85863 | 213,4551 | 11,45238 | 2,87E-13 |
| day3 h.aka sal15 temp22 - day2 h.rot sal15 temp22 | 83,30357 | 7,85863 | 213,4551 | 10,60027 | 6,99E-13 |
| day3 h.aka sal15 temp22 - day3 h.rot sal15 temp22 | 90 | 7,85863 | 213,4551 | 11,45238 | 2,87E-13 |
| day3 h.aka sal15 temp22 - day4 h.rot sal15 temp22 | 90 | 7,85863 | 213,4551 | 11,45238 | 2,87E-13 |
| day3 h.aka sal15 temp22 - day5 h.rot sal15 temp22 | 80,17857 | 7,85863 | 213,4551 | 10,20261 | 7,59E-13 |
| day3 h.aka sal15 temp22 - day0 both sal30 temp22 | 90 | 7,85863 | 213,4551 | 11,45238 | 2,87E-13 |
| day3 h.aka sal15 temp22 - day1 both sal30 temp22 | 90 | 7,85863 | 213,4551 | 11,45238 | 2,87E-13 |
| day3 h.aka sal15 temp22 - day2 both sal30 temp22 | 90 | 7,85863 | 213,4551 | 11,45238 | 2,87E-13 |
| day3 h.aka sal15 temp22 - day3 both sal30 temp22 | 84,375 | 7,85863 | 213,4551 | 10,7366 | 5,96E-13 |
| day3 h.aka sal15 temp22 - day4 both sal30 temp22 | 90 | 7,85863 | 213,4551 | 11,45238 | 2,87E-13 |
| day3 h.aka sal15 temp22 - day5 both sal30 temp22 | 81,66667 | 7,85863 | 213,4551 | 10,39197 | 7,11E-13 |
| day3 h.aka sal15 temp22 - day0 h.aka sal30 temp22 | 90 | 7,85863 | 213,4551 | 11,45238 | 2,87E-13 |
| day3 h.aka sal15 temp22 - day1 h.aka sal30 temp22 | 90 | 7,85863 | 213,4551 | 11,45238 | 2,87E-13 |
| day3 h.aka sal15 temp22 - day2 h.aka sal30 temp22 | 68,75 | 7,85863 | 213,4551 | 8,748345 | 2,44E-12 |
| day3 h.aka sal15 temp22 - day0 h.rot sal30 temp22 | 90 | 7,85863 | 213,4551 | 11,45238 | 2,87E-13 |
| day3 h.aka sal15 temp22 - day1 h.rot sal30 temp22 | 90 | 7,85863 | 213,4551 | 11,45238 | 2,87E-13 |
| day3 h.aka sal15 temp22 - day2 h.rot sal30 temp22 | 85,83333 | 7,85863 | 213,4551 | 10,92218 | 5,24E-13 |
| day3 h.aka sal15 temp22 - day3 h.rot sal30 temp22 | 81,25 | 7,85863 | 213,4551 | 10,33895 | 7,3E-13 |
| day3 h.aka sal15 temp22 - day4 h.rot sal30 temp22 | 90 | 7,85863 | 213,4551 | 11,45238 | 2,87E-13 |
| day3 h.aka sal15 temp22 - day5 h.rot sal30 temp22 | 80,83333 | 7,85863 | 213,4551 | 10,28593 | 7,39E-13 |
| day4 h.aka sal15 temp22 - day0 h.rot sal15 temp22 | 100 | 7,85863 | 213,4551 | 12,72486 | 9,56E-14 |
| day4 h.aka sal15 temp22 - day1 h.rot sal15 temp22 | 100 | 7,85863 | 213,4551 | 12,72486 | 9,56E-14 |
| day4 h.aka sal15 temp22 - day2 h.rot sal15 temp22 | 93,30357 | 7,85863 | 213,4551 | 11,87275 | 1,62E-13 |
| day4 h.aka sal15 temp22 - day3 h.rot sal15 temp22 | 100 | 7,85863 | 213,4551 | 12,72486 | 9,56E-14 |
| day4 h.aka sal15 temp22 - day4 h.rot sal15 temp22 | 100 | 7,85863 | 213,4551 | 12,72486 | 9,56E-14 |
| day4 h.aka sal15 temp22 - day5 h.rot sal15 temp22 | 90,17857 | 7,85863 | 213,4551 | 11,4751 | 2,86E-13 |
| day4 h.aka sal15 temp22 - day0 both sal30 temp22 | 100 | 7,85863 | 213,4551 | 12,72486 | 9,56E-14 |
| day4 h.aka sal15 temp22 - day1 both sal30 temp22 | 100 | 7,85863 | 213,4551 | 12,72486 | 9,56E-14 |
| day4 h.aka sal15 temp22 - day2 both sal30 temp22 | 100 | 7,85863 | 213,4551 | 12,72486 | 9,56E-14 |
| day4 h.aka sal15 temp22 - day3 both sal30 temp22 | 94,375 | 7,85863 | 213,4551 | 12,00909 | 1,37E-13 |
| day4 h.aka sal15 temp22 - day4 both sal30 temp22 | 100 | 7,85863 | 213,4551 | 12,72486 | 9,56E-14 |
| day4 h.aka sal15 temp22 - day5 both sal30 temp22 | 91,66667 | 7,85863 | 213,4551 | 11,66446 | 2,1E-13 |
| day4 h.aka sal15 temp22 - day0 h.aka sal30 temp22 | 100 | 7,85863 | 213,4551 | 12,72486 | 9,56E-14 |
| day4 h.aka sal15 temp22 - day1 h.aka sal30 temp22 | 100 | 7,85863 | 213,4551 | 12,72486 | 9,56E-14 |
| day4 h.aka sal15 temp22 - day2 h.aka sal30 temp22 | 78,75 | 7,85863 | 213,4551 | 10,02083 | 7,76E-13 |
| day4 h.aka sal15 temp22 - day4 h.aka sal30 temp22 | 33,33333 | 7,85863 | 213,4551 | 4,241622 | 0,047643 |
| day4 h.aka sal15 temp22 - day0 h.rot sal30 temp22 | 100 | 7,85863 | 213,4551 | 12,72486 | 9,56E-14 |
| day4 h.aka sal15 temp22 - day1 h.rot sal30 temp22 | 100 | 7,85863 | 213,4551 | 12,72486 | 9,56E-14 |
| day4 h.aka sal15 temp22 - day2 h.rot sal30 temp22 | 95,83333 | 7,85863 | 213,4551 | 12,19466 | 1,15E-13 |
| day4 h.aka sal15 temp22 - day3 h.rot sal30 temp22 | 91,25 | 7,85863 | 213,4551 | 11,61144 | 2,25E-13 |
| day4 h.aka sal15 temp22 - day4 h.rot sal30 temp22 | 100 | 7,85863 | 213,4551 | 12,72486 | 9,56E-14 |
| day4 h.aka sal15 temp22 - day5 h.rot sal30 temp22 | 90,83333 | 7,85863 | 213,4551 | 11,55842 | 2,44E-13 |
| day5 h.aka sal15 temp22 - day0 h.rot sal15 temp22 | 93,75 | 7,85863 | 213,4551 | 11,92956 | 1,43E-13 |
| day5 h.aka sal15 temp22 - day1 h.rot sal15 temp22 | 93,75 | 7,85863 | 213,4551 | 11,92956 | 1,43E-13 |
| day5 h.aka sal15 temp22 - day2 h.rot sal15 temp22 | 87,05357 | 7,85863 | 213,4551 | 11,07745 | 4,65E-13 |
| day5 h.aka sal15 temp22 - day3 h.rot sal15 temp22 | 93,75 | 7,85863 | 213,4551 | 11,92956 | 1,43E-13 |
| day5 h.aka sal15 temp22 - day4 h.rot sal15 temp22 | 93,75 | 7,85863 | 213,4551 | 11,92956 | 1,43E-13 |
| day5 h.aka sal15 temp22 - day5 h.rot sal15 temp22 | 83,92857 | 7,85863 | 213,4551 | 10,6798 | 6,32E-13 |
| day5 h.aka sal15 temp22 - day0 both sal30 temp22 | 93,75 | 7,85863 | 213,4551 | 11,92956 | 1,43E-13 |
| day5 h.aka sal15 temp22 - day1 both sal30 temp22 | 93,75 | 7,85863 | 213,4551 | 11,92956 | 1,43E-13 |
| day5 h.aka sal15 temp22 - day2 both sal30 temp22 | 93,75 | 7,85863 | 213,4551 | 11,92956 | 1,43E-13 |

| | | | | | |
|---|---|---|---|---|---|
| day5 h.aka sal15 temp22 - day3 both sal30 temp22 | 88,125 | 7,85863 | 213,4551 | 11,21379 | 3,92E-13 |
| day5 h.aka sal15 temp22 - day4 both sal30 temp22 | 93,75 | 7,85863 | 213,4551 | 11,92956 | 1,43E-13 |
| day5 h.aka sal15 temp22 - day5 both sal30 temp22 | 85,41667 | 7,85863 | 213,4551 | 10,86916 | 5,63E-13 |
| day5 h.aka sal15 temp22 - day0 h.aka sal30 temp22 | 93,75 | 7,85863 | 213,4551 | 11,92956 | 1,43E-13 |
| day5 h.aka sal15 temp22 - day1 h.aka sal30 temp22 | 93,75 | 7,85863 | 213,4551 | 11,92956 | 1,43E-13 |
| day5 h.aka sal15 temp22 - day2 h.aka sal30 temp22 | 72,5 | 7,85863 | 213,4551 | 9,225527 | 8,38E-13 |
| day5 h.aka sal15 temp22 - day0 h.rot sal30 temp22 | 93,75 | 7,85863 | 213,4551 | 11,92956 | 1,43E-13 |
| day5 h.aka sal15 temp22 - day1 h.rot sal30 temp22 | 93,75 | 7,85863 | 213,4551 | 11,92956 | 1,43E-13 |
| day5 h.aka sal15 temp22 - day2 h.rot sal30 temp22 | 89,58333 | 7,85863 | 213,4551 | 11,39936 | 2,97E-13 |
| day5 h.aka sal15 temp22 - day3 h.rot sal30 temp22 | 85 | 7,85863 | 213,4551 | 10,81614 | 5,98E-13 |
| day5 h.aka sal15 temp22 - day4 h.rot sal30 temp22 | 93,75 | 7,85863 | 213,4551 | 11,92956 | 1,43E-13 |
| day5 h.aka sal15 temp22 - day5 h.rot sal30 temp22 | 84,58333 | 7,85863 | 213,4551 | 10,76311 | 6E-13 |
| day0 h.rot sal15 temp22 - day3 h.aka sal30 temp22 | -68,75 | 7,85863 | 213,4551 | -8,74834 | 2,44E-12 |
| day0 h.rot sal15 temp22 - day4 h.aka sal30 temp22 | -66,6667 | 7,85863 | 213,4551 | -8,48324 | 1,01E-11 |
| day0 h.rot sal15 temp22 - day5 h.aka sal30 temp22 | -91,6667 | 7,85863 | 213,4551 | -11,6645 | 2,1E-13 |
| day1 h.rot sal15 temp22 - day3 h.aka sal30 temp22 | -68,75 | 7,85863 | 213,4551 | -8,74834 | 2,44E-12 |
| day1 h.rot sal15 temp22 - day4 h.aka sal30 temp22 | -66,6667 | 7,85863 | 213,4551 | -8,48324 | 1,01E-11 |
| day1 h.rot sal15 temp22 - day5 h.aka sal30 temp22 | -91,6667 | 7,85863 | 213,4551 | -11,6645 | 2,1E-13 |
| day2 h.rot sal15 temp22 - day3 h.aka sal30 temp22 | -62,0536 | 7,85863 | 213,4551 | -7,89623 | 3,81E-10 |
| day2 h.rot sal15 temp22 - day4 h.aka sal30 temp22 | -59,9702 | 7,85863 | 213,4551 | -7,63113 | 1,94E-09 |
| day2 h.rot sal15 temp22 - day5 h.aka sal30 temp22 | -84,9702 | 7,85863 | 213,4551 | -10,8123 | 5,98E-13 |
| day3 h.rot sal15 temp22 - day3 h.aka sal30 temp22 | -68,75 | 7,85863 | 213,4551 | -8,74834 | 2,44E-12 |
| day3 h.rot sal15 temp22 - day4 h.aka sal30 temp22 | -66,6667 | 7,85863 | 213,4551 | -8,48324 | 1,01E-11 |
| day3 h.rot sal15 temp22 - day5 h.aka sal30 temp22 | -91,6667 | 7,85863 | 213,4551 | -11,6645 | 2,1E-13 |
| day4 h.rot sal15 temp22 - day3 h.aka sal30 temp22 | -68,75 | 7,85863 | 213,4551 | -8,74834 | 2,44E-12 |
| day4 h.rot sal15 temp22 - day4 h.aka sal30 temp22 | -66,6667 | 7,85863 | 213,4551 | -8,48324 | 1,01E-11 |
| day4 h.rot sal15 temp22 - day5 h.aka sal30 temp22 | -91,6667 | 7,85863 | 213,4551 | -11,6645 | 2,1E-13 |
| day5 h.rot sal15 temp22 - day3 h.aka sal30 temp22 | -58,9286 | 7,85863 | 213,4551 | -7,49858 | 4,33E-09 |
| day5 h.rot sal15 temp22 - day4 h.aka sal30 temp22 | -56,8452 | 7,85863 | 213,4551 | -7,23348 | 2,1E-08 |
| day5 h.rot sal15 temp22 - day5 h.aka sal30 temp22 | -81,8452 | 7,85863 | 213,4551 | -10,4147 | 7,25E-13 |
| day0 both sal30 temp22 - day3 h.aka sal30 temp22 | -68,75 | 7,85863 | 213,4551 | -8,74834 | 2,44E-12 |
| day0 both sal30 temp22 - day4 h.aka sal30 temp22 | -66,6667 | 7,85863 | 213,4551 | -8,48324 | 1,01E-11 |
| day0 both sal30 temp22 - day5 h.aka sal30 temp22 | -91,6667 | 7,85863 | 213,4551 | -11,6645 | 2,1E-13 |
| day1 both sal30 temp22 - day3 h.aka sal30 temp22 | -68,75 | 7,85863 | 213,4551 | -8,74834 | 2,44E-12 |
| day1 both sal30 temp22 - day4 h.aka sal30 temp22 | -66,6667 | 7,85863 | 213,4551 | -8,48324 | 1,01E-11 |
| day1 both sal30 temp22 - day5 h.aka sal30 temp22 | -91,6667 | 7,85863 | 213,4551 | -11,6645 | 2,1E-13 |
| day2 both sal30 temp22 - day3 h.aka sal30 temp22 | -68,75 | 7,85863 | 213,4551 | -8,74834 | 2,44E-12 |
| day2 both sal30 temp22 - day4 h.aka sal30 temp22 | -66,6667 | 7,85863 | 213,4551 | -8,48324 | 1,01E-11 |
| day2 both sal30 temp22 - day5 h.aka sal30 temp22 | -91,6667 | 7,85863 | 213,4551 | -11,6645 | 2,1E-13 |
| day3 both sal30 temp22 - day3 h.aka sal30 temp22 | -63,125 | 7,85863 | 213,4551 | -8,03257 | 1,64E-10 |
| day3 both sal30 temp22 - day4 h.aka sal30 temp22 | -61,0417 | 7,85863 | 213,4551 | -7,76747 | 8,43E-10 |
| day3 both sal30 temp22 - day5 h.aka sal30 temp22 | -86,0417 | 7,85863 | 213,4551 | -10,9487 | 5,24E-13 |
| day4 both sal30 temp22 - day3 h.aka sal30 temp22 | -68,75 | 7,85863 | 213,4551 | -8,74834 | 2,44E-12 |
| day4 both sal30 temp22 - day4 h.aka sal30 temp22 | -66,6667 | 7,85863 | 213,4551 | -8,48324 | 1,01E-11 |
| day4 both sal30 temp22 - day5 h.aka sal30 temp22 | -91,6667 | 7,85863 | 213,4551 | -11,6645 | 2,1E-13 |
| day5 both sal30 temp22 - day3 h.aka sal30 temp22 | -60,4167 | 7,85863 | 213,4551 | -7,68794 | 1,37E-09 |
| day5 both sal30 temp22 - day4 h.aka sal30 temp22 | -58,3333 | 7,85863 | 213,4551 | -7,42284 | 6,82E-09 |
| day5 both sal30 temp22 - day5 h.aka sal30 temp22 | -83,3333 | 7,85863 | 213,4551 | -10,6041 | 6,96E-13 |
| day0 h.aka sal30 temp22 - day3 h.aka sal30 temp22 | -68,75 | 7,664356 | 180 | -8,97009 | 1,55E-12 |
| day0 h.aka sal30 temp22 - day4 h.aka sal30 temp22 | -66,6667 | 7,664356 | 180 | -8,69827 | 5,9E-12 |
| day0 h.aka sal30 temp22 - day5 h.aka sal30 temp22 | -91,6667 | 7,664356 | 180 | -11,9601 | 0 |
| day1 h.aka sal30 temp22 - day3 h.aka sal30 temp22 | -68,75 | 7,664356 | 180 | -8,97009 | 1,55E-12 |
| day1 h.aka sal30 temp22 - day4 h.aka sal30 temp22 | -66,6667 | 7,664356 | 180 | -8,69827 | 5,9E-12 |
| day1 h.aka sal30 temp22 - day5 h.aka sal30 temp22 | -91,6667 | 7,664356 | 180 | -11,9601 | 0 |
| day2 h.aka sal30 temp22 - day3 h.aka sal30 temp22 | -47,5 | 7,664356 | 180 | -6,19752 | 9,21E-06 |
| day2 h.aka sal30 temp22 - day4 h.aka sal30 temp22 | -45,4167 | 7,664356 | 180 | -5,9257 | 3,66E-05 |
| day2 h.aka sal30 temp22 - day5 h.aka sal30 temp22 | -70,4167 | 7,664356 | 180 | -9,18755 | 8,06E-13 |
| day3 h.aka sal30 temp22 - day0 h.rot sal30 temp22 | 68,75 | 7,85863 | 213,4551 | 8,748345 | 2,44E-12 |
| day3 h.aka sal30 temp22 - day1 h.rot sal30 temp22 | 68,75 | 7,85863 | 213,4551 | 8,748345 | 2,44E-12 |
| day3 h.aka sal30 temp22 - day2 h.rot sal30 temp22 | 64,58333 | 7,85863 | 213,4551 | 8,218142 | 5,14E-11 |
| day3 h.aka sal30 temp22 - day3 h.rot sal30 temp22 | 60 | 7,85863 | 213,4551 | 7,634919 | 1,9E-09 |
| day3 h.aka sal30 temp22 - day4 h.rot sal30 temp22 | 68,75 | 7,85863 | 213,4551 | 8,748345 | 2,44E-12 |
| day3 h.aka sal30 temp22 - day5 h.rot sal30 temp22 | 59,58333 | 7,85863 | 213,4551 | 7,581899 | 2,62E-09 |
| day4 h.aka sal30 temp22 - day0 h.rot sal30 temp22 | 66,66667 | 7,85863 | 213,4551 | 8,483243 | 1,01E-11 |
| day4 h.aka sal30 temp22 - day1 h.rot sal30 temp22 | 66,66667 | 7,85863 | 213,4551 | 8,483243 | 1,01E-11 |
| day4 h.aka sal30 temp22 - day2 h.rot sal30 temp22 | 62,5 | 7,85863 | 213,4551 | 7,953041 | 2,68E-10 |
| day4 h.aka sal30 temp22 - day3 h.rot sal30 temp22 | 57,91667 | 7,85863 | 213,4551 | 7,369818 | 9,37E-09 |
| day4 h.aka sal30 temp22 - day4 h.rot sal30 temp22 | 66,66667 | 7,85863 | 213,4551 | 8,483243 | 1,01E-11 |
| day4 h.aka sal30 temp22 - day5 h.rot sal30 temp22 | 57,5 | 7,85863 | 213,4551 | 7,316797 | 1,28E-08 |
| day5 h.aka sal30 temp22 - day0 h.rot sal30 temp22 | 91,66667 | 7,85863 | 213,4551 | 11,66446 | 2,1E-13 |
| day5 h.aka sal30 temp22 - day1 h.rot sal30 temp22 | 91,66667 | 7,85863 | 213,4551 | 11,66446 | 2,1E-13 |
| day5 h.aka sal30 temp22 - day2 h.rot sal30 temp22 | 87,5 | 7,85863 | 213,4551 | 11,13426 | 4,35E-13 |
| day5 h.aka sal30 temp22 - day3 h.rot sal30 temp22 | 82,91667 | 7,85863 | 213,4551 | 10,55103 | 6,57E-13 |
| day5 h.aka sal30 temp22 - day4 h.rot sal30 temp22 | 91,66667 | 7,85863 | 213,4551 | 11,66446 | 2,1E-13 |
| day5 h.aka sal30 temp22 - day5 h.rot sal30 temp22 | 82,5 | 7,85863 | 213,4551 | 10,49801 | 7,08E-13 |

**Table S8:** Post-hoc results for all significant pairwise comparisons (response: *A. tonsa* nauplii relative population grazing rate in full-factorial grazing assay across three different prey composition treatments, two levels of salinity and two levels of temperature; pooled algae biovolumes in the polycultures)

```
emmeans(model1, pairwise ~ prey * sal * temp)
$emmeans

Degrees-of-freedom method: satterthwaite
P value adjustment: tukey method for comparing a family of 12 estimates
```

| contrast | estimate | SE | df | t.ratio | p.value |
|---|---|---|---|---|---|
| both sal15 temp16 - both sal15 temp22 | -0,17684 | 0,037662 | 11,25535 | -4,69555 | 0,017989 |
| both sal15 temp16 - h.rot sal15 temp22 | -0,33472 | 0,036678 | 9,921284 | -9,12569 | 0,00013 |
| both sal15 temp16 - h.rot sal30 temp22 | -0,18642 | 0,036678 | 18,39329 | -5,08253 | 0,00308 |
| h.aka sal15 temp16 - both sal15 temp22 | -0,25086 | 0,037559 | 3,40514 | -6,67916 | 0,04426 |
| h.aka sal15 temp16 - h.rot sal15 temp22 | -0,40874 | 0,036573 | 2,983112 | -11,1759 | 0,013801 |
| h.aka sal15 temp16 - both sal30 temp22 | -0,17148 | 0,037559 | 15,67843 | -4,56572 | 0,011981 |
| h.aka sal15 temp16 - h.rot sal30 temp22 | -0,26044 | 0,036573 | 6,210411 | -7,12108 | 0,006528 |
| h.rot sal15 temp16 - both sal30 temp22 | -0,1754 | 0,037379 | 10,86407 | -4,69237 | 0,019342 |
| h.rot sal15 temp16 - h.rot sal15 temp22 | -0,33327 | 0,036388 | 9,54217 | -9,15879 | 0,00016 |
| h.rot sal15 temp16 - h.rot sal30 temp22 | -0,18497 | 0,036388 | 17,98869 | -5,08333 | 0,003247 |
| both sal30 temp16 - h.rot sal15 temp22 | -0,2867 | 0,036678 | 3,026202 | -7,81645 | 0,036567 |
| h.aka sal30 temp16 - both sal15 temp22 | -0,25197 | 0,037559 | 7,822098 | -6,70868 | 0,004169 |
| h.aka sal30 temp16 - h.rot sal15 temp22 | -0,40984 | 0,036573 | 6,800966 | -11,2062 | 0,000291 |
| h.aka sal30 temp16 - both sal30 temp22 | -0,17259 | 0,037559 | 14,1499 | -4,59525 | 0,013725 |
| h.aka sal30 temp16 - h.rot sal30 temp22 | -0,26155 | 0,036573 | 17,8134 | -7,1514 | 5,98E-05 |
| h.rot sal30 temp16 - both sal15 temp22 | -0,2317 | 0,037379 | 7,628049 | -6,19868 | 0,00746 |
| h.rot sal30 temp16 - h.rot sal15 temp22 | -0,38957 | 0,036388 | 6,620595 | -10,7061 | 0,000457 |
| h.rot sal30 temp16 - both sal30 temp22 | -0,15232 | 0,037379 | 14,19856 | -4,07506 | 0,034041 |
| h.rot sal30 temp16 - h.rot sal30 temp22 | -0,24128 | 0,036388 | 17,40155 | -6,63066 | 0,000176 |
| both sal15 temp22 - h.aka sal15 temp22 | 0,187446 | 0,046601 | 28,47622 | 4,02238 | 0,016355 |
| both sal15 temp22 - h.aka sal30 temp22 | 0,203169 | 0,046601 | 13,96949 | 4,359781 | 0,021191 |
| h.aka sal15 temp22 - h.rot sal15 temp22 | -0,34532 | 0,04581 | 27,73204 | -7,53813 | 2,03E-06 |
| h.aka sal15 temp22 - h.rot sal30 temp22 | -0,19702 | 0,04581 | 19,43882 | -4,30089 | 0,01418 |
| h.rot sal15 temp22 - both sal30 temp22 | 0,237252 | 0,046099 | 28,02853 | 5,146534 | 0,000941 |
| h.rot sal15 temp22 - h.aka sal30 temp22 | 0,361043 | 0,04581 | 12,64559 | 7,881354 | 0,000126 |

**Table S9:** Post-hoc results for all significant main effects and interactive effects (response: *A. tonsa* nauplii relative population grazing rate in the polycultures with algae biovolumes separated by algae species)

```
  Tukey multiple comparisons of means
    95% family-wise confidence level

Fit: aov(formula = grazing_rate ~ species*temperature*salinity, data =
polycultures)

Tukey_grazing_polycultures$species[Tukey_rmax_H.aka$species[,4]<0.05,]
                                diff          lwr         upr        p adj
h.rot - h.aka              0.1157561  0.0626244500  0.16888776 1.494167e-04

Tukey_grazing_polycultures$temperature[Tukey_rmax_H.aka$temperature[,4]<0.0
5,]
                                diff          lwr         upr        p adj
22 - 15                    0.1400765  0.0869448467  0.19320815 1.365769e-05

Tukey_grazing_polycultures$species:temperature[Tukey_rmax_H.aka$species:tem
perature[,4]<0.05,]
                                diff          lwr         upr        p adj
h.rot:22 - h.aka:16        0.2558326  0.1554010615  0.35626414 1.656686e-06
h.rot:22 - h.rot:16        0.2307129  0.1302814030  0.33114448 8.499416e-06
h.rot:22 - h.aka:22        0.2063925  0.1059610063  0.30682409 4.337993e-05

Tukey_grazing_polycultures$species:salinity[Tukey_rmax_H.aka$species:salini
ty[,4]<0.05,]
                                diff          lwr         upr        p adj
h.rot:15 - h.aka:15        0.1078497  0.0074181596  0.20828124 3.209528e-02
h.aka:30 - h.rot:15       -0.1563635 -0.2567950015 -0.05593192 1.335341e-03
h.rot:30 - h.aka:30        0.1236625  0.0232309647  0.22409405 1.187852e-02

Tukey_grazing_polycultures$temperature:salinity[Tukey_rmax_H.aka$species:sa
linity[,4]<0.05,]
                                diff          lwr         upr        p adj
22:15 - 15:15              0.2323675  0.1319359205  0.33279900 7.619136e-06
15:30 - 22:15             -0.1806839 -0.2811153982 -0.08025232 2.514563e-04
22:30 - 22:15             -0.1328983 -0.2333298606 -0.03246678 6.493373e-03

Tukey_grazing_polycultures$species:temperature:salinity[Tukey_rmax_H.aka$sp
ecies:temperature:salinity[,4]<0.05,]
                                diff          lwr         upr        p adj
h.rot:22:15 - h.aka:16:15  0.3402172  0.1696977195  0.51073660 1.906650e-05
h.rot:22:30 - h.aka:16:15  0.2338989  0.0633794687  0.40441835 2.848633e-03
h.rot:22:15 - h.rot:16:15  0.3043302  0.1338107967  0.47484968 1.009340e-04
h.rot:22:30 - h.rot:16:15  0.1980120  0.0274925460  0.36853143 1.504009e-02
h.rot:22:15 - h.rot:22:15  0.1798125  0.0092930358  0.35033192 3.365917e-02
h.aka:16:30 - h.rot:22:15 -0.2777663 -0.4482857352 -0.10724685 3.544467e-04
h.rot:16:30 - h.rot:22:15 -0.2634139 -0.4339333410 -0.09289446 7.015478e-04
h.aka:22:30 - h.rot:22:15 -0.3392909 -0.5098103085 -0.16877143 1.989328e-05
h.rot:22:30 - h.aka:15:30  0.1714480  0.0009286012  0.34196748 4.808839e-02
h.rot:22:30 - h.aka:22:30  0.2329726  0.0624531745  0.40349206 2.975915e-03
```

**Table S10:** Post-hoc results for all significant pairwise comparisons (response: *B. plicatilis* mortality)

```
emmeans(rotifer_mortality, pairwise ~ day * prey * sal)
$emmeans

Degrees-of-freedom method: kenward-roger
P value adjustment: tukey method for comparing a family of 18 estimates
```

| contrast | estimate | SE | df | t.ratio | p.value |
|---|---|---|---|---|---|
| sal15 both day0 - sal30 h.aka day6 | -64,6825 | 9,642739 | 33,43804 | -6,7079 | 1,49E-05 |
| sal30 both day0 - sal30 h.aka day6 | -64,6825 | 9,642739 | 33,43804 | -6,7079 | 1,49E-05 |
| sal15 h.aka day0 - sal30 h.aka day6 | -64,6825 | 9,642739 | 33,43804 | -6,7079 | 1,49E-05 |
| sal30 h.aka day0 - sal30 h.aka day6 | -64,6825 | 8,64773 | 24 | -7,47971 | 1,21E-05 |
| sal15 h.rot day0 - sal30 h.aka day6 | -64,6825 | 9,642739 | 33,43804 | -6,7079 | 1,49E-05 |
| sal30 h.rot day0 - sal30 h.aka day6 | -64,6825 | 9,642739 | 33,43804 | -6,7079 | 1,49E-05 |
| sal15 both day3 - sal30 h.aka day6 | -54,3651 | 9,642739 | 33,43804 | -5,63793 | 0,000323 |
| sal30 both day3 - sal30 h.aka day6 | -59,9206 | 9,642739 | 33,43804 | -6,21407 | 6,16E-05 |
| sal15 h.aka day3 - sal30 h.aka day6 | -58,0159 | 9,642739 | 33,43804 | -6,01653 | 0,000109 |
| sal30 h.aka day3 - sal30 h.aka day6 | -42,2288 | 8,64773 | 24 | -4,88323 | 0,005198 |
| sal15 h.rot day3 - sal30 h.aka day6 | -51,9553 | 9,642739 | 33,43804 | -5,38802 | 0,000659 |
| sal30 h.rot day3 - sal30 h.aka day6 | -64,6825 | 9,642739 | 33,43804 | -6,7079 | 1,49E-05 |
| sal15 both day6 - sal30 h.aka day6 | -57,7381 | 9,642739 | 33,43804 | -5,98773 | 0,000118 |
| sal30 both day6 - sal30 h.aka day6 | -54,7849 | 9,642739 | 33,43804 | -5,68146 | 0,000285 |
| sal15 h.aka day6 - sal30 h.aka day6 | -44,4444 | 9,642739 | 33,43804 | -4,60911 | 0,005795 |
| sal30 h.aka day6 - sal15 h.rot day6 | 58,93541 | 9,642739 | 33,43804 | 6,111896 | 8,26E-05 |

**Table S11:** Post-hoc results for all significant main effects and interactions (response: *B. plicatilis* relative population grazing rate based on pooled algae biovolumes in the polycultures)

```
  Tukey multiple comparisons of means
    95% family-wise confidence level

Fit: aov(formula = grazing_rate ~ salinity*prey, data = rotifer_grazing)

Tukey_ rotifer_grazing$salinity[Tukey_rotifer_grazing$salinity [,4]<0.05,]
                                  diff        lwr         upr        p adj
30-15                      -0.08682374 -0.1182524 -0.05539510 6.037188e-05

Tukey_rotifer_grazing$salinity:prey[Tukey_rotifer_grazing$salinity:prey
[,4]<0.05,]
                                  diff        lwr         upr        p adj
30:h.rot-15:both           -0.11868298 -0.2026031 -0.03476284 4.855431e-03
30:h.rot-30:both           -0.10033778 -0.1842579 -0.01641765 1.650307e-02
30:h.rot-15:h.rot          -0.16040108 -0.2443212 -0.07648095 3.671096e-04
30:h.aka-15:h.rot          -0.10591979 -0.1898399 -0.02199966 1.133247e-02
15:h.aka-30:h.rot           0.13620624  0.0522861  0.22012638 1.578154e-03
```

**Table S12:** Post-hoc results for all significant main effects and interactive effects (response: *B. plicatilis* relative population grazing rate in the polycultures with algae biovolumes separated by algae species)

```
  Tukey multiple comparisons of means
    95% family-wise confidence level

Fit: aov(formula = grazing_rate ~ salinity*species, data =
rotifer_polycultures)

Tukey_rotifer_polycultures$salinity[Tukey_rotifer_grazing$salinity
[,4]<0.05,]
                                  diff        lwr           upr       p adj
30-15                      -0.07048231 -0.1108405 -3.012408e-02 0.001618397

Tukey_rotifer_polycultures$salinity:species[Tukey_rotifer_grazing$salinity:
species [,4]<0.05,]
                                  diff        lwr           upr       p adj
h.rot:30-h.aka:15          -0.09101071 -0.1675939 -1.442752e-02 0.016327674
h.aka:30-h.aka:15          -0.07660810 -0.1531913 -2.490458e-05 0.049907096
```

**Table S13:** ANOVA results testing the effects of salinity and prey composition and consumer identity on the on mortality (left) and population grazing rates (right). Degrees of freedom (Df), F and p-values are given for each effect. Values marked with an asterisk (*) indicate significant effects ($p < 0.05$).

| | Mortality (GLS) | | | Population Grazing Rate (AOV) | | |
|---|---|---|---|---|---|---|
| Effect | Df | *F* | *p* | Df | *F* | *p* |
| Salinity | 1 | 0.4 | 0.510 | 1 | 14.2 | <0.001 * |
| Prey Composition | 2 | 35.9 | <0.001 * | 2 | 12.6 | <0.001 * |
| Consumer | 1 | 0.3 | 0.600 | 1 | 23.6 | <0.001 * |
| Consumer × Prey Composition | 2 | 0.9 | 0.400 | 2 | 5.9 | 0.007 * |
| Consumer × Salinity | 1 | 8.2 | 0.008 * | 1 | 16.0 | <0.001 * |
| Prey Composition × Salinity | 2 | 0.5 | 0.626 | 2 | 11.3 | <0.001 * |
| Consumer × Prey Composition × Salinity | 2 | 4.0 | 0.030 * | 2 | 0.3 | 0.760 |

**Table S14:** Post-hoc results for all significant pairwise comparisons (response: Mortalities of both grazer species at 16°C across different prey combinations and two salinity levels (15 PSU and 30 PSU) over time.

```
emmeans(grazer_mortality, pairwise ~ day * prey * sal * grazer)
$emmeans

Degrees-of-freedom method: kenward-roger
P value adjustment: tukey method for comparing a family of 84 estimates
```

| contrast | estimate | SE | df | t.ratio | p.value |
|---|---|---|---|---|---|
| both sal15 day0 A.tonsa - h.aka sal15 day4 A.tonsa | -31,7453 | 7,264575 | 142,6408 | -4,36988 | 0,044412 |
| both sal15 day0 A.tonsa - h.aka sal15 day5 A.tonsa | -65,0786 | 7,264575 | 142,6408 | -8,95835 | 6,4E-12 |
| both sal15 day0 A.tonsa - h.aka sal30 day5 A.tonsa | -41,8643 | 7,264575 | 142,6408 | -5,76281 | 0,000149 |
| both sal15 day0 A.tonsa - h.aka sal30 day6 B.plicatilis | -64,5797 | 7,817794 | 141,8142 | -8,2606 | 3,19E-10 |
| h.aka sal15 day0 A.tonsa - h.aka sal15 day4 A.tonsa | -31,5476 | 7,210963 | 141,0426 | -4,37495 | 0,043893 |
| h.aka sal15 day0 A.tonsa - h.aka sal15 day5 A.tonsa | -64,881 | 7,210963 | 141,0426 | -8,99754 | 5,64E-12 |
| h.aka sal15 day0 A.tonsa - h.aka sal30 day5 A.tonsa | -41,6667 | 7,210963 | 141,0426 | -5,77824 | 0,000142 |
| h.aka sal15 day0 A.tonsa - h.aka sal30 day6 B.plicatilis | -64,382 | 7,806136 | 141,6077 | -8,24762 | 3,43E-10 |
| h.rot sal15 day0 A.tonsa - h.aka sal15 day4 A.tonsa | -31,5476 | 7,210963 | 141,0426 | -4,37495 | 0,043893 |
| h.rot sal15 day0 A.tonsa - h.aka sal15 day5 A.tonsa | -64,881 | 7,210963 | 141,0426 | -8,99754 | 5,64E-12 |
| h.rot sal15 day0 A.tonsa - h.aka sal30 day5 A.tonsa | -41,6667 | 7,210963 | 141,0426 | -5,77824 | 0,000142 |
| h.rot sal15 day0 A.tonsa - h.aka sal30 day6 B.plicatilis | -64,382 | 7,806136 | 141,6077 | -8,24762 | 3,43E-10 |
| both sal30 day0 A.tonsa - h.aka sal15 day5 A.tonsa | -64,6833 | 7,264575 | 142,6408 | -8,90393 | 8,49E-12 |
| both sal30 day0 A.tonsa - h.aka sal30 day5 A.tonsa | -41,469 | 7,264575 | 142,6408 | -5,70838 | 0,000192 |
| both sal30 day0 A.tonsa - h.aka sal30 day6 B.plicatilis | -64,1843 | 7,8934 | 143,5583 | -8,13139 | 6,17E-10 |
| h.aka sal30 day0 A.tonsa - h.aka sal15 day4 A.tonsa | -31,5476 | 7,210963 | 141,0426 | -4,37495 | 0,043893 |
| h.aka sal30 day0 A.tonsa - h.aka sal15 day5 A.tonsa | -64,881 | 7,210963 | 141,0426 | -8,99754 | 5,64E-12 |
| h.aka sal30 day0 A.tonsa - h.aka sal30 day5 A.tonsa | -41,6667 | 7,210963 | 141,0426 | -5,77824 | 0,000142 |
| h.aka sal30 day0 A.tonsa - h.aka sal30 day6 B.plicatilis | -64,382 | 7,806136 | 141,6077 | -8,24762 | 3,43E-10 |
| h.rot sal30 day0 A.tonsa - h.aka sal15 day4 A.tonsa | -31,5476 | 7,210963 | 141,0426 | -4,37495 | 0,043893 |
| h.rot sal30 day0 A.tonsa - h.aka sal15 day5 A.tonsa | -64,881 | 7,210963 | 141,0426 | -8,99754 | 5,64E-12 |
| h.rot sal30 day0 A.tonsa - h.aka sal30 day5 A.tonsa | -41,6667 | 7,210963 | 141,0426 | -5,77824 | 0,000142 |
| h.rot sal30 day0 A.tonsa - h.aka sal30 day6 B.plicatilis | -64,382 | 7,806136 | 141,6077 | -8,24762 | 3,43E-10 |
| both sal15 day1 A.tonsa - h.aka sal15 day4 A.tonsa | -31,7453 | 7,264575 | 142,6408 | -4,36988 | 0,044412 |
| both sal15 day1 A.tonsa - h.aka sal15 day5 A.tonsa | -65,0786 | 7,264575 | 142,6408 | -8,95835 | 6,4E-12 |
| both sal15 day1 A.tonsa - h.aka sal30 day5 A.tonsa | -41,8643 | 7,264575 | 142,6408 | -5,76281 | 0,000149 |
| both sal15 day1 A.tonsa - h.aka sal30 day6 B.plicatilis | -64,5797 | 7,817794 | 141,8142 | -8,2606 | 3,19E-10 |
| h.aka sal15 day1 A.tonsa - h.aka sal15 day4 A.tonsa | -31,5476 | 7,210963 | 141,0426 | -4,37495 | 0,043893 |
| h.aka sal15 day1 A.tonsa - h.aka sal15 day5 A.tonsa | -64,881 | 7,210963 | 141,0426 | -8,99754 | 5,64E-12 |
| h.aka sal15 day1 A.tonsa - h.aka sal30 day5 A.tonsa | -41,6667 | 7,210963 | 141,0426 | -5,77824 | 0,000142 |
| h.aka sal15 day1 A.tonsa - h.aka sal30 day6 B.plicatilis | -64,382 | 7,806136 | 141,6077 | -8,24762 | 3,43E-10 |
| h.rot sal15 day1 A.tonsa - h.aka sal15 day5 A.tonsa | -56,5476 | 7,210963 | 141,0426 | -7,8419 | 3,41E-09 |
| h.rot sal15 day1 A.tonsa - h.aka sal30 day5 A.tonsa | -33,3333 | 7,210963 | 141,0426 | -4,62259 | 0,018369 |
| h.rot sal15 day1 A.tonsa - h.aka sal30 day6 B.plicatilis | -56,0487 | 7,806136 | 141,6077 | -7,18008 | 1,22E-07 |
| both sal30 day1 A.tonsa - h.aka sal15 day5 A.tonsa | -64,6833 | 7,264575 | 142,6408 | -8,90393 | 8,49E-12 |
| both sal30 day1 A.tonsa - h.aka sal30 day5 A.tonsa | -41,469 | 7,264575 | 142,6408 | -5,70838 | 0,000192 |
| both sal30 day1 A.tonsa - h.aka sal30 day6 B.plicatilis | -64,1843 | 7,8934 | 143,5583 | -8,13139 | 6,17E-10 |
| h.aka sal30 day1 A.tonsa - h.aka sal15 day4 A.tonsa | -31,5476 | 7,210963 | 141,0426 | -4,37495 | 0,043893 |
| h.aka sal30 day1 A.tonsa - h.aka sal15 day5 A.tonsa | -64,881 | 7,210963 | 141,0426 | -8,99754 | 5,64E-12 |
| h.aka sal30 day1 A.tonsa - h.aka sal30 day5 A.tonsa | -41,6667 | 7,210963 | 141,0426 | -5,77824 | 0,000142 |
| h.aka sal30 day1 A.tonsa - h.aka sal30 day6 B.plicatilis | -64,382 | 7,806136 | 141,6077 | -8,24762 | 3,43E-10 |
| h.rot sal30 day1 A.tonsa - h.aka sal15 day4 A.tonsa | -31,5476 | 7,210963 | 141,0426 | -4,37495 | 0,043893 |
| h.rot sal30 day1 A.tonsa - h.aka sal15 day5 A.tonsa | -64,881 | 7,210963 | 141,0426 | -8,99754 | 5,64E-12 |
| h.rot sal30 day1 A.tonsa - h.aka sal30 day5 A.tonsa | -41,6667 | 7,210963 | 141,0426 | -5,77824 | 0,000142 |
| h.rot sal30 day1 A.tonsa - h.aka sal30 day6 B.plicatilis | -64,382 | 7,806136 | 141,6077 | -8,24762 | 3,43E-10 |
| both sal15 day2 A.tonsa - h.aka sal15 day4 A.tonsa | -31,7453 | 7,264575 | 142,6408 | -4,36988 | 0,044412 |
| both sal15 day2 A.tonsa - h.aka sal15 day5 A.tonsa | -65,0786 | 7,264575 | 142,6408 | -8,95835 | 6,4E-12 |
| both sal15 day2 A.tonsa - h.aka sal30 day5 A.tonsa | -41,8643 | 7,264575 | 142,6408 | -5,76281 | 0,000149 |
| both sal15 day2 A.tonsa - h.aka sal30 day6 B.plicatilis | -64,5797 | 7,817794 | 141,8142 | -8,2606 | 3,19E-10 |
| h.aka sal15 day2 A.tonsa - h.aka sal15 day4 A.tonsa | -31,5476 | 7,210963 | 141,0426 | -4,37495 | 0,043893 |
| h.aka sal15 day2 A.tonsa - h.aka sal15 day5 A.tonsa | -64,881 | 7,210963 | 141,0426 | -8,99754 | 5,64E-12 |
| h.aka sal15 day2 A.tonsa - h.aka sal30 day5 A.tonsa | -41,6667 | 7,210963 | 141,0426 | -5,77824 | 0,000142 |
| h.aka sal15 day2 A.tonsa - h.aka sal30 day6 B.plicatilis | -64,382 | 7,806136 | 141,6077 | -8,24762 | 3,43E-10 |
| h.rot sal15 day2 A.tonsa - h.aka sal15 day4 A.tonsa | -31,5476 | 7,210963 | 141,0426 | -4,37495 | 0,043893 |
| h.rot sal15 day2 A.tonsa - h.aka sal15 day5 A.tonsa | -64,881 | 7,210963 | 141,0426 | -8,99754 | 5,64E-12 |
| h.rot sal15 day2 A.tonsa - h.aka sal30 day5 A.tonsa | -41,6667 | 7,210963 | 141,0426 | -5,77824 | 0,000142 |
| h.rot sal15 day2 A.tonsa - h.aka sal30 day6 B.plicatilis | -64,382 | 7,806136 | 141,6077 | -8,24762 | 3,43E-10 |
| both sal30 day2 A.tonsa - h.aka sal15 day5 A.tonsa | -64,6833 | 7,264575 | 142,6408 | -8,90393 | 8,49E-12 |
| both sal30 day2 A.tonsa - h.aka sal30 day5 A.tonsa | -41,469 | 7,264575 | 142,6408 | -5,70838 | 0,000192 |
| both sal30 day2 A.tonsa - h.aka sal30 day6 B.plicatilis | -64,1843 | 7,8934 | 143,5583 | -8,13139 | 6,17E-10 |
| h.aka sal30 day2 A.tonsa - h.aka sal15 day4 A.tonsa | -31,5476 | 7,210963 | 141,0426 | -4,37495 | 0,043893 |
| h.aka sal30 day2 A.tonsa - h.aka sal15 day5 A.tonsa | -64,881 | 7,210963 | 141,0426 | -8,99754 | 5,64E-12 |
| h.aka sal30 day2 A.tonsa - h.aka sal30 day5 A.tonsa | -41,6667 | 7,210963 | 141,0426 | -5,77824 | 0,000142 |
| h.aka sal30 day2 A.tonsa - h.aka sal30 day6 B.plicatilis | -64,382 | 7,806136 | 141,6077 | -8,24762 | 3,43E-10 |
| h.rot sal30 day2 A.tonsa - h.aka sal15 day4 A.tonsa | -31,5476 | 7,210963 | 141,0426 | -4,37495 | 0,043893 |
| h.rot sal30 day2 A.tonsa - h.aka sal15 day5 A.tonsa | -64,881 | 7,210963 | 141,0426 | -8,99754 | 5,64E-12 |
| h.rot sal30 day2 A.tonsa - h.aka sal30 day5 A.tonsa | -41,6667 | 7,210963 | 141,0426 | -5,77824 | 0,000142 |
| h.rot sal30 day2 A.tonsa - h.aka sal30 day6 B.plicatilis | -64,382 | 7,806136 | 141,6077 | -8,24762 | 3,43E-10 |
| both sal15 day3 A.tonsa - h.aka sal15 day5 A.tonsa | -56,7453 | 7,264575 | 142,6408 | -7,81123 | 3,81E-09 |
| both sal15 day3 A.tonsa - h.aka sal30 day5 A.tonsa | -33,531 | 7,264575 | 142,6408 | -4,61569 | 0,018701 |
| both sal15 day3 A.tonsa - h.aka sal30 day6 B.plicatilis | -56,2464 | 7,817794 | 141,8142 | -7,19466 | 1,12E-07 |

| | | | | | |
|---|---|---|---|---|---|
| h.aka sal15 day3 A.tonsa - h.aka sal15 day5 A.tonsa | -41,5476 | 7,210963 | 141,0426 | -5,76173 | 0,000153 |
| h.aka sal15 day3 A.tonsa - h.aka sal30 day6 B.plicatilis | -41,0487 | 7,806136 | 141,6077 | -5,25851 | 0,001441 |
| h.rot sal15 day3 A.tonsa - h.aka sal15 day4 A.tonsa | -31,5476 | 7,210963 | 141,0426 | -4,37495 | 0,043893 |
| h.rot sal15 day3 A.tonsa - h.aka sal15 day5 A.tonsa | -64,881 | 7,210963 | 141,0426 | -8,99754 | 5,64E-12 |
| h.rot sal15 day3 A.tonsa - h.aka sal30 day5 A.tonsa | -41,6667 | 7,210963 | 141,0426 | -5,77824 | 0,000142 |
| h.rot sal15 day3 A.tonsa - h.aka sal30 day6 B.plicatilis | -64,382 | 7,806136 | 141,6077 | -8,24762 | 3,43E-10 |
| both sal30 day3 A.tonsa - h.aka sal15 day5 A.tonsa | -64,6833 | 7,264575 | 142,6408 | -8,90393 | 8,49E-12 |
| both sal30 day3 A.tonsa - h.aka sal30 day5 A.tonsa | -41,469 | 7,264575 | 142,6408 | -5,70838 | 0,000192 |
| both sal30 day3 A.tonsa - h.aka sal30 day6 B.plicatilis | -64,1843 | 7,8934 | 143,5583 | -8,13139 | 6,17E-10 |
| h.aka sal30 day3 A.tonsa - h.aka sal15 day5 A.tonsa | -53,4226 | 7,210963 | 141,0426 | -7,40853 | 3,65E-08 |
| h.aka sal30 day3 A.tonsa - h.aka sal30 day6 B.plicatilis | -52,9237 | 7,806136 | 141,6077 | -6,77975 | 9,98E-07 |
| h.rot sal30 day3 A.tonsa - h.aka sal15 day4 A.tonsa | -31,5476 | 7,210963 | 141,0426 | -4,37495 | 0,043893 |
| h.rot sal30 day3 A.tonsa - h.aka sal15 day5 A.tonsa | -64,881 | 7,210963 | 141,0426 | -8,99754 | 5,64E-12 |
| h.rot sal30 day3 A.tonsa - h.aka sal30 day5 A.tonsa | -41,6667 | 7,210963 | 141,0426 | -5,77824 | 0,000142 |
| h.rot sal30 day3 A.tonsa - h.aka sal30 day6 B.plicatilis | -64,382 | 7,806136 | 141,6077 | -8,24762 | 3,43E-10 |
| both sal15 day4 A.tonsa - h.aka sal15 day5 A.tonsa | -60,0786 | 7,264575 | 142,6408 | -8,27008 | 2,92E-10 |
| both sal15 day4 A.tonsa - h.aka sal30 day5 A.tonsa | -36,8643 | 7,264575 | 142,6408 | -5,07454 | 0,003107 |
| both sal15 day4 A.tonsa - h.aka sal30 day6 B.plicatilis | -59,5797 | 7,817794 | 141,8142 | -7,62104 | 1,12E-08 |
| h.aka sal15 day4 A.tonsa - h.rot sal30 day4 A.tonsa | 31,54762 | 7,210963 | 141,0426 | 4,374952 | 0,043893 |
| h.aka sal15 day4 A.tonsa - h.aka sal15 day5 A.tonsa | -33,3333 | 7,210963 | 141,0426 | -4,62259 | 0,018369 |
| h.aka sal15 day4 A.tonsa - h.rot sal30 day5 A.tonsa | 31,54762 | 7,210963 | 141,0426 | 4,374952 | 0,043893 |
| h.rot sal15 day4 A.tonsa - h.aka sal15 day5 A.tonsa | -56,5476 | 7,210963 | 141,0426 | -7,8419 | 3,41E-09 |
| h.rot sal15 day4 A.tonsa - h.aka sal30 day5 A.tonsa | -33,3333 | 7,210963 | 141,0426 | -4,62259 | 0,018369 |
| h.rot sal15 day4 A.tonsa - h.aka sal30 day6 B.plicatilis | -56,0487 | 7,806136 | 141,6077 | -7,18008 | 1,22E-07 |
| both sal30 day4 A.tonsa - h.aka sal15 day5 A.tonsa | -56,3499 | 7,264575 | 142,6408 | -7,75681 | 5,16E-09 |
| both sal30 day4 A.tonsa - h.aka sal30 day5 A.tonsa | -33,1357 | 7,264575 | 142,6408 | -4,56126 | 0,022794 |
| both sal30 day4 A.tonsa - h.aka sal30 day6 B.plicatilis | -55,851 | 7,8934 | 143,5583 | -7,07566 | 2,03E-07 |
| h.aka sal30 day4 A.tonsa - h.aka sal15 day5 A.tonsa | -61,3095 | 7,210963 | 141,0426 | -8,50227 | 8,46E-11 |
| h.aka sal30 day4 A.tonsa - h.aka sal30 day5 A.tonsa | -38,0952 | 7,210963 | 141,0426 | -5,28296 | 0,001306 |
| h.aka sal30 day4 A.tonsa - h.aka sal30 day6 B.plicatilis | -60,8106 | 7,806136 | 141,6077 | -7,7901 | 4,41E-09 |
| h.rot sal30 day4 A.tonsa - h.aka sal15 day5 A.tonsa | -64,881 | 7,210963 | 141,0426 | -8,99754 | 5,64E-12 |
| h.rot sal30 day4 A.tonsa - h.aka sal30 day5 A.tonsa | -41,6667 | 7,210963 | 141,0426 | -5,77824 | 0,000142 |
| h.rot sal30 day4 A.tonsa - h.aka sal30 day6 B.plicatilis | -64,382 | 7,806136 | 141,6077 | -8,24762 | 3,43E-10 |
| both sal15 day5 A.tonsa - h.aka sal15 day5 A.tonsa | -56,7453 | 7,264575 | 142,6408 | -7,81123 | 3,81E-09 |
| both sal15 day5 A.tonsa - h.aka sal30 day5 A.tonsa | -33,531 | 7,264575 | 142,6408 | -4,61569 | 0,018701 |
| both sal15 day5 A.tonsa - h.aka sal30 day6 B.plicatilis | -56,2464 | 7,817794 | 141,8142 | -7,19466 | 1,12E-07 |
| h.aka sal15 day5 A.tonsa - h.rot sal15 day5 A.tonsa | 55,71429 | 7,210963 | 141,0426 | 7,726331 | 6,46E-09 |
| h.aka sal15 day5 A.tonsa - both sal30 day5 A.tonsa | 58,43327 | 7,264575 | 142,6408 | 8,043591 | 1,04E-09 |
| h.aka sal15 day5 A.tonsa - h.rot sal30 day5 A.tonsa | 64,88095 | 7,210963 | 141,0426 | 8,997543 | 5,64E-12 |
| h.aka sal15 day5 A.tonsa - both sal15 day0 B.plicatilis | 65,18148 | 7,806136 | 141,6077 | 8,350031 | 1,92E-10 |
| h.aka sal15 day5 A.tonsa - h.aka sal15 day0 B.plicatilis | 65,18148 | 7,806136 | 141,6077 | 8,350031 | 1,92E-10 |
| h.aka sal15 day5 A.tonsa - h.rot sal15 day0 B.plicatilis | 65,18148 | 7,806136 | 141,6077 | 8,350031 | 1,92E-10 |
| h.aka sal15 day5 A.tonsa - both sal30 day0 B.plicatilis | 65,18148 | 7,806136 | 141,6077 | 8,350031 | 1,92E-10 |
| h.aka sal15 day5 A.tonsa - h.aka sal30 day0 B.plicatilis | 65,18148 | 7,806136 | 141,6077 | 8,350031 | 1,92E-10 |
| h.aka sal15 day5 A.tonsa - h.rot sal30 day0 B.plicatilis | 65,18148 | 7,806136 | 141,6077 | 8,350031 | 1,92E-10 |
| h.aka sal15 day5 A.tonsa - both sal15 day3 B.plicatilis | 54,86402 | 7,806136 | 141,6077 | 7,02832 | 2,72E-07 |
| h.aka sal15 day5 A.tonsa - h.aka sal15 day3 B.plicatilis | 58,51481 | 7,806136 | 141,6077 | 7,496002 | 2,22E-08 |
| h.aka sal15 day5 A.tonsa - h.rot sal15 day3 B.plicatilis | 52,45421 | 7,806136 | 141,6077 | 6,719612 | 1,36E-06 |
| h.aka sal15 day5 A.tonsa - both sal30 day3 B.plicatilis | 60,41958 | 7,806136 | 141,6077 | 7,74001 | 5,82E-09 |
| h.aka sal15 day5 A.tonsa - h.aka sal30 day3 B.plicatilis | 42,72778 | 7,806136 | 141,6077 | 5,473614 | 0,000563 |
| h.aka sal15 day5 A.tonsa - h.rot sal30 day3 B.plicatilis | 65,18148 | 7,806136 | 141,6077 | 8,350031 | 1,92E-10 |
| h.aka sal15 day5 A.tonsa - both sal15 day6 B.plicatilis | 58,23704 | 7,806136 | 141,6077 | 7,460417 | 2,69E-08 |
| h.aka sal15 day5 A.tonsa - h.aka sal15 day6 B.plicatilis | 44,94339 | 7,806136 | 141,6077 | 5,757443 | 0,000155 |
| h.aka sal15 day5 A.tonsa - h.rot sal15 day6 B.plicatilis | 59,43435 | 7,806136 | 141,6077 | 7,613799 | 1,16E-08 |
| h.aka sal15 day5 A.tonsa - both sal30 day6 B.plicatilis | 55,28382 | 7,806136 | 141,6077 | 7,082098 | 2,05E-07 |
| h.aka sal15 day5 A.tonsa - h.rot sal30 day6 B.plicatilis | 55,28405 | 7,806136 | 141,6077 | 7,082126 | 2,05E-07 |
| h.rot sal15 day5 A.tonsa - h.aka sal30 day5 A.tonsa | -32,5 | 7,210963 | 141,0426 | -4,50703 | 0,027844 |
| h.rot sal15 day5 A.tonsa - h.aka sal30 day6 B.plicatilis | -55,2153 | 7,806136 | 141,6077 | -7,07333 | 2,14E-07 |
| both sal30 day5 A.tonsa - h.aka sal30 day5 A.tonsa | -35,219 | 7,264575 | 142,6408 | -4,84804 | 0,00774 |
| both sal30 day5 A.tonsa - h.aka sal30 day6 B.plicatilis | -57,9343 | 7,8934 | 143,5583 | -7,33959 | 4,93E-08 |
| h.aka sal30 day5 A.tonsa - h.rot sal30 day5 A.tonsa | 41,66667 | 7,210963 | 141,0426 | 5,778239 | 0,000142 |
| h.aka sal30 day5 A.tonsa - both sal15 day0 B.plicatilis | 41,9672 | 7,806136 | 141,6077 | 5,37618 | 0,000865 |
| h.aka sal30 day5 A.tonsa - h.aka sal15 day0 B.plicatilis | 41,9672 | 7,806136 | 141,6077 | 5,37618 | 0,000865 |
| h.aka sal30 day5 A.tonsa - h.rot sal15 day0 B.plicatilis | 41,9672 | 7,806136 | 141,6077 | 5,37618 | 0,000865 |
| h.aka sal30 day5 A.tonsa - both sal30 day0 B.plicatilis | 41,9672 | 7,806136 | 141,6077 | 5,37618 | 0,000865 |
| h.aka sal30 day5 A.tonsa - h.aka sal30 day0 B.plicatilis | 41,9672 | 7,806136 | 141,6077 | 5,37618 | 0,000865 |
| h.aka sal30 day5 A.tonsa - h.rot sal30 day0 B.plicatilis | 41,9672 | 7,806136 | 141,6077 | 5,37618 | 0,000865 |
| h.aka sal30 day5 A.tonsa - h.aka sal15 day3 B.plicatilis | 35,30053 | 7,806136 | 141,6077 | 4,522151 | 0,026306 |
| h.aka sal30 day5 A.tonsa - both sal30 day3 B.plicatilis | 37,20529 | 7,806136 | 141,6077 | 4,766159 | 0,010679 |
| h.aka sal30 day5 A.tonsa - h.rot sal30 day3 B.plicatilis | 41,9672 | 7,806136 | 141,6077 | 5,37618 | 0,000865 |
| h.aka sal30 day5 A.tonsa - both sal15 day6 B.plicatilis | 35,02275 | 7,806136 | 141,6077 | 4,486567 | 0,029828 |
| h.aka sal30 day5 A.tonsa - h.rot sal15 day6 B.plicatilis | 36,22007 | 7,806136 | 141,6077 | 4,639948 | 0,017168 |
| h.rot sal30 day5 A.tonsa - h.aka sal30 day6 B.plicatilis | -64,382 | 7,806136 | 141,6077 | -8,24762 | 3,43E-10 |
| both sal15 day0 B.plicatilis - h.aka sal30 day6 B.plicatilis | -64,6825 | 8,326503 | 141,0426 | -7,76827 | 5,13E-09 |
| h.aka sal15 day0 B.plicatilis - h.aka sal30 day6 B.plicatilis | -64,6825 | 8,326503 | 141,0426 | -7,76827 | 5,13E-09 |
| h.rot sal15 day0 B.plicatilis - h.aka sal30 day6 B.plicatilis | -64,6825 | 8,326503 | 141,0426 | -7,76827 | 5,13E-09 |
| both sal30 day0 B.plicatilis - h.aka sal30 day6 B.plicatilis | -64,6825 | 8,326503 | 141,0426 | -7,76827 | 5,13E-09 |
| h.aka sal30 day0 B.plicatilis - h.aka sal30 day6 B.plicatilis | -64,6825 | 8,326503 | 141,0426 | -7,76827 | 5,13E-09 |
| h.rot sal30 day0 B.plicatilis - h.aka sal30 day6 B.plicatilis | -64,6825 | 8,326503 | 141,0426 | -7,76827 | 5,13E-09 |
| both sal15 day3 B.plicatilis - h.aka sal30 day6 B.plicatilis | -54,3651 | 8,326503 | 141,0426 | -6,52916 | 3,66E-06 |
| h.aka sal15 day3 B.plicatilis - h.aka sal30 day6 B.plicatilis | -58,0159 | 8,326503 | 141,0426 | -6,96762 | 3,82E-07 |
| h.rot sal15 day3 B.plicatilis - h.aka sal30 day6 B.plicatilis | -51,9553 | 8,326503 | 141,0426 | -6,23975 | 1,55E-05 |
| both sal30 day3 B.plicatilis - h.aka sal30 day6 B.plicatilis | -59,9206 | 8,326503 | 141,0426 | -7,19637 | 1,14E-07 |
| h.aka sal30 day3 B.plicatilis - h.aka sal30 day6 B.plicatilis | -42,2288 | 8,326503 | 141,0426 | -5,07162 | 0,003182 |

| | | | | | |
|---|---|---|---|---|---|
| h.rot sal30 day3 B.plicatilis - h.aka sal30 day6 B.plicatilis | -64,6825 | 8,326503 | 141,0426 | -7,76827 | 5,13E-09 |
| both sal15 day6 B.plicatilis - h.aka sal30 day6 B.plicatilis | -57,7381 | 8,326503 | 141,0426 | -6,93426 | 4,55E-07 |
| h.aka sal15 day6 B.plicatilis - h.aka sal30 day6 B.plicatilis | -44,4444 | 8,326503 | 141,0426 | -5,33771 | 0,001031 |
| h.rot sal15 day6 B.plicatilis - h.aka sal30 day6 B.plicatilis | -58,9354 | 8,326503 | 141,0426 | -7,07805 | 2,14E-07 |
| both sal30 day6 B.plicatilis - h.aka sal30 day6 B.plicatilis | -54,7849 | 8,326503 | 141,0426 | -6,57958 | 2,84E-06 |
| h.aka sal30 day6 B.plicatilis - h.rot sal30 day6 B.plicatilis | 54,7851 | 8,326503 | 141,0426 | 6,579606 | 2,84E-06 |

**Table S15:** Post-hoc results for all significant main effects and interactive effects (response: Relative population grazing rates of both grazer species at 16°C across different prey combinations and two salinity levels (15 PSU and 30 PSU).

```
  Tukey multiple comparisons of means
    95% family-wise confidence level

Fit: aov(formula = grazing_rates ~ salinity*grazer*prey, data =
grazing_both_grazers)

Tukey_both_grazers$grazer[Tukey_both_grazers$grazer [,4]<0.05,]
                                  diff         lwr          upr        p adj
B.plic-A.tonsa             -0.05084189 -0.07219394 -0.029489847 3.430216e-05

Tukey_both_grazers$prey[Tukey_both_grazers$prey [,4]<0.05,]
                                  diff         lwr          upr        p adj
h.aka-both                 -0.06240525 -0.09364868 -0.031161814 8.367168e-05
h.rot-both                 -0.04154128 -0.07278471 -0.010297843 7.252390e-03

Tukey_both_grazers$salinity[Tukey_both_grazers$salinity [,4]<0.05,]
                                  diff         lwr          upr        p adj
30-15                      -0.03899909 -0.06013213 -0.017866040 7.177760e-04

Tukey_both_grazers$grazer:prey[Tukey_both_grazers$grazer:prey [,4]<0.05,]
                                  diff         lwr          upr        p adj
B.plic:both-A.tonsa:both   -0.08611610 -0.14119532 -0.031036885 6.089784e-04
A.tonsa:h.aka-A.tonsa:both -0.09858422 -0.14957767 -0.047590777 2.696041e-05
B.plic:h.aka-A.tonsa:both  -0.10028272 -0.15536193 -0.045203502 6.973572e-05
B.plic:h.rot-A.tonsa:both  -0.11542594 -0.17050516 -0.060346725 6.903155e-06
B.plic:h.rot-A.tonsa:h.rot -0.06471109 -0.11979030 -0.009631869 1.399428e-02

Tukey_both_grazers$grazer:salinity[Tukey_both_grazers$grazer:salinity
[,4]<0.05,]
                                  diff         lwr          upr        p adj
B.plic:30-A.tonsa:15       -0.09581906 -0.13602290 -0.055615224 2.127065e-06
B.plic:30-B.plic:15        -0.08682374 -0.12980346 -0.043844033 3.304359e-05
B.plic:30-A.tonsa:30       -0.09268847 -0.13289231 -0.052484631 3.811448e-06

Tukey_both_grazers$prey:salinity[Tukey_both_grazers$prey:salinity
[,4]<0.05,]
                                  diff         lwr          upr        p adj
h.aka:30-both:15           -0.07044533 -0.12495962 -0.015931046 5.610784e-03
h.rot:30-both:15           -0.08221063 -0.13672492 -0.027696346 9.667109e-04
h.aka:30-h.rot:15          -0.08915179 -0.14366608 -0.034637506 3.332228e-04
h.rot:30-h.rot:15          -0.10091709 -0.15543138 -0.046402805 5.389051e-05
h.aka:30-both:30           -0.09002371 -0.14453800 -0.035509428 2.912625e-04
h.rot:30-both:30           -0.10178901 -0.15630330 -0.047274728 4.708031e-05

Tukey_both_grazers$grazer:prey:salinity[Tukey_both_grazers$grazer:prey:sali
nity [,4]<0.05,]
                                  diff         lwr          upr        p adj
B.plic:h.aka:30-           -0.11713466 -0.20769245 -0.026576869 3.721226e-03
A.tonsa:both:15

B.plic:h.rot:30-           -0.17161595 -0.26217374 -0.081058157 1.154955e-05
A.tonsa:both:15

A.tonsa:both:30-            0.10095404  0.01039624  0.191511826 1.889448e-02
B.plic:both:15

B.plic:h.rot:30-           -0.11868298 -0.21549333 -0.021872629 6.960348e-03
B.plic:both:15
```

| | | | | |
|---|---|---|---|---|
| A.tonsa:both:30-A.tonsa:h.aka:15 | 0.12204038 | 0.03820016 | 0.205880605 | 7.976243e-04 |
| B.plic:h.rot:30-A.tonsa:h.aka:15 | -0.09759663 | -0.18815442 | -0.007038843 | 2.606856e-02 |
| B.plic:h.rot:30-B.plic:h.aka:15 | -0.13620624 | -0.23301659 | -0.039395889 | 1.271743e-03 |
| B.plic:h.aka:30-A.tonsa:h.rot:15 | -0.11858239 | -0.20914018 | -0.028024597 | 3.203427e-03 |
| B.plic:h.rot:30-A.tonsa:h.rot:15 | -0.17306368 | -0.26362147 | -0.082505885 | 9.917934e-06 |
| B.plic:h.aka:30-B.plic:h.rot:15 | -0.10591979 | -0.20273014 | -0.009109444 | 2.263363e-02 |
| B.plic:h.rot:30-B.plic:h.rot:15 | -0.16040108 | -0.25721143 | -0.063590732 | 1.142289e-04 |
| B.plic:both:30-A.tonsa:both:30 | -0.11929923 | -0.20985702 | -0.028741443 | 2.973768e-03 |
| A.tonsa:h.aka:30-A.tonsa:both:30 | -0.12314913 | -0.20698935 | -0.039308907 | 7.024149e-04 |
| B.plic:h.aka:30-A.tonsa:both:30 | -0.16515573 | -0.25571352 | -0.074597937 | 2.285127e-05 |
| A.tonsa:h.rot:30-A.tonsa:both:30 | -0.10287744 | -0.18671766 | -0.019037215 | 6.888355e-03 |
| B.plic:h.rot:30-A.tonsa:both:30 | -0.21963702 | -0.31019481 | -0.129079225 | 8.732056e-08 |
| B.plic:h.rot:30-B.plic:both:30 | -0.10033778 | -0.19714813 | -0.003527431 | 3.700338e-02 |
| B.plic:h.rot:30-A.tonsa:h.aka:30 | -0.09648789 | -0.18704568 | -0.005930095 | 2.894917e-02 |
| B.plic:h.rot:30-A.tonsa:h.rot:30 | -0.11675958 | -0.20731737 | -0.026201786 | 3.868162e-03 |
